# High-resolution lineage tracking maps the multi-phase evolution and adaptive potential of biofilms

**DOI:** 10.64898/2026.09.04.749373

**Authors:** Alaksh Choudhury, Stanislas Thiriet-Rupert, Zoya Dixit, Melanie Magnan, Jean-Marc Ghigo, Christophe Beloin, Olivier Tenaillon

## Abstract

Although biofilms represent the dominant bacterial lifestyle and are major contributors to recalcitrant, treatment-resistant infections, their evolutionary dynamics at the individual lineage resolution remain poorly understood. We combine DNA barcoding-mediated lineage tracking and spatial sampling of > 100,000 lineages of *Escherichia coli* to characterize sequential phases of biofilm evolution quantitatively. Severe founder effects during surface colonization initially reduce the effective population size by nearly three orders of magnitude, establishing significant spatial segregation. This structure is subsequently eroded by a previously underappreciated secondary migration phase, during which cells released upon perturbation recolonize biofilms with a rate 10^3^-fold higher than naive planktonic cells. Adaptive evolution subsequently drives specialization through mutations targeting these distinct stages of the biofilm life cycle. Under severe antibiotic bottlenecks, active secondary migration continuously resuscitates local lineage diversity, enabling the parallel emergence of resistance across hundreds of independent lineages. Once locally established, migration drives a rapid spatial spread of resistant mutants. Together, these findings reveal how spatial segregation and secondary migration shape biofilm dynamics, coupling short-term ecological resilience with long-term adaptive potential.

## 1. Introduction

Evolutionary theory, largely based on studies using well-mixed liquid cultures, demonstrates that bacterial adaptation is a balance between mutation rate, genetic drift, and selection ^1–5^. However, most microbial lifeforms in nature and clinical settings exist in spatially structured habitats such as biofilms rather than homogenized cultures^6–9^. In biofilms, the fundamental assumptions of mixed cultures - such as global resource competition and uniform exposure to a common environment or stress - break down, thereby profoundly altering population and evolutionary dynamics. This distinct ecology is exemplified by the biofilm’s intrinsic antibiotic tolerance and accelerated emergence of resistance relative to well-mixed cultures, rendering biofilm-associated infections notoriously recalcitrant to treatment^10–17^.

Most biofilms are surface-attached communities that develop through dynamic cycles of attachment, growth, and dispersal^6^. During surface attachment, cells encounter pronounced population bottlenecks due to stochastic binding and clonal expansion of lineages ^18–20^. Rather than global competition for resources, spatially segregated lineages engage in intense local competition^21–23^. The population structure is further reshaped by passive detachment, dispersal, recolonization, and adaptive evolution ^7,24–27^. Biofilms establish complex internal chemical and nutrient gradients that generate non-uniform exposure to external stresses such as antibiotics ^28,29^. Consequently, evolutionary dynamics in biofilms are governed by repeated bottlenecks, local interactions, and heterogeneous nutrient and stress landscape, as opposed to the uniform selection pressures of well-mixed cultures.

Traditional approaches based on microscopy, population sequencing or bulk phenotypic measurements have provided important insights into biofilm architecture^30^. Yet, a precise lineage-level understanding of the surface colonisation, evolutionary dynamics and emergence of resistance remain elusive. For instance, the population bottlenecks upon colonisation remains largely uncharacterized ^16,22^. Furthermore, the relative contributions of initial colonization, dispersal, and selection—and the timescales over which each process dominates— remain unclear. Finally, how antibiotic exposure reshapes these lineage dynamics is also largely unknown.

Here, we address this gap using high-resolution lineage tracking based on genome barcoding ^31^ of the adherent-invasive *Escherichia coli* (AIEC) strain LF82, a well-established model for investigating biofilm-mediated persistence and inflammation in Crohn’s disease ^25,26^. We introduced randomized N20 barcodes at a neutral genomic locus of the LF82 strain using Cas9-mediated recombination, generating a library with over 100,000 uniquely tagged lineages or subpopulations (**Figure 1A**). We then used next-generation sequencing to track the lineage frequencies during biofilm formation and maturation. By combining spatial sampling with quantitative measures of diversity, we quantitatively disentangle the contributions of stochastic colonization, subsequent migration, and adaptive evolution towards biofilm population structure over time, and determine how intermittent antibiotic treatment influences these lineage-level dynamics.

**Figure 1.**
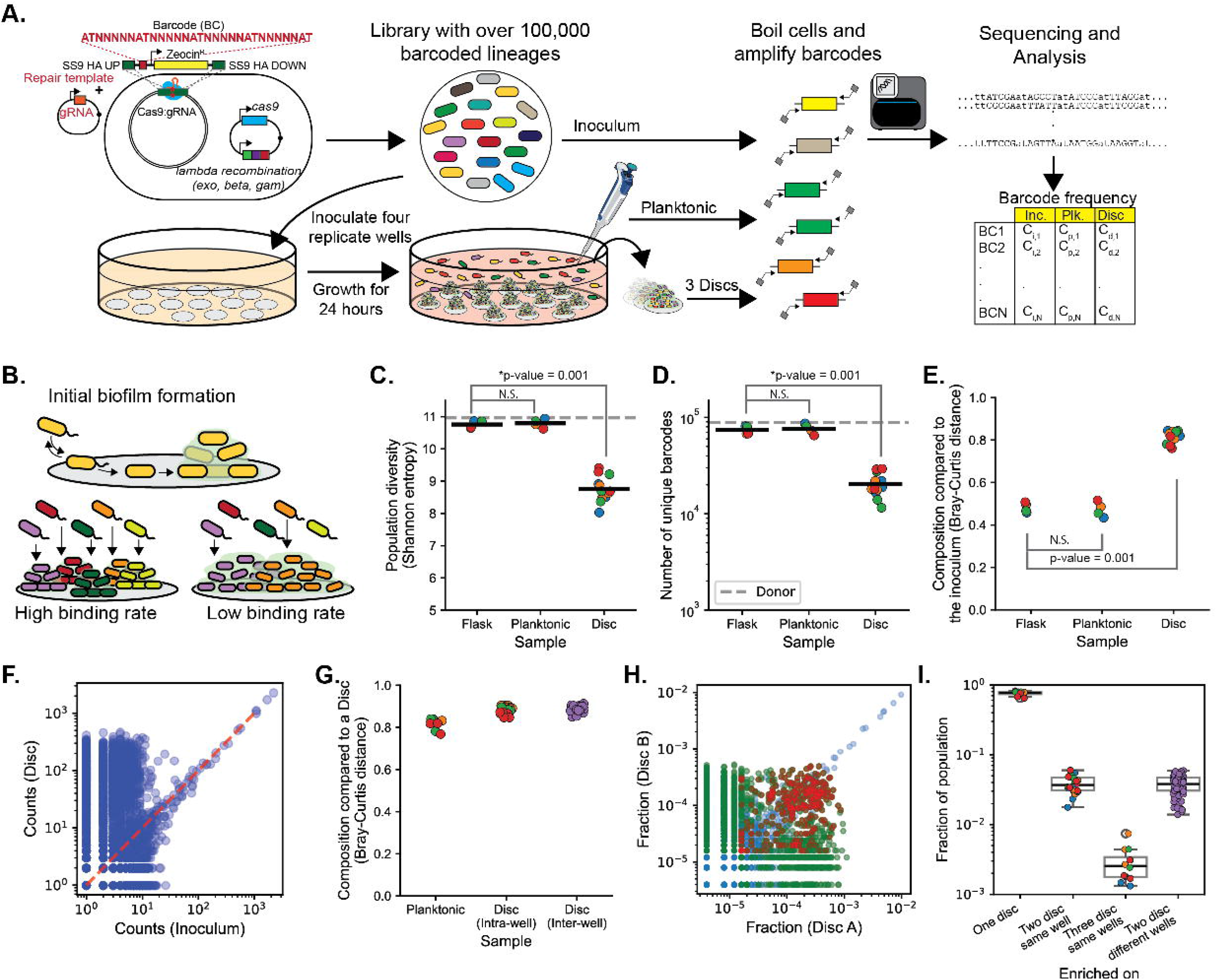
Initial binding dynamics impose bottlenecks that shape early biofilm population structure: **(A)** We used Cas9-mediated recombineering to introduce a randomized N20 barcode at a neutral locus (safe site 9) on the genome of *E. coli* strain LF82 to obtain a library with over 100,000 unique barcodes detected across experiments. We inoculated four replicate wells containing 18 small silicone discs each with the same inoculum of a library of barcoded cells. The cells attach to the discs and develop biofilms. Therefore, the discs allow us to sample a subsection of the surface-attached biofilm population, and we sample the planktonic population from the surrounding liquid media. We sequenced the barcodes from the inoculum, planktonic, and disc-bound populations to estimate the change in lineage frequencies in each population. **(B)** Conceptual schematic illustrating how surface-binding rates influence early biofilm population structure. High binding rates would promote uniform colonisation and preserve diversity, whereas low binding rates would impose strong stochastic bottlenecks that favor early colonizers and reduce diversity. **(C)** Population diversity measured as Shannon entropy (Σ *p*ₙ log(*p*ₙ), where *p*ₙ is the frequency of variant *n* for the inoculum (gray line), shake-flask (no-bottleneck and no-biofilm control), biofilm-associated planktonic, and disc-associated biofilm populations. **(D)** Number of unique barcodes recovered from the inoculum (gray line), shake-flask control, biofilm-associated planktonic and disc-associated biofilm populations after 24 h. **(E)** Compositional dissimilarity between the inoculum (reference) and the populations recovered from the shake flask control, biofilm-associated planktonic, and disc-associated biofilm populations, quantified using Bray– Curtis distance. **(F)** Comparison of barcode counts between the inoculum and a representative disc-associated biofilm from replicate A. Corresponding comparisons for all discs and replicate wells are shown in Supplementary Fig. 1. **(G)** Bray–Curtis distances comparing disc-associated populations to the biofilm-associated planktonic population, to other discs from the same well, and to discs from replicate wells. **(H)** Pairwise comparison of barcode counts between two discs recovered from the same well. Variants without any significant enrichment are blue, significantly enriched on one disc are green, and variants significantly enriched on both discs are red. Comparisons across replicate wells are shown in Supplementary Fig. 2. **(I)** Fraction of the disc-associated population represented by variants significantly enriched on one disc (green lineages from 1H and supplementary figure 2), on two discs from the same well (red lineages from 1H and supplementary figure 2), and on three discs from the same well, or on two discs from different replicate wells. As a no-bottleneck control, we inoculated four shake flasks with the same inoculum size at 4.3*10^7^ cells in each well. The inoculum size corresponds to ∼250-fold coverage of the ∼169,000 detected barcodes. All metrics were measured 24 h after inoculation from four independently inoculated wells (n = 4) and four shake-flask controls, with three discs sampled per well (total n = 12 discs). Circles represent individual measurements. Statistical significance was assessed using two-sided Kolmogorov–Smirnov tests; p-values are indicated (NS, not significant and significant when <0.01). For enrichment analyses (G, H), significance thresholds were determined using the distribution of barcode enrichment in the no-bottleneck populations. Variants were classified as significantly enriched if their enrichment exceeded the mean planktonic enrichment plus 1.96 standard deviations of the control (threshold = 0.623).

## 2. Results

### 2.1 Initial biofilm populations exhibit local bottlenecks and a segregated spatial structure

Upon initial colonization, only a stochastic subset of surface-attached cells commit to colony formation, and growth is localized to colony boundaries, which results in spatially structured and segregated populations ^6,9,18,20,23,24^. We however lack quantitative estimates of bottlenecks and effective population sizes (N_e_) associated with the stochastic binding.

We first quantified the surface colonization associated bottlenecks by combining spatial sampling of biofilm subpopulations^26^ with high-resolution lineage tracking. We inoculated four culture wells containing 18 small silicone discs with an *E. coli* library of over 100,000 uniquely-barcoded variants (**Figure 1A**). During a static incubation in the wells, cells grow in the planktonic phase while simultaneously attaching to the silicone discs, where they subsequently grow and develop into a biofilm. We sequenced the barcodes from the inoculum and three discs and the planktonic population from each replicate well after 24 hours of growth. Each disc with a population of ∼3*10^6^ CFUs, compared to ∼10^10^ CFUs in the planktonic phase, allowed us to track lineage dynamics in a discrete subsection of the total surface-attached population.

We reasoned that the rate of irreversible binding and subsequent growth on the surface jointly determine the final biofilm population structure (**Figure 1B**). High binding rates should increase the number of founders, yielding diverse biofilm communities. Conversely, low binding rates and rapid growth of early binders should decrease diversity via severe founder bottlenecks.

We quantified the population diversity using Shannon entropy, which accounts for both the number of lineages present and the evenness of their relative abundances. The Shannon entropy is high when many lineages are evenly distributed and low when a few lineages dominate. While Shannon entropy remained statistically indistinguishable between the inoculum, the no-biofilm no-bottleneck shake-flask controls, and the biofilm-associated planktonic populations, it was significantly lower within disc-associated biofilms (Komogrov-Smirnoff test, p-value = 10^-7^, **Figure 1C**). There were 20,410 ± 5,500 unique barcodes detected on the discs compared to the 76,388 ± 8,078 detected in the biofilm-associated planktonic phase (∼70% reduction; Komogrov-Smirnoff test, p = 10^-8^, **Figure 1D**). The observed strong loss of diversity confirmed the presence of an initial local bottleneck during surface colonization.

We then compared the composition between populations using the Bray-Curtis index, which ranges from 0 for population with identical lineage composition and relative abundances to 1 for completely distinct composition (**Figure 1E**). While the inoculum-flask control and inoculum-(biofilm-associated)-planktonic population distances were low (∼0.3), the inoculum-disc composition distance was high (∼0.8, **Figure 1E**). Many lineages on the discs had expanded disproportionately, with more than 100-fold increase in frequency relative to the inoculum (∼2.32 ± 0.18 log₁₀-fold; **Figure 1F** and **Supplementary Figures 1**). We estimated an effective population size (N_e_), which is the number of founding cells that contribute to the future generations, of 2332 +/- 570, which is three orders of magnitude lower than total biofilm population size (3.16*10^6^ +/- 1.3 * 10^6^). Therefore, only a few thousand founding lineages contribute to the final disc population with millions of cells.

Bray-Curtis dissimilarity between discs was also high (0.9) and was indistinguishable between discs recovered from the same well and those recovered from different replicate wells (**Figure 1G**). Within individual wells, the lineages enriched on a single disc constituted 75.6% of the population, whereas variants enriched on two discs comprised only 3.8% - a value again indistinguishable between discs from replicate wells (**Figure 1H-1I** and **Supplementary Figure 2)**. Therefore, initial stochastic attachment followed by a disproportionate expansion of a smaller subset of lineages compared to the inoculum, resulted in distinct population structures across discs.

### 2.2 Temporal differences in initial binding drive stochastic lineage expansion during colonization

We could not explain the expansion of rare lineages using a simple model where all cells experience a single bottleneck (**Supplementary Note 3**). Therefore, we developed a stochastic model with a Poisson approximation of binding (with rate *K_b_)* and logistic growth in the planktonic and surface-associated phases (with growth rates *µ_p_ and µ_d_*) (**Figure 2A**). We constrained the binding and growth by a maximum carrying capacities N_p,max_ and N_d,max_ in each phase. We could measure all model parameters experimentally (**Supplementary table 1** & **Supplementary Note 4**), except for the binding constant *K_b_*. To determine *K_b_*, we simulated the initial biofilm formation for a range of *K_b_* values (10^-9^ to 10^-5^ min^-1^). We showed, as hypothesized (**Figure 1B**), that the number of unique barcodes (and diversity) increased monotonically with the binding constant (**Figure 2B**).

**Figure 2:**
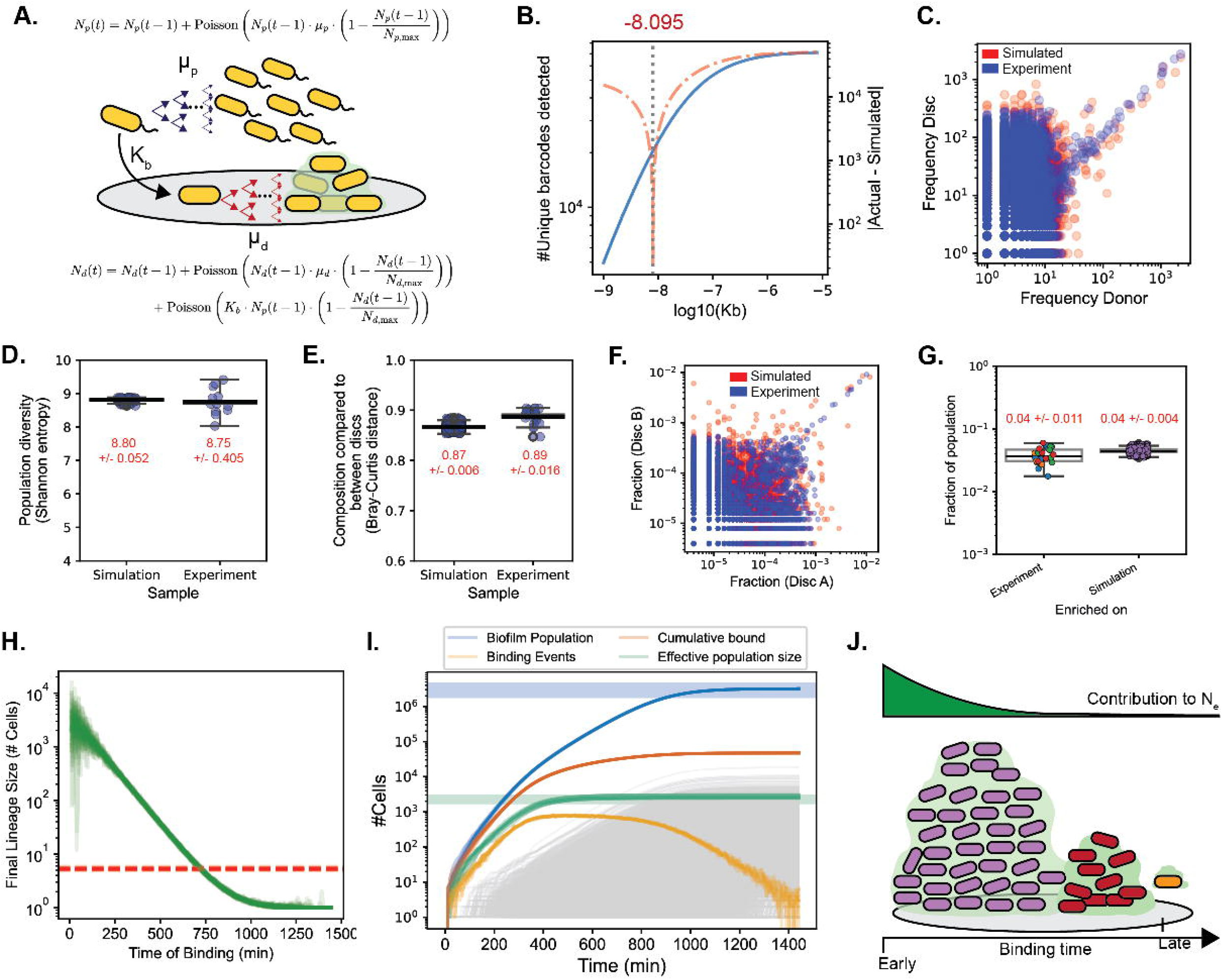
A binding–growth model recapitulates early biofilm colonisation dynamics. **(A)** Schematic of the stochastic binding–growth model used to simulate early biofilm formation, incorporating surface binding and population growth. Model is described in detail in the Materials and Methods section. **(B)** Model predictions across a range of binding constants (*K*_b_). Left axis (blue curve): number of unique barcodes recovered from simulations plotted against log₁₀(*K*_b_). Right axis: absolute difference between the mean numbers of unique barcodes recovered in simulations and the experimentally observed value. Ten independent simulations were performed for each *K*_b_. The optimum *K*_b_ value was defined as the value minimizing the difference between simulated and experimentally observed barcode diversity, represented with the gray vertical line. **(C)** Comparison of barcode counts between the inoculum and a representative experimental disc-associated biofilm from replicate A (blue), and between the inoculum and a representative disc from a simulation using the optimum *K*_b_ value (red). **(D)** Population diversity measured as Shannon entropy for experimental disc-associated populations and simulations at the optimum *K*_b_ value. **(E)** Bray–Curtis distances between disc-associated populations from simulations at the optimum *K*_b_ and experimental discs. **(F)** Pairwise comparison of barcode frequencies between two discs recovered from the same experimental well and between two discs recovered from simulations at the optimum *K*_b_ value. **(G)** Fraction of the disc-associated population represented by variants significantly enriched on two discs in the experiment and in the simulation, respectively. **(H)** The stochastic founder effect imposed a drift-based expansion of lineages represented as the final lineage size as a function of the time of binding for 10 simulations. The red line indicates the average lineage expansion at the end of the experiment. **(I)** The change in total biofilm population size (blue), the binding events (yellow), the cumulative binding events (red), the effective population size (green) over the first 24 hours of initial biofilm population for 10 simulations. The actual total population size (blue shaded area) and the actual effective population size (green shaded area) from 12 replicate discs at the end of the experiment. The gray lines represent the increase in individual lineages for one of the simulations. We calculated the effective population size (N_e_) as N_e_ = 1/p ^2^ where p_i_ is the relative abundance of barcode lineage i. **(J)** The effective population size Ne i.e., contribution of a lineage to the final population is a function of the binding time. Early binders have orders of magnitude higher success than late binders, making their contribution to the effective population size higher. Enrichment thresholds were defined as in Fig. 2. Box plots show the median (center line), interquartile range (box), and whiskers extending to 1.5× the interquartile range. In panels D, E, and G, we indicate the mean values for each sample with the standard deviation.

We determined an optimal *K_b_* of 10^−8.095^ by minimizing the difference between the actual and simulated number of unique barcodes (recovered from 10 simulations). At this optimal value, the simulated and the experimental communities showed similar expansion of the rare barcodes, Shannon entropy, disc-disc Bray-Curtis distance, and fraction of population composed of lineages significantly enriched on two discs (**Figure 2C-2G and Supplementary Figure 3-4).**

Our model revealed that differences in the surface attachment times generated significant biases in the contribution of individual lineages to the final population (**Figure 2H**). Cells binding at time = 0 are 57.4 times more successful than the average binder and three to four orders of magnitude more successful than the late arrivals (**Figure 2H**). The expansion of stochastically sampled early surface-binding lineages generated strong founder effects and local population bottlenecks, resulting in an increasing difference between the effective and the actual population size (green and blue curves, **Figure 2I**). The binding-growth simulation showed that early attachment conferred a strong lineage expansion advantage that shaped the spatially segregated lineage architecture in a sampled subsection of the biofilm population on a disc (**Figure 2J**).

### 2.3 Migration of variants erodes segregation and homogenizes the biofilm population

The life of the biofilm extends well beyond initial binding and macro-scale dispersion, recolonization, and adaptive evolution are well documented^7,16,32^. However, the underlying lineage-level temporal dynamics that reshape the population remain unresolved. Crucially the timescales over which different processes dominate are not known.

To characterize the temporal dynamics, we removed the spent media and replaced it with fresh media in 16 daily cycles, sampling the biofilm-associated planktonic population daily and in discs every 3 cycles (**Figure 3A**). Upon media replacement, the total CFUs per disc decreased from 5.4*10^6^ +/- 3.74*10^6^ to 5.4*10^5^ +/- 2.82*10^5^ and 1.3*10^7^ +/- 7.9*10^6^ released cells reseeded the planktonic phase (**Supplementary Figure 5**). Iterative sampling simulations across 1 to 12 discs confirmed that release from six discs captured total population diversity (p > 0.05), ensuring maintenance of population diversity through day 12 (**Supplementary Figure 6**). The biofilm-associated planktonic population diversity, number of unique barcodes detected, and composition remained close to the inoculum for several days (replicate A for 11 days; replicate B for 8 days; **Figure 3B-3C & Supplementary Figure 7A**). Thus, despite early spatial segregation across individual discs, the total biofilm population preserved diversity of the inoculum and reseeded a planktonic population with an identical lineage composition.

**Figure 3:**
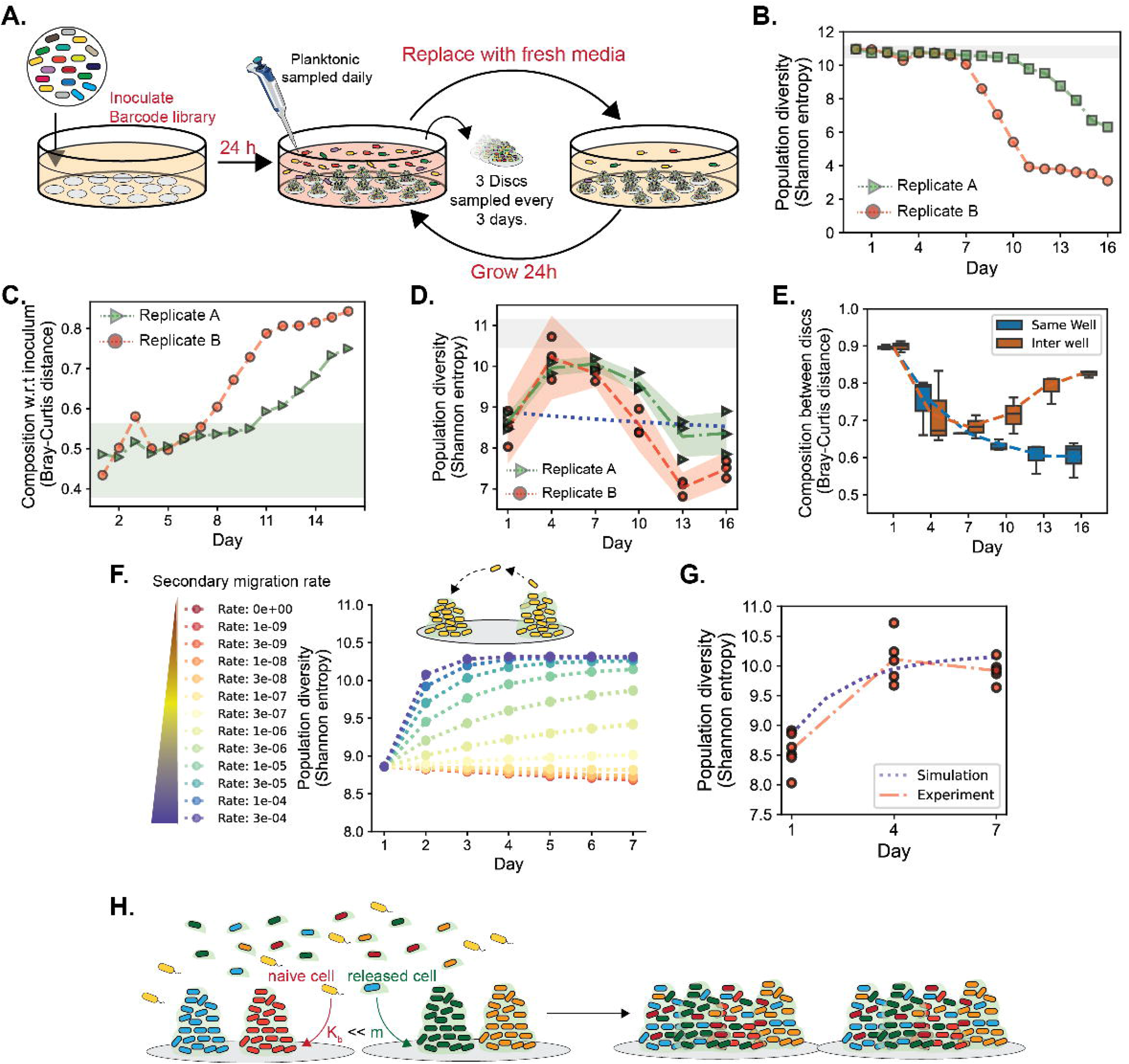
Temporal tracking of biofilm population’s homogenisation driven by secondary migration: **(A)** We periodically replaced the spent media with fresh media and left the culture static to grow for 24 hours in replicate wells each with 18 discs. We repeated 16 replacement cycles, sampling the planktonic population every cycle and three discs per replicate every three cycles. **(B)** Population diversity measured as Shannon entropy for biofilm-associated planktonic populations for replicate A (green), replicate B (red). Shaded grey regions indicate the 99% confidence interval (mean ± 2.52*standard deviations) of Shannon entropy measured across four independent wells at *t* = 0 in the no-bottleneck shake-flask control. **(C)** Compositional dissimilarity between the inoculum and populations recovered from the biofilm-associated planktonic phase for replicate A (green) and replicate B (red). Shaded grey regions indicate the 99% confidence interval (mean ± 2.52*standard deviations) of compositional dissimilarity between the inoculum and populations measured across four independent wells at *t* = 0 in the no-bottleneck shake-flask control. **(D)** Population diversity measured as Shannon entropy for disc-associated biofilm populations for replicate A (green) and replicate B (red) over time. The dashed line represents the mean Shannon entropy and the shaded region represents mean ± standard deviations for 3 different sampled discs from each replicate. The blue dashed line represent the expected change in Shannon entropy using a temporal simulation performed with optimal binding rate estimated in our initial binding simulation. **(E)** Bray–Curtis distances between disc-associated populations over time for discs recovered from the same well (blue) and from different wells (orange) of replicate A. Box plots show the median, interquartile range, and whiskers extending to 1.5× the interquartile range. **(F)** Simulated Shannon entropy of the disc-associated population as a function of increasing secondary migration rates. Color gradient (red to blue) denotes an increasing migration rate parameter in the binding-growth model. **(G)** Comparison between experimental (red) and simulated (blue) Shannon entropy using the optimal migration rate. Red circles represent Shannon entropy from 3 discs each from 2 biological replicates at Cycles 1, 4, and 7, highlighting the variance in observed diversity dynamics. **(H)** Cells released upon disturbance re-enter (immigrate into) the biofilm at a rate m, which is three orders of magnitude higher than naïve cells, driving spatial homogenization of the biofilm population.

While mature biofilms allow external entry of new cells over time, edge dominated growth is proposed to maintain segregated population structures over short timescales^12,18,20–22,24^. Therefore, we expected the initially observed distinct population structure and low diversity to be maintained over short timescales. However, during the period of planktonic stability (7 days in replicate A and 10 days in replicate B), the diversity and number of barcodes recovered in discs increased significantly (**Figure 3D** & **Supplementary Figure 7B,** Kolmogorov-Smirnoff test, p-value < 0.01). Over the same period, the population composition between discs in the same well became increasingly similar and homogenized towards the composition of the inoculum (**Figure 3E** & **Supplementary Figure 7C-7D**). The composition on discs between different replicate wells also became increasingly similar (**Figure 3E** & **Supplementary Figure 7C-7D**), reflecting a deterministic mixing of lineages resulting in homogenization of the biofilm population towards the same inoculum.

A temporal simulation using the initially estimated optimal binding rate as the rate of binding of released cells predicted no increase in disc population diversity (**Figure 3D, blue line**). We therefore hypothesized that the released cells re-entered the biofilms at a rate higher than initial binding. By introducing a new secondary seeding (migration) rate in our model, we captured the increase in the disc population diversity over time (**Figure 3F**). Upon minimizing the Root Mean Square Error (RMSE) between the simulated and experimental change in diversity, a migration rate of 3.16*10^-5^ provided the best fit (**Supplementary Figure 7E**), replicating the increase in diversity observed (**Figure 3F**). Therefore, cells dislodged from an already-established biofilm following perturbation had three orders of magnitude higher biofilm re-entry rate (∼10^-5^) compared to the initial binding rate (∼10^-8^) of naïve cells (**Figure 3G**). This suggested that physical disturbance facilitated robust exchange of cells to drive erosion of segregation and lineage homogenization in biofilms (**Figure 3H**).

### 2.4 Interplay between adaptive sweeps and spatial shielding during adaptive evolution in biofilms

Following homogenization, both biofilm-associated planktonic and biofilm populations underwent a rapid decline in diversity (after 7 days for replicate A and 11 days for replicate B) (**Figure 3B-3D**). While the discs within the same well became increasingly similar, the compositional distance between discs from different wells increased, reflecting a local intra-well change in population composition (**Figure 3E**). Distinct subsets of barcodes increased in frequency in each replicate experiment, with the dominant lineage reaching ∼20% in replicate A and ∼50% in replicate B (**Figure 4A-4B** & **Supplementary Figure 8A-8B**).

**Figure 4.**
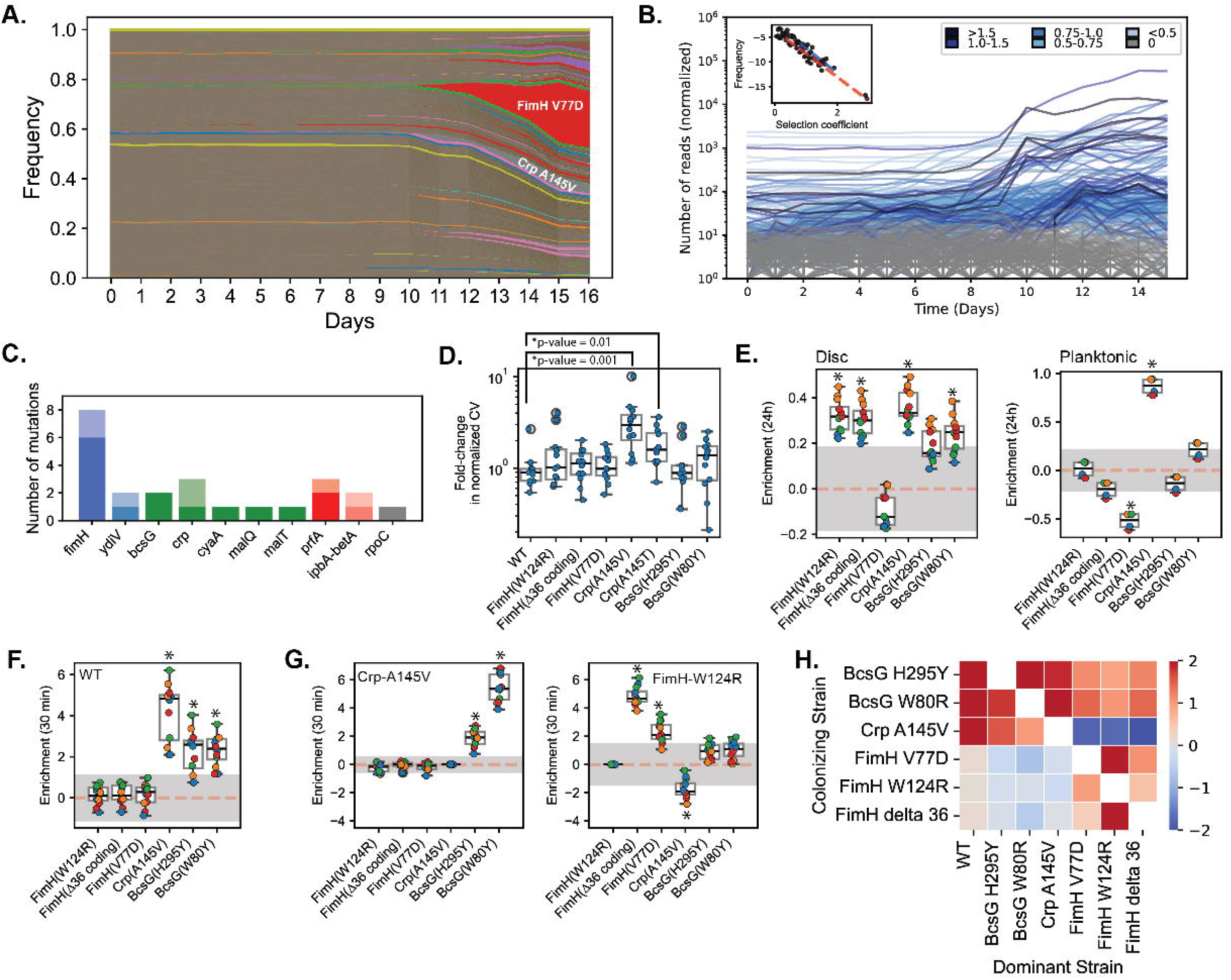
Diversity of adaptive targets and mechanisms underlying biofilm evolution. **(A)** Muller plot showing the temporal dynamics of barcoded variants in biofilm-associated planktonic population in replicate A, where each colored region represents the relative frequency of a single variant over time. Repeated but separated colors represent lineages with different barcodes. **(B)** Temporal trajectories of individual variants classified as beneficial (blue; inferred fitness > 0) or neutral (grey). Line color intensity for beneficial variants increases with inferred fitness. (Inset) the frequency of mutations of different effect sizes for both replicates. Selection coefficient inference and frequency estimates are described in the Materials and Methods section. **(C)** Number of independently arising mutations per genetic target among variants inferred to have positive fitness. Dark shading indicates clones carrying a single mutation in the indicated gene, while light shading indicates clones carrying additional mutations in other loci (**Supplementary Table 3**). **(D)** Biofilm biomass quantified by crystal violet staining and normalized to colony-forming units (CFUs) for beneficial variants, shown relative to the ancestral *Escherichia coli* LF82 strain. Crystal violet staining and CFU enumeration were performed after 24 h of biofilm formation for each variant (see Materials and Methods; raw data shown in **Supplementary** Figure 12). Because biofilm biomass scales with total cellular biomass, crystal violet optical density values were normalized by dividing by total CFUs. Fold changes in normalized crystal violet absorbance were then calculated relative to the wild-type *E. coli* LF82 strain. Each point represents an independent biological replicate (n = 12). Asterisks indicate statistically significant differences compared with wild type (two-sided Mann–Whitney U test with Benjamini–Hochberg correction for multiple testing). **(E)** Enrichment scores (fitness) of beneficial variants during initial colonisation of empty silicone discs (left) and in the planktonic population (right) after 24 h of inoculation with variant pool. **(F)** Enrichment scores during invasion into discs pre-colonized for 24 h by wild-type *E. coli* LF82, measured after 30 minutes of inoculation. **(G)** Enrichment scores during invasion into discs pre-colonized for 24 h by the Crp A145V variant (left) and into discs pre-colonized by the FimH W124R variant (right), measured after 30 minutes of inoculation. **(H)** Heatmap showing the mean enrichment score of beneficial variants (y-axis) invading biofilms pre-colonized by different dominant variants (x-axis). Enrichment scores were calculated as the log change in frequency of each variant relative to four independently barcoded neutral wild-type lineages between the beginning and end of each experiment. In panels C–E, each point represents an independent measurement from three replicate wells against four reference barcodes. The grey shaded area represents the mean +/- 1.96*standard deviation of the enrichment measured for 4 neutral barcodes with no mutations. The variants with mean enrichment higher or lower than the shaded threshold have significantly higher or lower enrichment in the given context (2-tailed students’ t-test p-value < 0.05 for significant values). In panel F, points represent mean enrichment values averaged across four reference barcodes and three replicate wells (raw data in **Supplementary Fig. 13**). Box plots show the median, interquartile range, and whiskers extending to 1.5× the interquartile range (n = 3 × 4).

The observed decrease in diversity, together with the local expansion of specific lineages was consistent with adaptive evolution. To test this, we quantified the lineage fitness using two independent, semi-quantitative and highly correlated approaches (**Supplementary Figure 9A**; Spearman correlation coefficient 0.88). Lineages with positive fitness enriched significantly in both replicates (**Figure 4B** & **Supplementary Figure 8B, 9B-9C**, replicate A & B: Kolmogorov-Smirnoff test, p-value < 10^-10^). At the end of the experiment, we sequenced 106 lineages with unique barcodes from replicate A (with 19 variants recovered from the planktonic and 87 lineages recovered from two discs) (**Supplementary Table 3**). While all predicted beneficial lineages (n=19) harbored mutations, only 22.3% (19/85) of the predicted neutral lineages were mutated (Fisher’s exact test, odds ratio infinite for a predicted beneficial lineage to be mutated, p < 10⁻⁶). Overall, we classified 233 and 135 lineages across replicates as beneficial (**Figure 4B & Supplementary Figure 8B**). The co-occurrence of multiple beneficial lineages demonstrated extensive clonal interference.

Although the dominant beneficial lineage emerged at different times between the two replicates (day 11 for replicate A and day 7 for replicate B, **Figure 4A-****4**B and **Supplementary Figure 8A-8B**), the distribution of fitness effects (DFE) between replicates was similar. The DFE was well described by an exponentially decreasing function (R² > 0.9), indicating that mutations conferring larger fitness benefits were increasingly rarer (**Figure 4B, inset**).

At the end of the experiment, predicted fit lineages accounted for 62.4% and 82.5% of the planktonic population in Replicates A and B, respectively, but only 33.0% and 31.5% in corresponding disc-associated biofilms. Although shared lineages enriched relative to the inoculum initially represented 3.47% +/- 0.01% in segregated populations per disc, they expanded to 30.29% +/- 0.05% by the end (**Supplementary Fig. 10A-10B**). This increase closely mirrored the expansion of fit lineages (s > 0), which constituted 84.5% +/- 4.03% of the shared population. Crucially, all barcodes shared across multiple discs carried mutations (100%, 20/20).

Despite the adaptive sweep, the disc-associated biofilm populations retained substantially higher diversity than the biofilm-associated planktonic counterpart (Shannon entropy: planktonic replicate A= 3.1 & replicate B = 6.3; Disc replicate A = 7.48+/-0.17 & replicate B = 8.36 +/- 0.42). While stochastically expanded single-disc lineages comprised 75.6% of the disc population at the start of the experiment, 21.3% of the population was still composed of singled-disc enriched lineages at the experiment’s end (**Supplementary Figure 10B**). These lineages preserved uniquely on a single-discs were mutated significantly less frequently than shared lineages (**Supplementary Figure 10C;** 4/34 (11.76%) mutated; Fisher’s exact test, p < 10⁻⁵). Therefore, lineages with beneficial mutations increasingly spread during the adaptation, while the biofilm structure retained a substantial fraction of the enriched non-mutated local diversity.

### 2.5 Mutations targeting distinct stages of the biofilm life cycle drive adaptive evolution

Amongst the 40 mutated lineages detected in the planktonic and the biofilm populations, multiple signatures of convergence emerged, providing further evidence of adaptive evolution (**Supplementary Figure 11A** and **Supplementary Table 3)**. The most frequent target was *fimH*, the gene coding for the type I fimbriae tip pilin, with five different mutations arising in eight lineages (**Figure 4C**). All mutations mapped to the lectin domain responsible for FimH mannose-specific binding. Mutations in FmH lectin domain were shown to modulate biofilm formation on different surfaces in *E. coli* MG1655 and LF82 strains ^26,32^.

We also observed frequent mutations in the global regulator *crp*, showing residue-level convergence: with mutation A145V in two lineages and A145T in one (**Figure 4C**). The gene *cyaA*, encoding the enzyme synthetizing cAMP, the activator of Crp, was also mutated. Crp mutation A145V is known to compensate for the loss of a sugar phosphotransferase system, implying that this mutation may alter sugar transport or metabolism ^33^. Consistently, other sugar transport and metabolic genes were also detected, including *bcsG*, *malT*, *malQ*, and *mglB* (**Figure 4C**). Additionally, mutations occurred in genes *rpoC* (RNA polymerase), *prfA* (release factor 1, translation), *betA*, and *ompA* (osmotic stress) in inferred beneficial lineages (**Figure 4C** & **Supplementary Figure 11**).

To decouple biofilm-specific adaptation from media-driven evolution, we sequenced beneficial variants from biofilm-free, shaken-flask control. Strikingly, all 36 distinct sequenced lineages harbored mutations in the anaerobic regulator *arcAB* (**Supplementary Table 4**). In contrast, *arcAB* mutations were absent in biofilm-derived lineages, confirming adaptation to biofilm-specific selective pressures.

To understand adaptive mechanisms, we first tested the biofilm forming capacity of 7 variants using the crystal violet assay (3 *fimH*, 2 *crp*, and 2 *bcsG* mutations). Only the variants CRP A145V and A145T showed significantly greater biofilm-forming capacity than the ancestral strain (**Figure 4D, Supplementary Figure 12,** Kolmogorov-Smirnoff test, p-value < 0.01). This suggested that other complex adaptive mechanisms than just biofilm formation were at play for different beneficial mutations. To understand the fitness effects of the mutations, we performed pooled competition assays with six variants (FimH W124R, FimH Δ36 coding (86-121/903 nt), FimH V77D, Crp A145V, BcsG H295Y, BcsG W80Y) and measured the competitive index of each variant in different ecological contexts.

We first inoculated the pool into fresh medium containing new silicone discs and allowed biofilms to form for 24 hours, mimicking initial colonization. All variants except FimH V77D had significantly higher fitness in the biofilm fraction relative to wild type (**Figure 4E**, left). Only the *crp* variant displayed a significantly higher competitive index in the planktonic phase (**Figure 4E**, right), suggesting an impact on migration or growth.

To quantify a variant’s ability to migrate into a pre-existing biofilm, we introduced the mutant pool into wells with discs with wild-type pre-grown biofilms. Only the *crp* and *bcsG* variants had significantly higher fitness in the invasion assay (**Figure 4F**). However, the *crp* and *bcsG* variants showed a significant competitive advantage only when the resident biofilm was formed by wildtype LF82, and *crp* or *bcsG* variants (**Figure 4G–4H** and **supplementary Figure 13**). These mutants lost their advantage—and the *crp* variant even had negative fitness—when the biofilm was formed by *fimH* variants (**Figure 4G–4H** and **supplementary Figure 13**). Conversely, all *fimH* variants succeeded in biofilms formed by other *fimH* variants, including *fimH* V77D that performed poorly in the initial pooled biofilm-formation competition assay (**Figure 4E**, **4G–4H and supplementary Figure 13**).

Thus, adaptive evolution selects for mutations targeting distinct stages of the biofilm lifecycle. There are frequency-dependent ecological contingencies and interactions, whereby variants with mutations affecting similar traits preferentially colonize biofilms established by variants of the same functional class.

### 2.6 Homogenization dynamics upon antibiotic exposure

Biofilm-associated infections are notoriously recalcitrant to antibiotics and are therefore potent incubators for the emergence of resistance ^10,11,28,34^. To evaluate the impact of exposure to antibiotics on the temporal lineage-level dynamics, we subjected both evolving biofilms and well-mixed flask controls to intermittent 24-hour lethal dose (5X MIC) of amikacin during the media replacement cycles (**Figure 5A and Supplementary Figure 14A**). As previously reported using a similar set-up^26^, a significant increase in resistance was observed in the biofilm-derived population after the third treatment cycle (day 7), whereas no detectable resistance emerged in well-mixed flask controls even after 5 cycles (**Figure 5B** & **Supplementary Figure 14B**).

**Figure 5.**
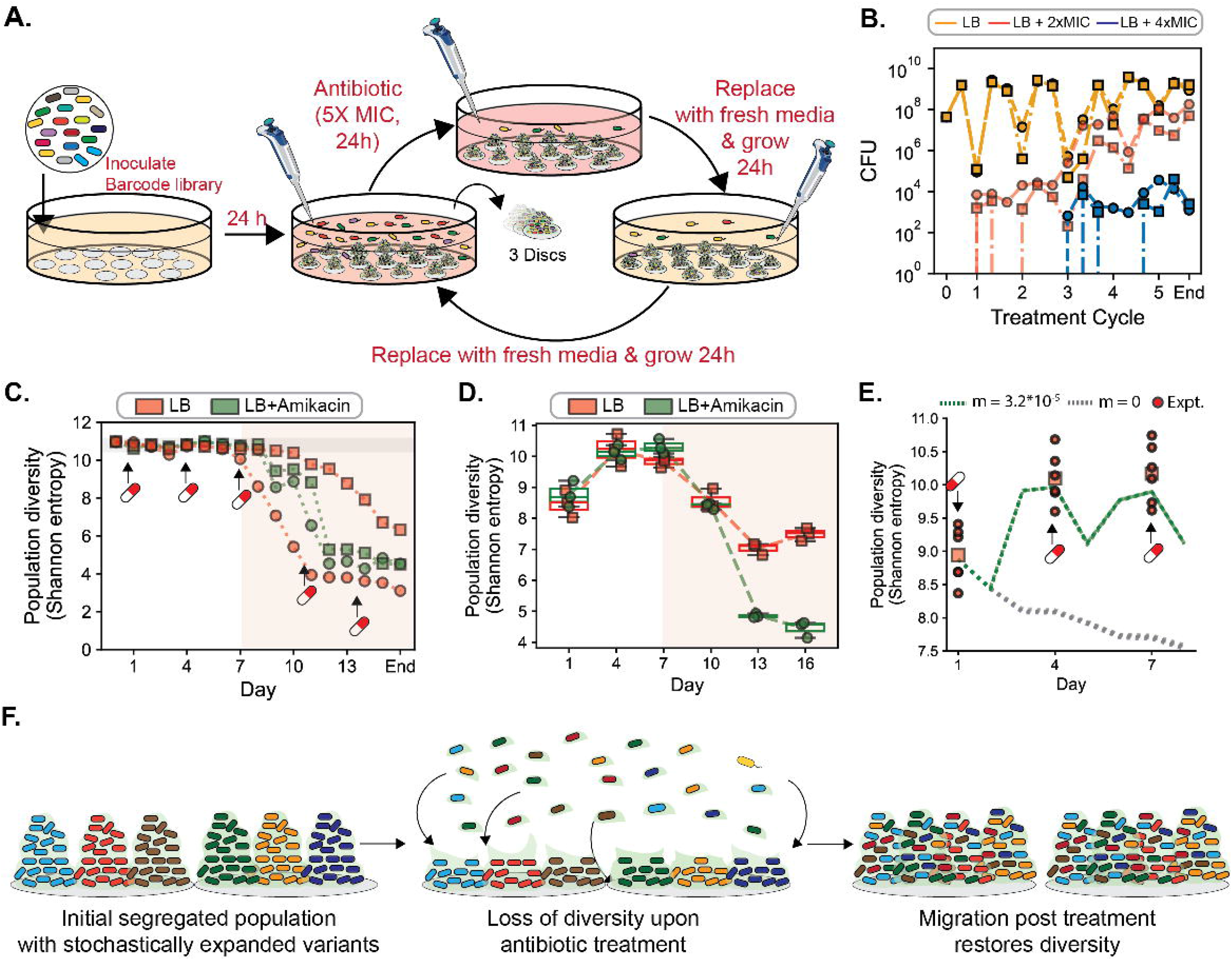
Temporal population dynamics under antibiotic treatment. **(A)** Schematic of the experimental design incorporating periodic antibiotic treatment between media replacement and growth cycles. After initial biofilm formation, populations were exposed to amikacin at five times the minimum inhibitory concentration (MIC) for 24 h. Following antibiotic treatment, the medium was replaced and cells were allowed to regrow for two consecutive 24 h periods. This antibiotic treatment and wash– regrowth cycle was repeated five times. **(B)** Colony-forming units (CFUs) measured over five antibiotic treatment cycles for populations in planktonic populations from wells containing biofilms (right). CFUs were quantified following growth in LB alone (yellow), LB supplemented with 2× MIC (red), or 4× MIC (blue). Trajectories with squares and circles distinguish the two biological replicates. **(C)** Population diversity measured as Shannon entropy for biofilm-associated planktonic populations for biofilms grown without antibiotics (red) and with antibiotics (green). The circle and square replicate an independent biological replicate for each case. The horizontal shaded grey regions indicate the 99% confidence interval (mean ± 2.52*standard deviations) of Shannon entropy measured across four independent wells at *t* = 0. The red-white pill represents the time-point prior to the antibiotic treatment cycle. The vertical red shaded area marks the period where significant exponential increase in resistance frequency was observed. **(D)** Population diversity measured as Shannon entropy for disc-associated biofilm populations sampled from experiments conducted without amikacin (red) and with amikacin (green) for replicate A. The vertical red shaded area marks the period where significant exponential increase in resistance frequency was observed. **(E)** Comparison between experimental (red) and simulated Shannon entropy using the optimal migration rate estimated in Figure 3 (green) and no migration (grey). Red circles represent Shannon entropy from 3 discs each from 2 biological replicates at Cycles 1, 4, and 7, highlighting the variance in observed diversity dynamics and the red square representing the mean Shannon entropy on discs between replicate discs. The red-white pill represents the time-point prior to the antibiotic treatment cycle. **(F)** A cartoon representation of antibiotic-mediated bottleneck followed by migration-mediated restoration of diversity before the exponential increase of resistance frequency (i.e., before day 7 and third antibiotic treatment). In panels E, points represent measurements from individual discs (three discs per well), and lines indicate the mean across replicate wells. Discs were destructively sampled at each time point. All box plots show the median, interquartile range, and whiskers extending to 1.5× the interquartile range.

Following initial exposure, 0.35 +/- 0.14% of the biofilm population survived, which was ∼35-fold higher than the survival in the shaken-flask control and the biofilm-associated planktonic population (**Supplementary Figure 14C**). After each cycle, the cells released from the biofilm population reseeded the planktonic fraction (Supplementary figure 5). Strikingly, before the start of an exponential increase in resistance frequency on day 7, despite the 99.65% killing within the biofilms, the diversity, number of unique barcodes and composition in the reseeded population remained comparable to those of the inoculum and the untreated control (**Figure 5C** & **Supplementary Figure 15A-15B).** Therefore, the biofilm structure effectively protected the resident population against a loss of diversity due to the antibiotic-mediated killing. During this pre-resistance period, localized diversity within biofilms on individual discs increased and population composition between discs homogenized at rates identical to the untreated controls (**Figure 5D & Supplementary Figure 15C-15E**).

To quantify the effect of the antibiotic treatment, we integrated a 24-hour antibiotic treatment-induced killing step in our stochastic binding-migration model (**Figure 5E-5F and Supplementary methods 5.4**). The model predicted that in the absence of migration, there would be a progressive decrease in population diversity after each antibiotic treatment (**Figure 5E**). In contrast, incorporating active inter-disc migration into the model dynamically restored the diversity, accurately matching the experimental values (**Figure 5E**). Thus, while the antibiotic treatment initially depleted the diversity on discs, active migration between discs continuously redistributed the locally preserved lineages in spatially segregated different subsections of the biofilm population (on discs) to restore local diversity (**Figure 5F**).

### 2.7 Expansion of antibiotic resistance is accelerated by intra-biofilm migration

The exponential increase in resistance frequency after day 7 was accompanied by the rapid enrichment of several dominant variants in both replicates (**Figure 6A & Supplementary Figure 16A**). Whole-genome sequencing of 87 unique enriched lineages from replicate A revealed a significant genetic convergence. Of these, 86 lineages harbored a mutation in the *sbmA* gene (a peptide antibiotic transporter), with the majority representing loss-of-function mutations (45 out of 58 unique mutations included 15 nonsense, 23 frameshift, and 7 structural deletions; **Supplementary Table 5**). *sbmA* loss-of-function mutations were previously repeatedly identified in multiple studies with both planktonic or biofilm populations exposed to aminoglycosides^25,26,35–37^. Variants with *sbmA* mutations had significantly higher minimum inhibitory concentration for amikacin (**Supplementary Figure 16B**).

**Figure 6.**
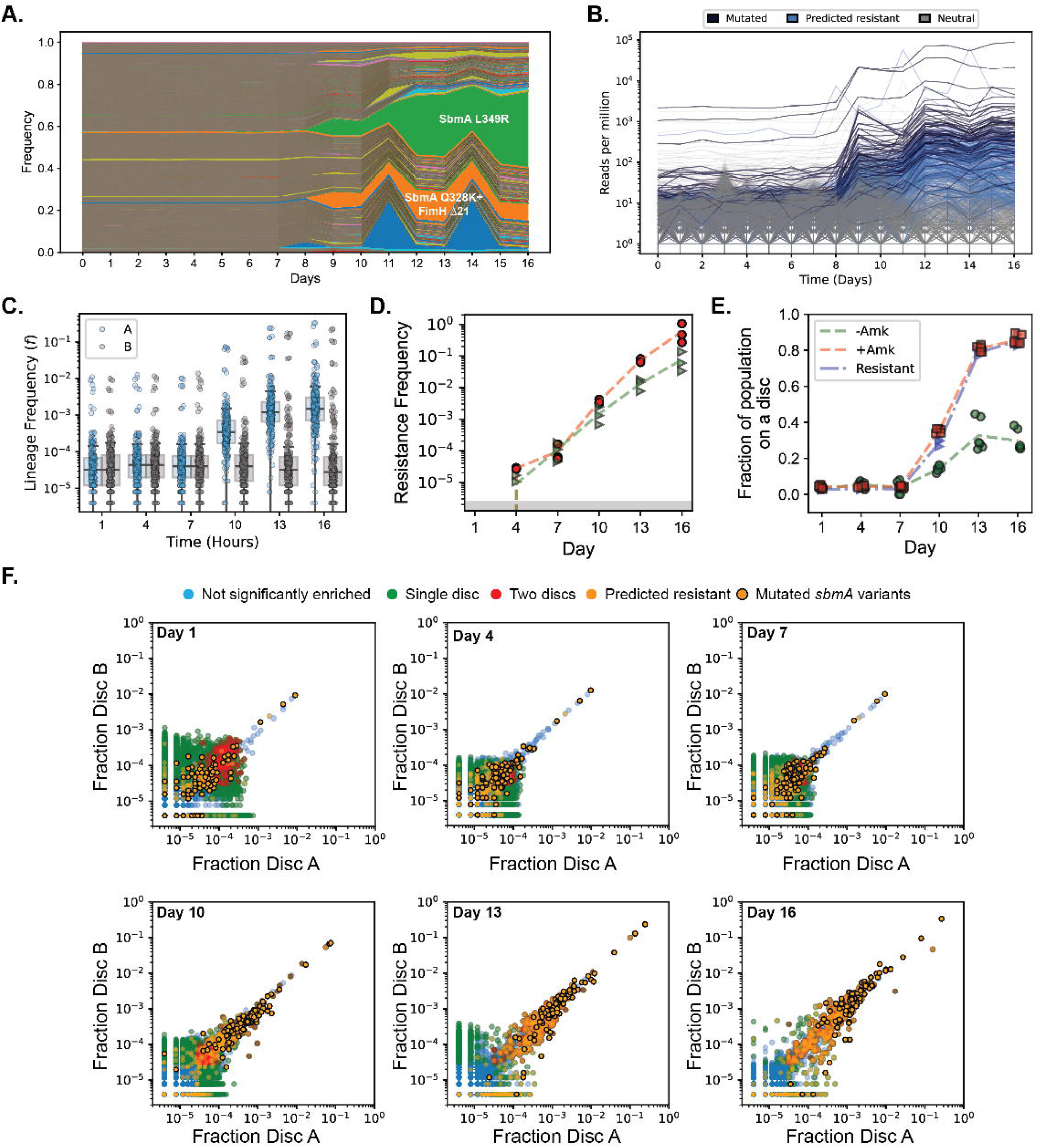
Emergence of resistance: **(A)** Muller plot showing the temporal dynamics of barcoded variants in biofilm-associated planktonic population in replicate A of the temporal biofilm experiment with antibiotic treatment, where each colored region represents the relative frequency of a single variant over time. Repeated but separated colors represent lineages with different barcodes. **(B)** Temporal trajectories of sequenced resistant with *sbmA* mutations (dark blue), predicted resistant variants (light blue) or neutral variants (grey). Predicted resistant variants are ones with fold enrichment comparable to the sequenced lineages (p-value > 0.05). **(C)** Frequency of the sequenced resistant lineages with *sbmA* mutations recovered from Replicate A (dark blue lineages in Figure 6B) in the discs recovered from the experiment replicate A (light blue) and in the discs recovered from experiment replicate B (grey). Box plots show the median, interquartile range, and whiskers extending to 1.5× the interquartile range. **(D)** Resistance frequency estimated for replicate A (red circles) and replicate B (green triangles). The three points are 3 independent discs from each replicate, and the dashed lines represent the mean of the three discs. **(E)** Fraction of the disc-associated population represented by variants significantly enriched on two discs from the same well for the experiment without (green) and with (red) Amikacin treatment, and that represented by predicted resistant variants (blue). The three points are 3 independent discs from each replicate and the dashed lines represent the mean of the three discs. **(F)** Pairwise comparison of barcode counts over days between two discs recovered from the same well. Variants without any significant enrichment are blue, significantly enriched on one disc are green, variants significantly enriched on both discs are red, variants with predicted resistance are orange, and variants with sequence-verified resistant *sbmA* mutations are orange with black edge colors.

Benchmarking against the fold-enrichment of the confirmed *sbmA* variants, we predicted 428 lineages in replicate A and 396 lineages in replicate B as resistant (**Figure 6B** & **Supplementary Figure 16C)**. Confirmed and predicted resistant lineages displayed an increase in frequency strictly within replicates where they arose, validating that resistance arose via *de novo* mutations rather than pre-existing variation (**Figure 6C** & **Supplementary Figure 16D**). Therefore, resistant mutations were recruited in parallel across multiple independent lineages.

An increased resistance frequency was observed in the biofilm-associated planktonic population as early as day 2 and in the recovered discs on day 4, following the first treatment cycle (**Figure 5B** and **Figure 6D**). However, the distribution of the frequencies of resistant lineages in the replicate they arose were comparable to that in the other replicate where they were non-resistant and were also comparable to predicted non-resistant lineages (**Figure 6C** & **Supplementary Figure 16D-16E).** Through Day 7, the fraction of shared enriched lineages between discs remained low and the discs preserved substantial local diversity comparable to the untreated controls (**Figure 6E & 6F).** Despite the presence of resistant variants early on, the initial migration rates were also comparable to the untreated controls (**Figure 5D**). After day 7, there was an exponential increase in resistance frequency at a rate of 0.36+/-0.047 day^-1^ (**Figure 6D**). The increase in resistance was tightly coupled with the increase in the fraction of shared enriched lineages between discs, with the shared population being almost entirely composed of mutated and predicted resistant lineages (**Figure 6E-6F & Supplementary Figure 17**). Therefore, once the resistant lineages achieved a certain frequency, they were increasingly shared by secondary migration between discs resulting in an exponential spread of resistance in the biofilm population.

Barcode-resolved lineage dynamics revealed the parallel emergence of resistance-conferring mutations across hundreds of initially low-frequency lineages, rather than a selective sweep driven by a single resistant variant. Subsequent inter-disc migration facilitates the rapid spatial dissemination of *de novo* resistant lineages, resulting in an exponential increase in resistance frequency.

## 3. Discussion

By combining high-throughput barcode tracking with spatial sampling, we quantitatively resolved the lineage-level dynamics that shape surface-attached biofilms population. We identified three sequential processes operate at different timescales, which remain consistent even with intermittent antibiotic treatment: (i) an initial stochastic binding associated with bottlenecks and segregation, (ii) a period of secondary migration-mediated homogenization, and (iii) a deterministic enrichment of adaptive lineages with replicable evolutionary dynamics.

While previous microscopy and modeling have described stochastic binding^18,21–24^, our results quantitatively describe its evolutionary consequences, demonstrating how it drives diversity loss and the lineage-level re-structuring of population, resulting in an effective population size three orders of magnitude smaller than the actual population size (**Figure 1-2**). By combining our data with a stochastic binding growth model, we show the multi-orders-of-magnitude stochastic advantage of founders that drives spatial segregation ^23^ and niche preemption, whereby early-arriving lineages preclude future colonisation ^38^ (**Figure 2**). By modulating the modeling and experimental parameters in our framework, we can now dissect the roles of surfaces, strains^6^, inoculum density^19^, nutrient flux^22^, and surface chemistry ^24^ on colonisation.

Previous studies have shown that while new cells can enter established biofilms, growth predominantly at the colony boundaries largely maintains the segregated population structure over short timescales ^12,18,20–22,24^. We uncovered a new, previously underappreciated phase of the temporal dynamics, where physical disturbance triggers secondary migration and dynamic reseeding, to progressively homogenize the population and eroding the spatial segregation established during initial colonization (**Figure 3**). Following a physical perturbation, cells dislodged from the biofilm matrix exhibit a three-order-of-magnitude higher rate to reintegrate into pre-existing biofilms (k_mig_ ∼ 10^-5^) compared to the initial attachment rate of “naive” planktonic cells (k_binding_ =∼10^-8^). While initial biofilm entry requires significant regulatory adaptation, cells released by perturbations are likely pre-adapted, allowing accelerated re-colonization. While naturally dispersing cells are known to be physiologically distinct from planktonic or biofilm-bound cells, we demonstrate that they possess an increased capacity to reseed new biofilms and quantify the resulting consequences on the population dynamics.^39,40^.

Subsequently, adaptation within biofilms did not converge to a single strategy and differed significantly from the adaptation in well-mixed cultures (**Figure 4**). Mutations target distinct phases of the biofilm life cycle including attachment (*fimH*), migration into established communities (*crp* & *bcsG*), and success within specific niches (*fimH V77D*). Strikingly, the fitness advantage conferred by these mutations is contingent on community composition: mutant lineages are preferentially favored when surrounded by cells carrying the same or similar mutations, revealing a strong “like-favors-like” ecological effect. For example, while *fimH* mutants successfully colonize biofilms composed of *fimH* variants, they lose their advantage when invading biofilms composed of *crp* & *bcsG* variants (**Figure 4)**. Therefore, evolution in biofilms promotes simultaneous emergence of distinct adaptive strategies with localized ecological contingency and frequency dependence.

While biofilms are notoriously recalcitrant to antibiotic therapy and often serve as reservoirs for the emergence of resistance ^11,28^, we show that their role extends far beyond just passive physical shielding. Despite a 99.65% decimation of the biofilm population under lethal antibiotic exposure, secondary migration remains active at a rate comparable to the absence of antibiotics. The secondary migration dynamically counteracts against a local loss of diversity by promoting dynamic exchange of lineages preserved in different local subsections of the biofilm (**Figure 5**).

Previous studies characterized the elevated frequency of resistance in biofilms ^25,26,41,42^. Here, high-resolution lineage tracking enabled us to map the simultaneous *de novo* emergence of resistance across hundreds of independent lineages (**Figure 6**). In addition to the previously reported elevated mutation rates in biofilms ^26^, our findings establish that initial segregation and active migration also critically contribute to this parallel emergence of resistance by shielding the diversity to maintain a high effective population size or Ne. Furthermore, once *de novo* resistant variants achieve local dominance, the same migration-mediated exchange drives an exponential dissemination of resistance across the biofilm population (**Figure 6**). Therefore, the biofilm environment does not merely shield individual cells; rather, the secondary migration actively reinforces the evolutionary potential of the population. In clinical settings exposed to intermittent or turbulent fluid flow—such as indwelling catheters or the urinary tract—this disturbance-mediated exchange and dissemination of variants may represent a key mechanism accelerating the rapid spread of recalcitrant infections and drug resistance ^7,16^.

Leveraging high-resolution lineage tracking unlocked a level of quantitative precision that was previously inaccessible, enabling us to disentangle the contributions of initial attachment, growth, spatial migration, and adaptation on biofilm development across different timescales. Together, these findings provide a quantitative framework for understanding the remarkable persistence of biofilm populations: spatial structure and migration simultaneously increase the ecological resilience and the evolutionary adaptation potential by buffering populations against external perturbations, preserving local diversity, and enabling both the repeated parallel recruitment of beneficial variants and their subsequent spread.

## Methods

### Barcode library construction

The barcodes were introduced in the safe site 9 (ss9) region of the *E. coli* genome using Cas9-mediated recombineering. The safe site 9 site is proposed to be a neutral insertion site ^1^. We used Cas9-mediated recombineering to introduce the barcodes in *E. coli* strain LF82.

### Plasmid library construction

The plasmid expressing the ss9-targeting gRNA was purchased from Addgene (https://www.addgene.org/71656/). We constructed a repair template encoding the barcode in the SS9 integration plasmid with the pBR322 backbone described by Bassalo *et al*, 2016 ^1^. The plasmid contains sequences with 600 bp of homology upstream and downstream to the MG1655 SS9 integration site containing a uvGFP cassette. We first replaced the MG1655-upGFP cassette with a 1200 bp long sequence homologous to the LF82 SS9 site. We amplified the LF82 SS9 site using the primers LF82_SS9_for: ACTCGGTTGAGAATACGCCG and LF82_SS9_rev: GCCTACGATTACGCATG-GCT’. We amplified the pBR322 backbone using primers pBR322_SS9bb_for (5’-GTCTGTCCAGCGCGTCGGCA – 3’) and pBR322_SS9bb_rev (5’-CCAGGGGCTGATTTTAA-CTT – 3’). All PCR amplifications were setup using the Q5 high fidelity polymerase using their standard proposed protocol with PCR cycling conditions: initial denaturation: 95°C-2 min, 30× (95°C-30 s, Tm-30 s, 72°C-1 min), and final extension: 72°C-2 min. We determined the PCR Tm using NEB’s Tm calculator for Q5 high fidelity polymerase.

1 µL of the *Dpn*I was directly added to the PCR mixture for every 25 µL of the PCR reaction and the mixture was incubated at 37^°^C for 2 hours or overnight. All PCR products were then purified using the Qiagen gel purification kit using the manufacturer’s protocol (Catalog #28704). Then, the SS9 barcode library was cloned using the standard NEBuilder cloning protocol using the manufacturer guidelines for reaction setup. For the DNA assembly reaction, we mixed at-least 100 ng of the vector backbone with two-fold excess of the molar equivalent of the insert. We incubated the reaction for 60 minutes at 50^°^C. 10 μl of the assembly reaction was dialyzed using a 0.45-micron dialysis membrane. The dialyzed mixture was transformed in commercial Lucigen Elite *E. cloni* electrocompetent cells (catalog #60061). 10 µL of the commercial cells were mixed with 40 µL of cold 10% glycerol. Then cells were recovered in 1 mL manufacturer provided recovery media per 50 µL transformation reaction. Several dilutions were plated on LB Agar + plasmid specific antibiotics. In case of plasmid construction, multiple single clones were isolated, and the correct plasmids were extracted after sequence verification.

To prepare the repair template, we amplified the plasmid backbone with primers pSS9_zeobb_rev: CTGCGCCAGAGGTAGGATTGAAAA and pSS9_zeobb_for: CAGCATCAAT-AATCAACGCGGAATGATGCAGAGATGTAAG. The amplification deleted a small region homologous the SS9 site targeting gRNA and the associated PAM sequence. The deletion provides immunity from Cas9 gRNA-mediated DNA double strand break. The primers also add homology required for the Gibson assembly to the Barcode zeocin cassette. We coupled the barcodes to a Zeocin resistance cassette^2^ with PCR using the primers Zeo_BC_for: AGAGCGTTTTCAATC-CTACCTCTGGCGCAGTTNNNNNATNNNNNATNNNNNATNNNNNATGTCATCGCTTGCATTAGAAAGG and Zeo_BC_rev: CAGCATCAATAATCAACGCGGAATGATGC-AGAGATGTAAG. This amplified barcode was cloned in the pBR322 backbone with homology to the LF82 ss9 using the NEB HiFi assembly as described above. For the plasmid barcode libraries, ∼500,000 colonies were collected using scraping with liquid LB. The barcode library was then extracted using Qiagen miniprep extraction kit. The repair template was then amplified using the primers SS9_600_for: ACTCGGTTGAGAATACGCCG and SS9_600_rev: GCGTGTAAGTTTAGCCG-GATAACG. 1 µL of the *Dpn*I was directly added to the PCR mixture for every 25 µL of the PCR reaction and the mixture was incubated at 37°C for 2 hours. All PCR products were then purified using the Qiagen gel purification kit (Catalog #28704).

### Genome library construction

We introduced the plasmid pAM053 ^3^. The pAM053 plasmid encodes the *cas9* gene expressed under a weak constitutive promoter, the lambda Red recombination proteins under the lambda phage pL promoter, controlled by the temperature sensitive cI857 repressor, and a Chloramphenicol resistance cassette. The pAM053 harbors the pSC101 origin, which is maintained at 30^°^C and cured at 37^°^C. The cells were plates on LB+Agar plates with 34 µg/ml of Chloramphenicol overnight at 30^°^C.

The LF82 strain with the pAM053 plasmid was grown overnight at 30°C in 5 mL cultures. The next morning, the overnight cultures were diluted a 100-fold into fresh LB media with (34 ug/ml chloramphenicol for the ancestor and 8 ug/ml chloramphenicol for the evolved strains). The cultures were grown at 30°C until mid-log optical density (measured at 600 nm) of 0.3 - 0.4. The cells were placed in a shaking water bath set at 42 °C to induce the lambda red recombination operon for 15 minutes. The heat-shocked cells were immediately placed on ice and cooled by swirling and chilled subsequently for at least 15 more minutes. The chilled cells were centrifuged in 50 ml tubes at 7500 x g for 3 minutes. The pellet was washed with 25 ml ice-cold 10% glycerol for every 50 ml culture by suspending the cells and centrifuging at 7500 x g for 3-4 minutes. The washing step was repeated 4 times. A final wash was done with 10 ml ice-cold 10% glycerol. The cells were finally concentrated 170-fold in ice-cold 10% glycerol solution (300 µL for every 50 mL of culture). At least 100 ng of the gRNA plasmid was mixed with at least 100 ng of the repair template and dialyzed in membranes for 30-45 minutes. The dialyzed plasmid and repair template were mixed with 250 µl of competent cells and the mixture was electroporated at 2.4 kV. The cells were finally recovered for 3 hours at 30^°^C in 5 mL LB. Finally, for the barcode library, the cells were subsequently plated on multiple LB agar plates with Zeocin and grown overnight at 37 ^°^C. We scraped 100,000-200,000 colonies in liquid LB with 25% glycerol to have a rich genomic barcode library. The library was immediately frozen at -80°C.

### Evolution experiment

We adapted the evolution protocol from a previous study ^4^.

#### Preparing of the wells

Two wells of two 6-well plates (4 wells in total, 2 exposed to 5xMIC of amikacin and 2 unexposed) were sheeted with eighteen silicon discs and sterilized by filling them with 70% ethanol and incubating at room temperature for 15 minutes followed by 30 minutes of UV light exposure.

#### Sample processing and estimation of resistance frequency

All samples were processed in three steps for stocking, analysis and sequencing (i) saved in glycerol, (ii) serially diluted and plated on LB, LB+2xMIC, LB+4xMIC for CFU count, (iii) ∼10^7^ cells were boiled for PCR amplification of the barcode region and further amplicon sequencing. In the experiment with antibiotic treatment, the cells were plated on LB, LB+2xMIC, LB+4xMIC to count the CFU count and estimate of resistance frequency

#### Preparation of the inoculum and initial biofilm formation

A whole cryotube of the barcoded *E. coli* LF82 library was grown in 100 mL of LB overnight at 37°C. A 100 µL of the resulting stationary phase culture was boiled and used to amplify the barcode region by PCR for subsequent amplicon sequencing. The bacterial culture was diluted to OD_600_ = 0.05 and used to inoculate four Erlenmeyer flasks (50 mL each) as well as each of the four wells sheeted with 18 silicon discs (5 mL per well), with two wells each in separate six-well plates. The four Erlenmeyer flasks and the two six-well plates were incubated 24h at 37°C under shaking and static condition, respectively. After 24 hours of growth, the supernatant was sampled and processed (**Supplementary Figure 18**). Each well was then washed twice with LB and three discs were sampled from each well and put in an Eppendorf tube containing 500 µL of PBS. Each Eppendorf tube was vortexed for 1 minute followed by 10 minutes of sonication to dislodge the biofilm cells from the silicon disc. The tubes were then briefly vortexed before (i) being serially diluted and plated for CFU count, (ii) boiling 250µL for PCR amplification of the barcode region and further amplicon sequencing, and (iii) saved in glycerol.

#### Growth and media replacement to track the temporal dynamics in the biofilms with and without amikacin (Supplementary Figure 18)

After washing the wells and recovering 3 discs post 24 hours of growth, we filled two wells in the first six-well plate with fresh LB. Then the samples were statically incubated 24h at 37°C. After 24 hours of growth in a static condition, a sample of the planktonic culture was saved and treated for sequencing as described above. Each well was washed twice with LB and then wells were filled with fresh LB followed by 24 hours of static incubation. The washing and sampling were repeated daily for 15 cyles. Every 3 cycles, each well was washed twice with LB and three discs were sampled from each well and put in an Eppendorf tube containing 500 µL of PBS. Each Eppendorf tube containing a disc was vortexed for 1 minute followed by 10 minutes of sonication to dislodge the biofilm cells from the silicon disc. The tubes were then briefly vortexed and the suspended cells were sampled and processed.

The two wells in the other six-well plate were treated with amikacin. In the experiment with antibiotic treatment, the growth and media replacement cycles were the same as the experiment without antibiotic treatment. However, after the cycle where 3 discs were sampled, instead of adding fresh LB, the media was replaced with LB+5xMIC amikacin and the wells were incubated statically for 24 hours. The resistance frequency of all samples were also measured.

#### Evolution in well-mixed shake-flask controls (Supplementary Figure 19)

In four flasks, the population was inoculated as the experiment with the biofilms are grown for 24 hours with shaking at 150 RPM. After every 24 hour growth, the culture was sampled and processed. The whole population of each flask was then pelleted and washed twice in LB. Five hundred microliter were then transferred in a new flask containing 50 mL LB with a100-fold dilution. In the case of antibiotic treatment, cultures in two flasks the cells were resuspended in LB+5XMIC Amikacin instead in every other cycle. After every cycle, the resistance frequency was estimated.

### Next generation sequencing

The barcodes libraries were sequenced using amplicon sequencing. 5 µl of the boiled samples was used as a template to amplify the sequence including the barcode. The primers used for each amplicon were LF82BC_NXT_for TCGTCGGCAGCGTCAGATGTGTATAAGAG-ACAGNNNNNN-CAATCCTACCTCTGGCGCAG and LF82BC_NXT_rev GTCTCGTGGGC-TCGGAGATGTGTATAAGAGACAGNNNNNNGTCAACACGTGCTCGGATCC. The primers included the overhangs to attach the Nextera adapter. We used the KAPA HiFi polymerase with a reaction mix containing 5 µL of the cell extract, 0.25 µL of 100 mM primers, 25 µL of KAPA polymerase, and water to make the reaction up to 50 µL to set up the PCR reaction. For the PCR we used the following protocol: 95 °C-2 min, n× (98°C-20 s, Tm-15 s, 72 °C1 min), and final extension: 72 °C-5 min. To avoid over-amplification, we performed PCR 1 to attach nextera adapters for 15-20 cycles (where we just observed the PCR product during electrophoresis 1.5% agarose gel). We then performed a second round of PCR to attach the indexes and P5-P7 adapters for to add Illumina adaptors and multiplexing indexes. We mixed 20 µl of KAPA HiFi polymerase, 0.6 μM each of forward and reverse primers with P5 and P7 Nextera Index Kit primers (Illumina), and 5 ul of the PCR 1 product. For the PCR we used the following protocol: 95 °C-2 min, 15× (98°C-20 s, Tm-15 s, 72°C 1 min), and final extension: 72°C-5 min. After PCR II all products were purified using the Qiagen’s gel extraction kit using a 1.5% agarose gel. The purified PCR product was quantified. Then all samples were mixed at equal concentrations to prepare the pool for next generation sequencing. The sequencing was performed using paired-end 2 × 150 np paired end sequencing kit on the Illumina NextSeq platform.

### Genome sequencing

#### Identifying unique clones

At the end of each experiment, we plated several dilutions of the saved samples (10^-4^-10^-6^). We picked 96 individual colonies and suspended them in 10 µl of LB+25% glycerol mix. Using primers LF82_BCsamger_for: GATAATTGAGATCCCTCTCCCTGAC and LF82_BCsamger_rev: GCACT-AAGGCGAACATAAGAGATGG, we amplified the barcoded locus using DreamTaq polymerase with a reaction mix containing 2 µL of the glycerol suspension, 0.25 µL of 100 mM primers, 25 µL of KAPA polymerase, and water to make the reaction up to 50 µL to set up the PCR reaction. We used to LF82_BCsamger_for to sequence the barcode. Clones with unique barcodes were streaked on LB plates and their genome was extracted for sequencing.

#### Whole genome sequencing

From each sample, we sequenced 96 individual clones and retained all clones with unique barcodes. WGS was performed using Illumina technology. Briefly, DNA samples were extracted using the genomic DNA NucleoMag tissue kit from Macherey-Nagel. The whole genome sequencing libraries were prepared and indexed using the Illumina DNA Prep Tagmentation kit with IDT for Illumina DNA/RNA UD Indexes kit A/B/C/D. The pooled libraries were paired-end sequenced to a read length of 2 by 150 bp with Illumina MiniSeq or NextSeq 500/550. The genomes were sequenced at an average depth of 30X.

### Data Analysis

All the code was written in Python 3.7 and packages within Python SciPy, SciKitLearn, Numpy, and Pandas. The codes can be found as submission code 0, and codes A-H covering different analyses.

#### Preprocessing and data normalisation

All preprocessing was performed using a custom analysis pipeline (Submitted Code 0.Preprocessing). We merged the paired-end reads using the Usearch mergepairs algorithm ^5^. Then we aligned the reads to a common reference sequence using the usearch global tool in the usearch package ^5^. In the case of the barcodes, we used the aligned queries to extract the barcode (Submission Code 0. Preprocessing). We constructed a table summarizing the barcodes from all samples. Then we used the shepherd pipeline to cluster the barcodes using the default parameters ^6^. We clustered the barcodes for across 183 samples into one table that we then filtered to normalized and filtered (Biofilm_filtered.csv). The number of reads detected for each sample is listed in Supplementary Table 2. We observed significant differences in read depths.

To account for differences in sequencing depth across the 183 samples, barcode read counts were normalized via multinomial resampling to a fixed total read depth (N). For each sample, the relative frequency of each barcode was calculated by dividing its individual read count by the total reads for that sample. A normalized count table was then generated by resampling from these proportions using a multinomial distribution with a fixed size, 250,000 reads per sample. The optimal read depth was determined by calibrating the normalization depth N against a reproducible baseline of the inoculum (**Supplementary Note 1**).

#### Quantification of barcode diversity and compositional distances

The number of unique barcodes were estimates as the number of all barcodes with normalized read counts greater than 0 in that sample. The unique barcodes detected and cumulative number of unique barcodes analyzed per figure are listed in **Supplementary Table 2**.

Shannon entropy (*H*) was then calculated as:

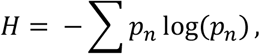

Where the sum is taken over all barcodes detected in the sample. Entropy calculations were performed separately for biofilm-associated planktonic, disc-associated, and flask populations across all replicates.

For analyses comparing populations to the inoculum, Bray–Curtis dissimilarity was calculated using the same normalized barcode frequency vectors, with missing values treated as zeros. All diversity and compositional distance calculations were performed using normalized count tables. The diversity and distance metrics were estimated using the standard Scipy library of python.

#### Quantifying the fraction of population represented by significantly enriched barcodes

To identify variants exhibiting significant enrichment in disc-associated biofilm populations, we first established an enrichment threshold based on the distribution of enrichment values observed in the no –bottleneck shaken-flask control population. Enrichment was defined as the log change in barcode frequency relative to the inoculum. For each of four independent control samples, an upper enrichment cutoff was calculated as the mean enrichment plus 1.96 standard deviations. These values were then combined to estimate a global enrichment threshold by taking the mean of the four cutoffs and adding 1.96 standard deviations of their distribution. This procedure yielded a conservative threshold corresponding to the upper 95% confidence bound of enrichment expected in the absence of selection. Variants with enrichment values greater than or equal to this threshold were classified as significantly enriched in a sample. All calculations were performed using normalized barcode frequency tables.

#### Quantifying fitness

Subpopulations that acquire beneficial mutations and rise to high frequencies, increase deterministically at compared to stochastic changes in the neutral lineages. We estimated the selection coefficient s using two previously published algorithms:

##### (i) Linear regression of log change in frequency w.r.t a wildtype reference^7^

The rate of increase of such fixed beneficial lineages is equal to the slope of the linear regression of log change in the frequency of the variant compared to a neutral lineage i.e., log(p/q), where p and q are frequencies of a variant and a wild-type reference respectively. At the end of the experiment, we sequenced several barcodes isolated from the planktonic and biofilm populations. Of these, the genomes of variants for a subset of barcodes did not harbor any mutations i.e., were wild-type. By combining the reads for all such neutral barcodes, we created a proxy wild-type reference barcode. We estimated the fitness using this proxy barcode as a reference using previously published algorithms ^7^.

For each variant in the population, we estimated the M_v,t_, as:

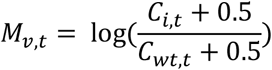

Where, C_i,t_ and C_wt,t_ are the counts of a variant i and wild-type at time t respectively.

To estimate the fitness, we performed a weighted linear regression using weights V_i,t_, where V_i,t_ is the variance based on Poisson assumptions and is measured as:

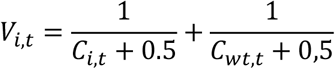

Over the course of the experiment, we estimate a fitness score as the slope of the linear regression of M_i,t_ using V_i,t_.

##### (ii) The popDMS algorithm using statistical methods from population genetics^8^

popDMS is a statistical framework based on population genetics for estimating the functional effects of mutations from DMS experiments ^8^. It models phenotypic selection rounds as analogous to reproduction in natural populations and quantifies the effect of each mutation with a selection coefficient. Assuming additive fitness across mutations, popDMS uses the Wright–Fisher model to link mutation fitness to observed changes in variant frequencies over time. To regularize estimates, it applies a Gaussian prior on selection coefficients, equivalent to ridge regression, a method that shrinks large effect sizes toward zero to prevent overfitting and improve stability of estimates.

Because *de novo* beneficial mutations arise at distinct time points and subsequent clonal interference shifts the mean population fitness over time, fitness was quantified across a rolling window of three consecutive time points for both methods. The maximum inferred slope across all windows was designated as the peak selection coefficient for that lineage (**Supplementary Figure 21 & Supplementary Note 2**). To eliminate spurious estimates driven by lineages with low read counts were removed from the analysis (**Supplementary Note 2**).

#### Estimating significant enrichment with antibiotics

At the end of the experiment with antibiotic treatment, resistant variants are significantly enriched. For replicate A, we had a set of 87 sequence-verified resistant variants. We estimated the enrichment of the verified variants. Candidate resistant variants were identified by establishing a replicate-specific enrichment threshold corresponding to the 5th percentile of confirmed *sbmA* mutants at the end of cycle 5 of the experiment (**Supplementary Figure 22**). Lineages with fold-enrichment values at or above this empirical cutoff were classified as positively enriched resistant candidates (Positive_enriched_A/B == ‘Yes’), ensuring a 95% sensitivity for detecting adaptive mutants.

### Modeling and simulation

#### The binding-growth model (Supplementary Figure 24)

We used a stochastic model to simulate the initial binding and growth of the *E. coli* in the biofilms. The code for the simulation is the supplementary code B. The model tracks the binding and growth dynamics of multiple barcoded lineages in the biofilm-associated planktonic phase and on multiple surface-associated discs.

The simulation is initialized with an inoculation with an inoculum size of N_p,0_ (4.3*10^7^). To determine the number of variants associated with each barcode in the population, we performed a multinomial sampling for a population of size N_p,0_ and using an initial lineage frequency x_0_ of each lineage, estimated by sequencing. Several low frequency barcodes are associated with 0 reads in the inoculum but have significant reads when they enrich on the discs. For these barcodes, we assign a pseudocount = 1, such that they are not excluded from being considered.

At each time step (Δt = 10 minutes), for each barcode, we estimated the number of cells that bind to the available discs at a per cell probability of k_b_. We modeled the binding process as Poisson process, adjusted to the carrying capacity of the disc. Both the biofilm-associated planktonic and the disc-associated cells grow logistically, with specific growth rates µ_p_ and µ_d_ respectively. In the planktonic phase, we also detected a growth lag, t_lag_. Both binding and growth in the planktonic and the suface-associated population occur up to the maximum carrying capacity of N_p,max_ and N_d,max_ respectively. Similarly, as more and more variants bind and grow on the disc and the disc approached saturation, less and less space is available to bind, thereby decreasing the binding constant.

The model had the following parameters: N_p,0,_ k_b_, µ_p_, µ_d,_ t_lag_, N_p,max_ and N_d,max_. We experimentally determined all parameters, except for kb (**Supplementary note 4** & **Supplementary Table 1**). So, we ran our simulation over a range of kb from 10^-9^ to 10^-5^ over 10 discs. For each simulation, we recorded:

i. the number of unique barcodes that successfully colonize each disc (a measure of the binding bottleneck),
ii. the number of binding events per disc, and
iii. the population structure at the end of 24 h i.e., the frequency of each barcode at the end of the experiment in the planktonic phase and on each of the 10 discs.

#### Temporal model with migration (Supplementary Figure 25)

We used a stochastic simulation model to investigate the temporal evolution of barcoded *Escherichia coli* lineages during repeated cycles of biofilm growth. The simulation code is provided as Supplementary Code C. The model explicitly tracks the abundance of each barcode in two connected compartments: a planktonic population and multiple surface-associated biofilm discs. In addition to cell growth and attachment, the model incorporates migration from the planktonic phase to the biofilm, allowing exchange between the two compartments over successive transfer cycles.

The simulation was initialized using the experimentally determined barcode frequencies at the end of initial binding. To account for stochastic sampling during inoculation, the initial biofilm population on each disc was generated by multinomial sampling from the experimentally observed lineage frequencies. At the beginning of each transfer cycle, the total biofilm population was used to determine the composition of the planktonic inoculum. We calculated that 10% of the total biofilm population was released into the planktonic phase (**Supplementary Figure 5**), and the released cells were sampled by multinomial sampling according to the relative abundance of each lineage in the biofilm. Each biofilm disc was independently inoculated with N_D,0_=5.4*10^5^ cells sampled from the lineage composition of the corresponding disc.

Each transfer cycle simulated 24 h of growth using discrete time steps of 10 minutes (144 time steps per cycle). During each time step, planktonic cells underwent logistic growth with specific growth rate *µp* and carrying capacity N_p,max_ cells. Independently, cells on each biofilm disc underwent logistic growth with specific growth rate *µ_d_* and carrying capacity N_d,max_ cells per disc. The parameters used are listed in **Supplementary Table 1**, estimated in the previous section. Growth events were modeled as Poisson processes, with the expected number of new cells proportional to the current population size, growth rate, time step, and remaining carrying capacity.

In parallel with growth, planktonic cells attached to each biofilm disc with the determined optimum per-cell binding probability *K_b_* =10^-8.095^. Attachment events were modeled as Poisson processes and scaled according to the remaining capacity of each disc, thereby reducing attachment as the biofilm approached saturation. In addition, planktonic cells migrated to each biofilm disc with a per-cell migration probability (m), which was varied across simulations. Migration events were also modeled as Poisson processes and were similarly constrained by the available capacity of each biofilm disc. Cells that attached to the biofilm were removed from the planktonic compartment before the next simulation step.

The simulation consisted of 15 consecutive transfer cycles. To mimic the experimental serial transfer protocol, the number of biofilm discs was reduced by three after every third cycle, resulting in a progressive decrease in the number of available colonization sites throughout the experiment. At the beginning of each new cycle, both the planktonic inoculum and the biofilm inoculum for each remaining disc were resampled from the populations obtained at the end of the previous cycle, thereby propagating stochastic lineage dynamics across transfers.

For each simulation, we recorded the lineage composition of both the planktonic population and the biofilm after every transfer cycle. To facilitate comparison with the experimental sequencing data, simulated populations were down sampled to 250,000 reads by multinomial sampling. Population diversity was quantified using Shannon entropy, and changes in lineage composition relative to the initial population were quantified using the Bray–Curtis dissimilarity. Simulations were repeated for multiple migration rates and independent stochastic replicates to characterize the effect of migration on biofilm population dynamics.

#### Temporal model with antibiotic treatment (Supplementary Figure 26)

The temporal model with antibiotic treatment was the same as the model without antibiotic treatment. However, we introduced an antibiotic bottleneck at cycles 1, 4, 7, 10, and 13 of the model (**Supplementary Figure 26)**. Due to the treatment, only 18,900 cells survived on the discs and 173,000 cells survived in the planktonic fraction. The surviving fraction was estimated as a multinomial sample of the disc and the planktonic population respectively.

### Identifying mutations after genome sequencing

To identify genetic variants, genomic DNA from isolated clones was sequenced and analyzed using Breseq (v0.38.1) ^9^. Bioinformatics processing was performed using default parameters to map raw sequencing reads against the reference genome of *Escherichia coli* LF82 with accession number CU651637.

### Quantifying biofilm formation capacity, fitness, and MIC of mutants

#### Crystal Violet Assay

Overnight cultures of LF82 clones grown in LB at 37°C were diluted to OD600 of 0.05. Five mL of the diluted culture was inoculated in a well containing 24 silicone coupons and incubated for 24 h at 37°C under static condition. Then, the exhausted medium containing floating cells was removed, and the well was washed with 5mL of LB medium twice. After collecting 6 coupons for CFU counts as described above, 6 mL of 1% crystal violet (CV) solution (Sigma-Aldrich) was added to the well and incubated for 15 min at room temperature. The well stained with CV was gently washed with 7 mL of PBS three times to remove excess CV, and the plate was dried up for 1 day in a chemical hood. On the next day, the remaining 18 coupons were collected in 6 microtubes (i.e., 3 coupons/microtube), in which 810 μL of the mixed solution of ethanol/acetone at 80%:20% ratio was added. Coupons were suspended in the solution for 15 min under 400 rpm agitation to allow the CV stain to be dissolved. After transferring 100 μL of dissolved CV solution in a 96-well plate, the OD570 value was measured to quantify the CV solution using a multimode plate reader (Tecan Infinite M200 PRO). As the background control, 6 coupons sheeted in a well containing 5mL of LB medium without bacteria were also treated similarly.

#### Competition for 24h biofilm formation

We pooled the variants FimH W124R, FimH Δ36 coding (86-121/903 nt), Fim H V77D, Crp A145V, BcsG H295Y, BcsG W80Y, and four non-mutated wild-type barcoded clones and performed competition assays. The variant pool was then diluted to OD_600_ = 0.05 and used to inoculate a well sheeted with 18 silicon discs (5mL per well). After 24 hours of static incubation, the planktonic fraction was sampled and subjected to serial dilution and plating for CFU count, as well as boiling for further amplicon sequencing. The well was then washed twice with LB. Three discs were sampled from it and put in an Eppendorf tube containing 500µL of PBS (one disc per tube) and processed for CFU count as described above. A 250µL sample was boiled for PCR amplification of the barcode region and further amplicon sequencing.

#### Competition for invasion of a pre-existing biofilm

A sample well with 18 silicon discs (5mL per well) was inoculated with the resident strain i.e., the strain used to form the biofilm. After 24 hours of static incubation, the well was then washed twice with LB and the variant pool, diluted to OD_600_ = 0.05, was used to inoculate the well. After 30 minutes of incubation allowing the variant pool to invade the pre-existing biofilms, the supernatant and discs were sampled as described in the previous competition assay.

#### Estimating fitness/enrichment score

Barcodes from the competition experiment were amplified and sequenced using the protocol described for the lineage tracking experiment. Variant fitness was estimated from changes in barcode frequency relative to wild-type reference strains. For each sample, raw barcode counts were first converted to relative frequencies by dividing each barcode count (after adding a pseudocount of 0.5 to avoid division by zero) by the total number of reads in that sample.

Enrichment for each variant was then calculated relative to four independently barcoded wild-type (*E. coli* LF82) reference strains. To account for potential batch effects across experiments, one of two wild-type reference pools (LF82-1 or LF82-22) was used depending on the sample index, with samples collected earlier in the experiment referenced to LF82-1 and later samples to LF82-22. For each variant and each wild-type reference barcode, enrichment was calculated as the natural logarithm of the ratio:

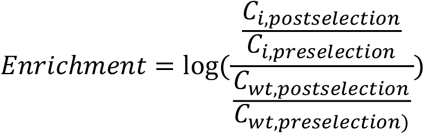

where C denotes the normalized barcode frequency. This formulation measures the change in relative frequency of a variant compared to wild-type over the course of the experiment.

Each variant therefore yielded four independent enrichment estimates per sample, corresponding to the four wild-type reference barcodes. These enrichment values were used as measures of relative fitness in downstream analyses.

## Code and data availability

The sequencing data can be accessed online at the ENA database under the entry with the accession number PRJEB113777. The sample metadata can be found in the excel “Sample_metadata.xlsx”. The code is currently available on github and can be accessed via the link: https://github.com/Alaksh/Biofilm_lineage_tracking.git. Processed data files of large size are availaible on Zenodo and can be accessed using the DOI: **10.5281/zenodo.22301322.**

