## Supplementary figures, tables, and notes for "High-resolution lineage tracking maps the multi-phase evolution and adaptive potential of biofilms"

### Index:

**Supplementary figures 1-19** **2-21**

**Supplementary note 1** **22**

**Supplementary note 2** **23-24**

**Supplementary note 3** **25-26**

**Supplementary note 4** **27-29**

**Supplementary Table 1:** Estimated parameters for the binding-growth model **30**

**Supplementary Table 2:** Sample wise read counts **31**

**Supplementary Table 3:** Mutations identified in individual lineages at the end of the biofilm experiment **32-35**

**Supplementary Table 4:** Mutations in individual lineages at the end of the well-mixed shake-flask experiment **36-38**

**Supplementary Table 5:** Mutations in individual lineages at the end of the biofilm evolution with antibiotic treatment **39-47**

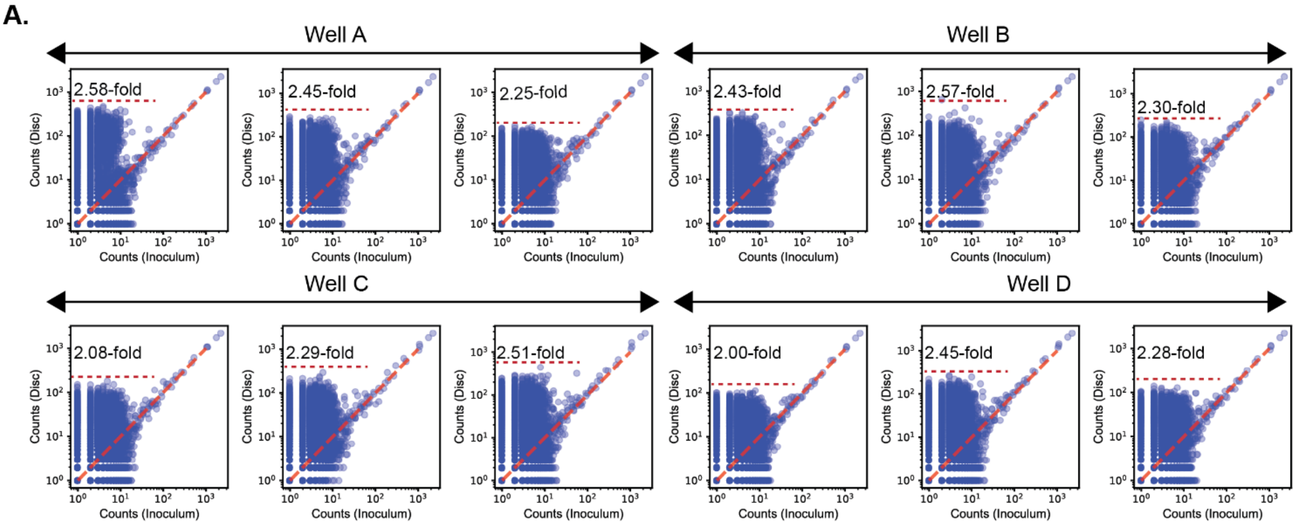


**Supplementary Figure 1:**

A. Comparison of barcode counts between the inoculum and a representative disc-associated biofilm. The maximum enrichment i.e., $log_{10}$-fold change in frequency compared to the inoculum observed for a barcoded variant in written in text.

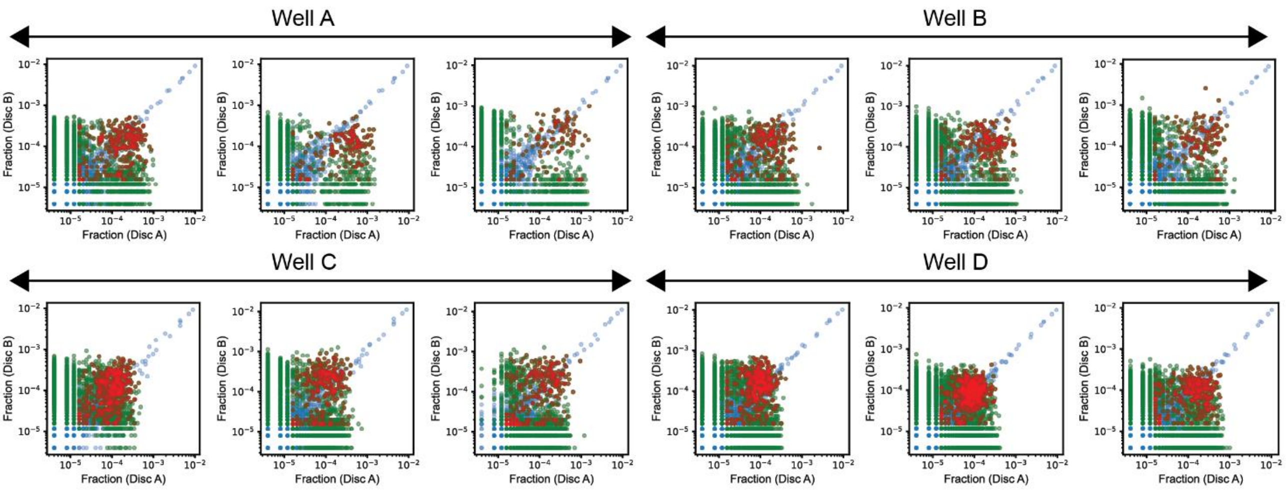


**Supplementary Figure 2:** Comparison of barcode counts between the two discs from the same well for three replicates each of all replicate wells. Variants without any significant enrichment are blue, significantly enriched on one disc are green, and variants significantly enriched on both discs are red. Variants were classified as significantly enriched if their enrichment exceeded the mean planktonic enrichment plus 1.96 standard deviations of the threshold = 0.623, representing enrichment due to noise in sampling.

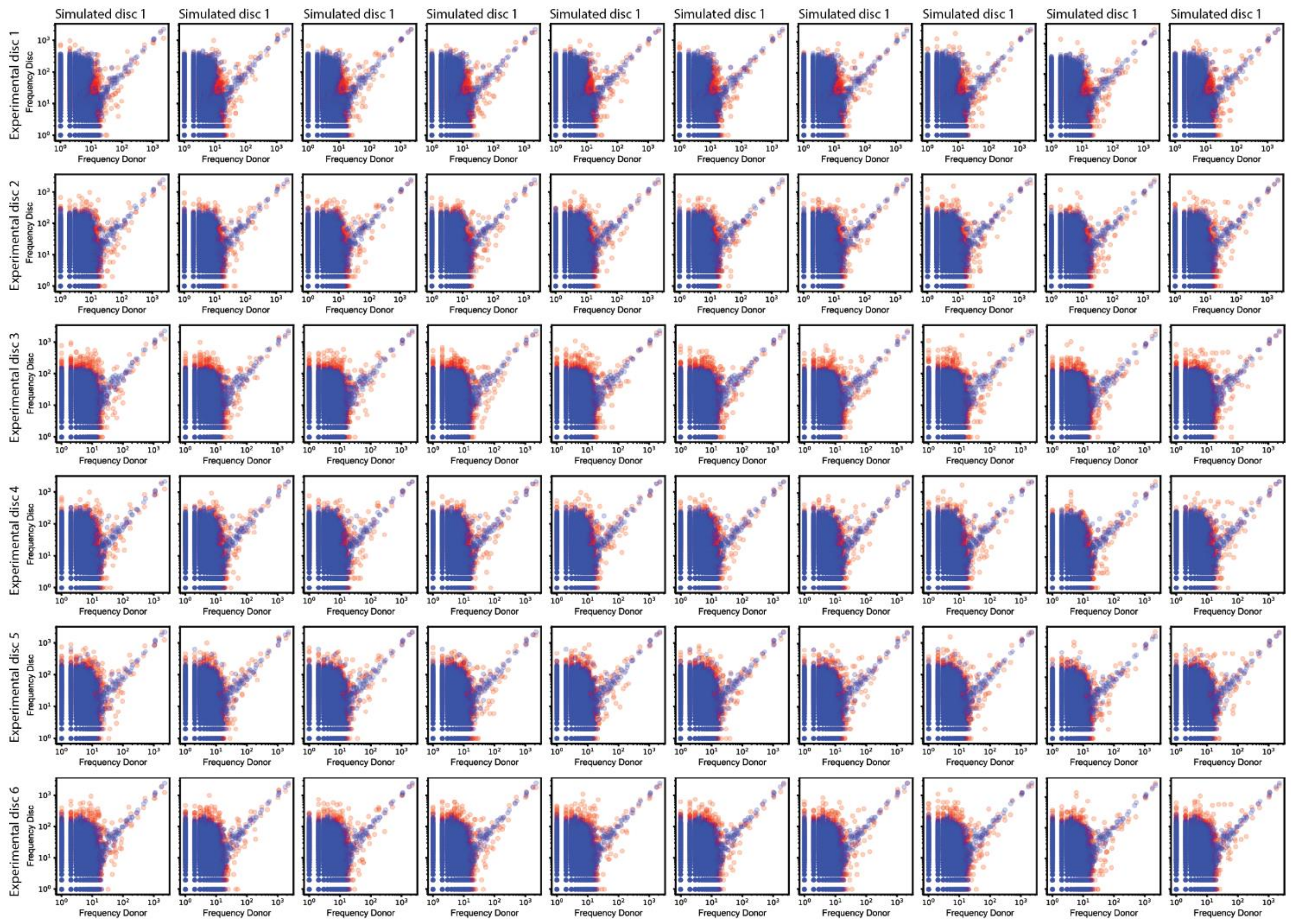


**Supplementary Figure 3:** Comparison of barcode counts between the inoculum and a representative experimental disc-associated biofilm from (blue) for all replicates (3 discs each from four replicate wells), and between the inoculum and a representative disc from all simulations using the optimum $K_d$ value (red).

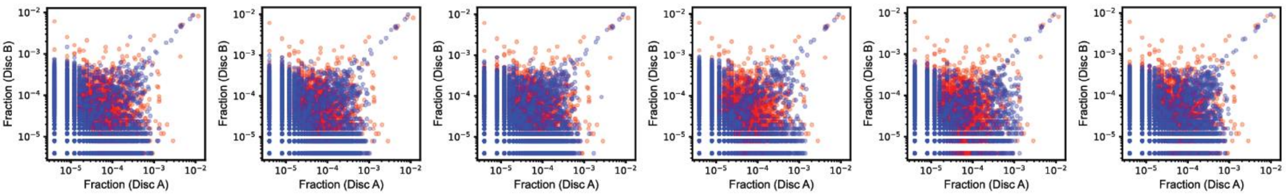


**Supplementary Figure 4:** Pairwise comparison of barcode frequencies between two discs recovered from the same experimental well and between two discs recovered from simulations at the optimum $K_d$.

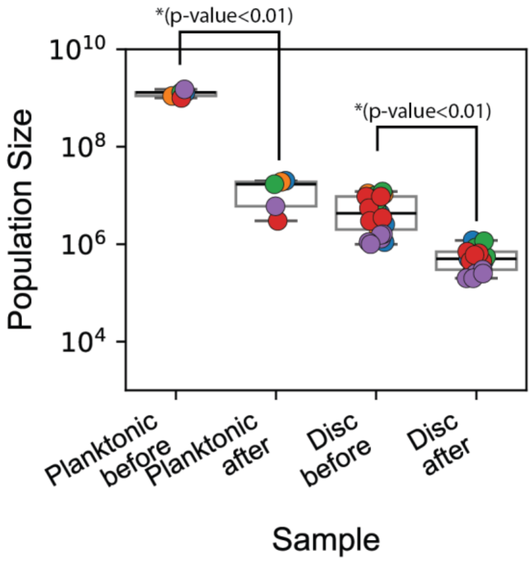


**Supplementary Figure 5: Impact of media replacement on population size.** The population size measured as Colony forming units (CFUs) in the biofilm-associated planktonic and surface-attached biofilm phase before and after replacement with fresh media. Significance estimated using a student's t-test. Box plots show the median (center line), interquartile range (box), and whiskers extending to 1.5× the interquartile range for n = 4 for planktonic and n = 3 discs x 4 wells for disc-attached biofilms.

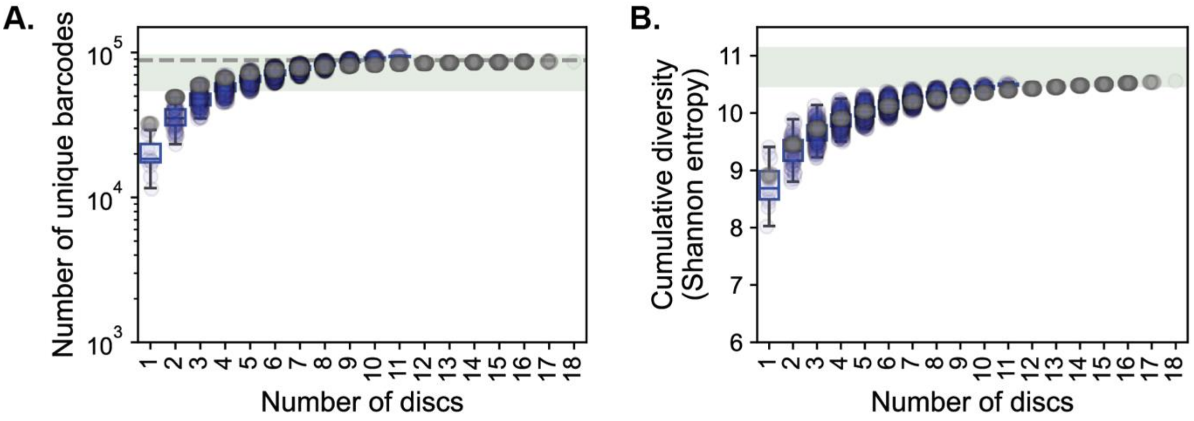


**Supplementary Figure 6: Cumulative diversity in randomly subsampled discs.** We used an iterative subsampling simulation to determine the minimum number of discs required to represent the total initial population diversity. We subsampled combinations of 1-12 discs from our experimental and simulated datasets with a population size of a disc's maximum carrying capacity. We then determined the population composition in the subsampled discs and a sum of the variant fraction on multiple samples discs provided the population composition of the overall biofilm.

A. Number of unique barcodes for different numbers (1-18) of randomly subsampled discs after initial biofilm formation in the experiment (blue) and simulation (grey). Shaded pale green regions indicate the mean ± 1.96 standard deviations of number of unique barcodes in four independent wells at t = 0, representing the initial diversity in the absence of a bottleneck.

B. Cumulative population diversity measured as Shannon entropy for different numbers (1-18) of randomly subsampled discs after initial biofilm formation in the experiment (blue) and simulation (grey). Shaded pale green regions indicate the mean ± 1.96 standard deviations of Shannon entropy measured across four independent wells at t = 0, representing the initial diversity in the absence of a bottleneck.

Box plots show the median, interquartile range, and whiskers extending to 1.5× the interquartile range. We found that as few as 3 discs captured all barcodes in the library and the cumulative diversity of up to seven discs showed no significant divergence from the initial population represented in the shaded pale green region($p > 0.05$).

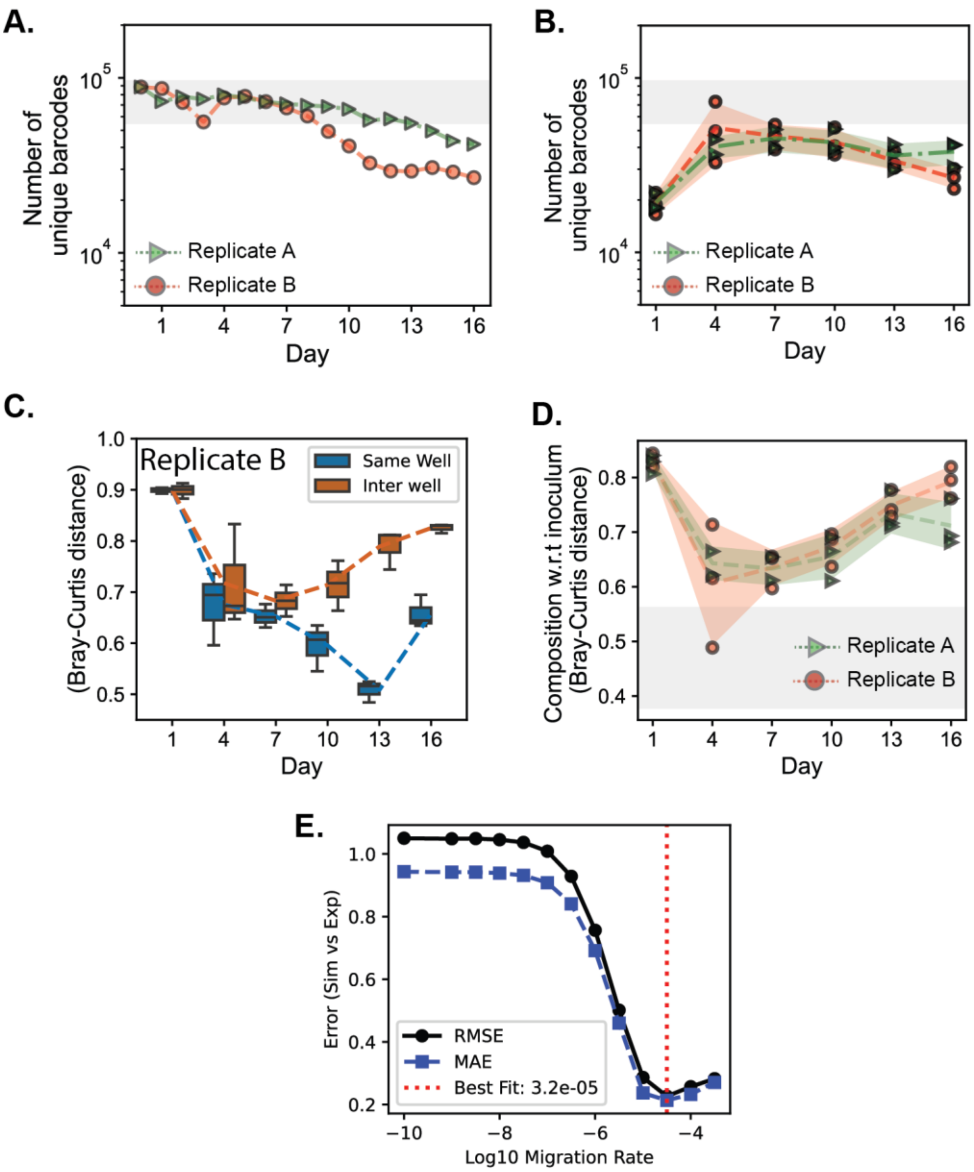


**Supplementary figure 7:**

**A-B.** The change in the number of unique barcodes in the biofilm-associated planktonic population (left) and three discs each (right) from each replicate over time (days). (Left) The dashed lined and the shaded area in both curves represent the mean and the standard deviation of the 3 individually sampled discs at each time point. Shaded grey regions indicate the 99% confidence interval (mean ± 2.52*standard deviations) of Shannon entropy measured across four independent wells at $t = 0$ in the absence of a bottleneck.

**C.** Bray–Curtis distances between disc-associated populations over time for discs recovered from the same well (blue) and from different wells (orange) of replicate A.

**D.** Compositional dissimilarity between the inoculum and populations recovered from the disc-associated biofilms for replicate A (green) and replicate B (red) over time, quantified using Bray–Curtis distance. The dashed line represents the mean population compositional dissimilarity to the inoculum and the shaded region represents mean ± standard deviations. Shaded grey regions indicate the 99% confidence interval (mean ± 2.52*standard deviations) of the Bray-Curtis distance from the inoculum measured across four independent wells at $t = 0$ in the absence of a bottleneck.

**E.** Evaluation of model performance using Root Mean Square Error (RMSE; solid line) and Mean Absolute Error (MAE; dashed line) between simulated and experimental Shannon entropy across Cycles 1, 4, and 7. The optimal secondary migration rate was identified at the minimum RMSE.

In panels B and D, points represent measurements from individual discs (three discs per well), and lines indicate the mean across replicate wells. Box plots show the median, interquartile range, and whiskers extending to 1.5× the interquartile range.

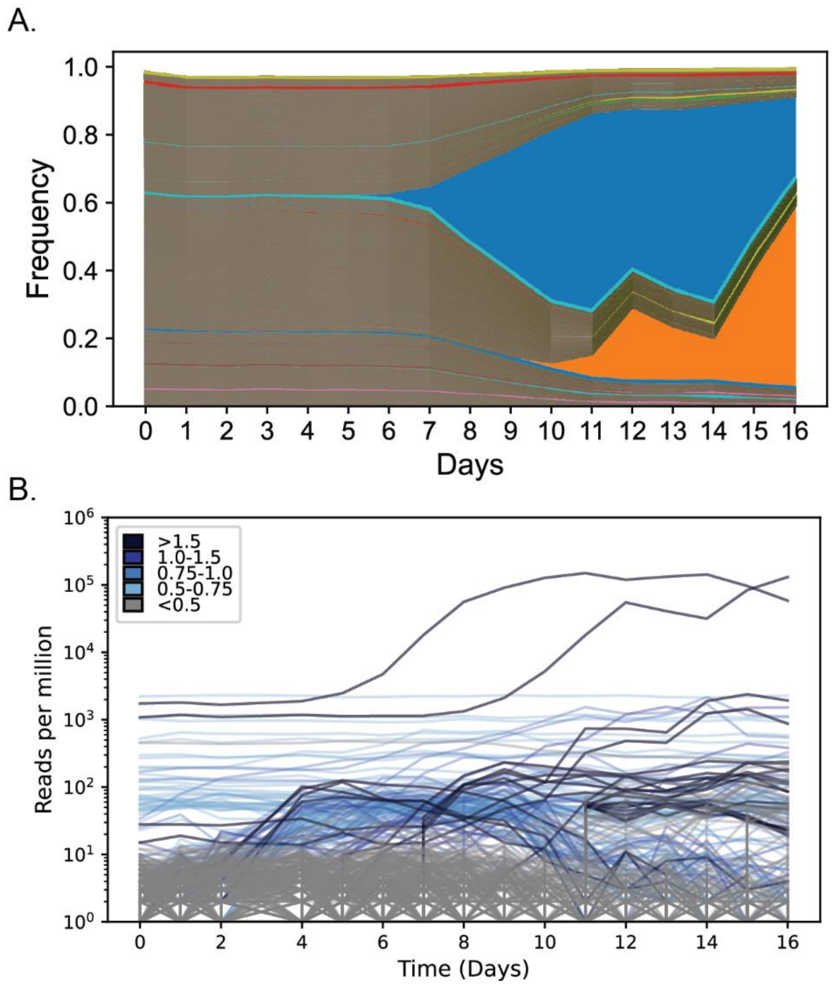


**Supplementary Figure 8: Adaptation dynamics in replicate B.**

A. Muller plot showing the temporal dynamics of barcoded variants in biofilm-associated planktonic population in replicate B, where each colored region represents the relative frequency of a single variant over time.

B. Temporal trajectories of individual variants classified as beneficial (blue; inferred fitness > 0) or neutral (grey). Line color intensity increases with inferred fitness. Fitness inference is described in the Methods.

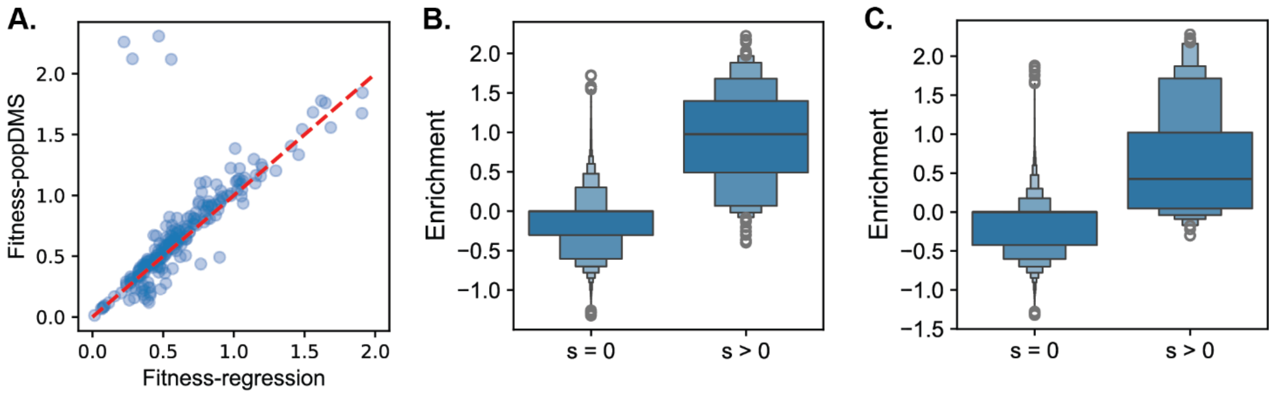


**Supplementary Figure 9:**

A. Correlation between fitness estimated for lineages in the biofilm-associated planktonic population using regression (x-axis) and the pop-DMS algorithm (y-axis). Details of the fitness measurements can be found in Supplementary methods section 4.6.

B. Comparing the distribution of enrichment for variants with inferred fitness s = 0 versus those with inferred fitness > 0 for replicate A.

C. Comparing the distribution of enrichment for variants with inferred fitness s = 0 versus those with inferred fitness > 0 for replicate B.

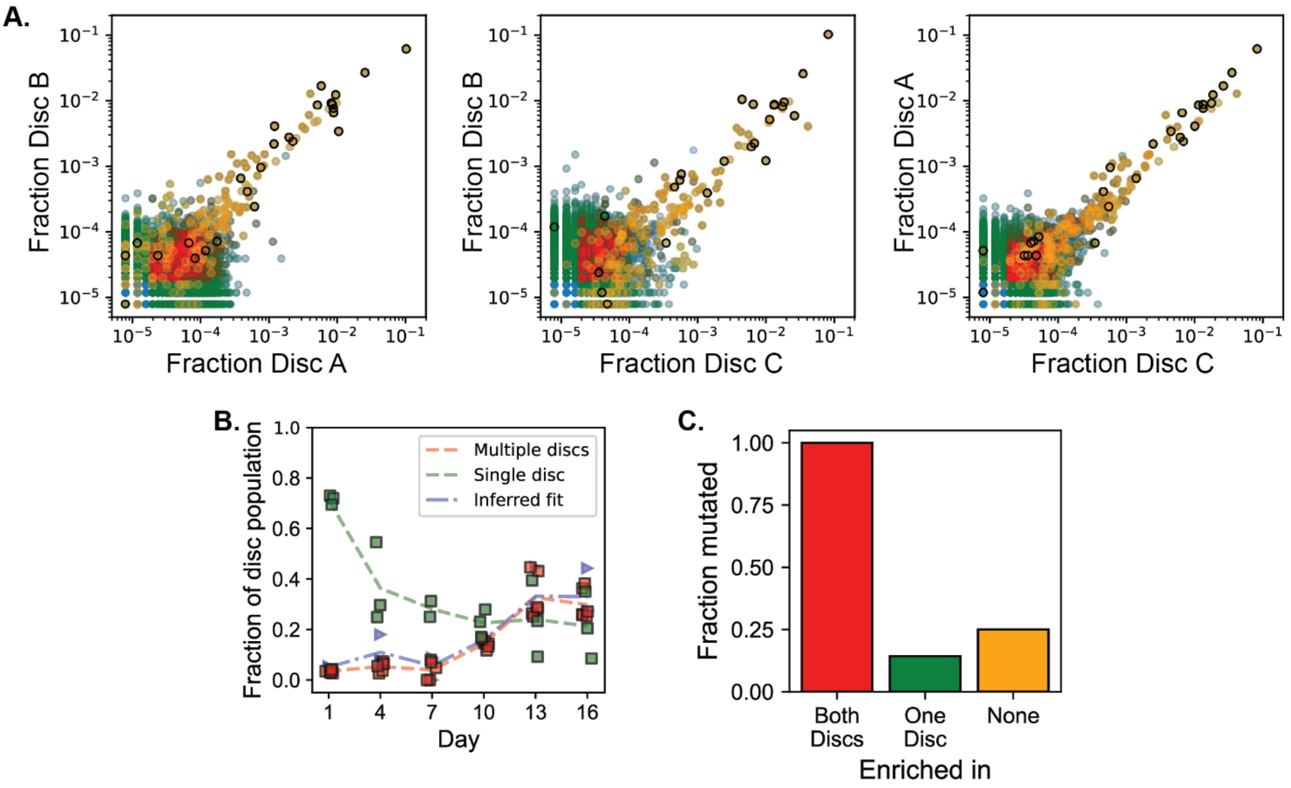


**Supplementary Figure 10: Mutants enriched in different biofilm and planktonic phases**

A. Comparison of barcode counts between the two discs from the same well for three replicate disc combinations. Variants without any significant enrichment are blue, significantly enriched on one disc are green, and variants significantly enriched on both discs are red. Variants with inferred fitness > 0 are represented in orange. Sequenced variants with mutations are highlighted with the black edge. Variants were classified as significantly enriched if their enrichment exceeded the mean planktonic enrichment plus 1.96 standard deviations of the threshold = 0.623, representing enrichment due to noise in sampling.

B. The change in the fraction of population represented by enriched lineages shared between multiple discs (red curves representing sum of red variants in panel A over time), enriched lineages on a single disc (green squares representing sum of green variants in panel A over time) and population fraction with fitness > 0 (blue triangles representing orange variants in panel A over time).

C. We sampled variants that were shared (significantly enriched) on multiple discs (Red), in one disc only (green), and neither discs (yellow) and sequenced their genomes to calculate the fraction of variants mutated for each category. We show all discs for replicate A, as it was the only replicate where the genomes were sequenced.

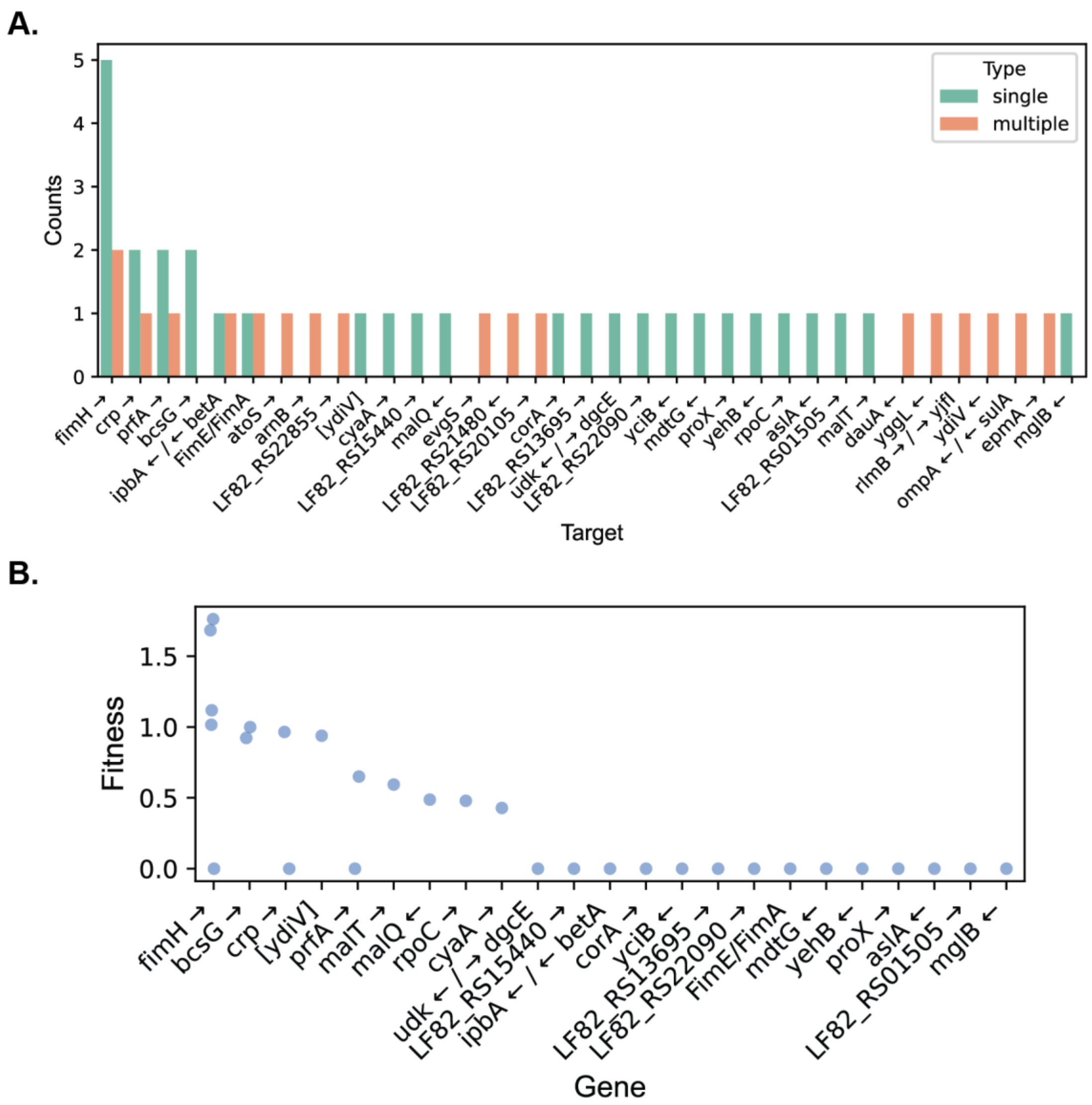


**Supplementary Figure 11: Genetic targets and fitness effects of adaptive mutations**

A. At the end of the experiment we sequenced 106 lineages with unique barcodes for replicate A (with 19 variants recovered from the biofilm-associated planktonic and 87 lineages recovered from the disc). Amongst the 40 mutated lineages, panel A shows the Number of independently arising mutations per genetic target among barcoded variants. Green bars indicate clones carrying a single mutation in the indicated gene, whereas orange bars indicate clones carrying additional mutations in other loci.

B. Inferred fitness scores for mutations occurring as single mutations, as indicated in panel A. Mutations detected in several other genes showed no clear evidence of adaptation: they either appeared without measurable positive fitness effects or occurred only in combinations with other mutations that showed the actual convergence and/or positive selection

Variants in which mutations arose independently multiple times in the same target, or that exhibited positive inferred fitness, were classified as adaptive.

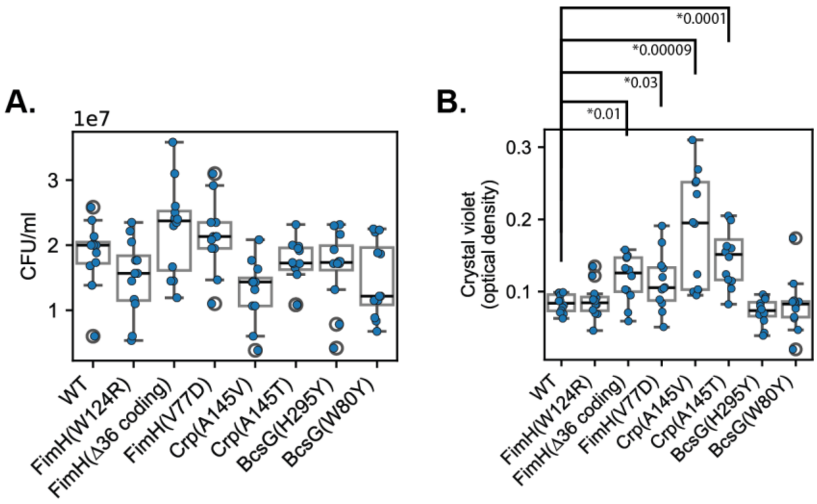


**Supplementary Figure 12: Raw data for Crystal Violet assay. We measured biofilm formation on silicone discs using the crystal violet assay. Crystal violet stains reflect the total biomass in biofilms.**

A. CFU enumeration after 24 h of biofilm formation for each variant.

B. Crystal violet OD after 24 h of biofilm formation for each variant.

Each point represents an independent biological replicate (n = 12). Asterisks indicate statistically significant differences compared with wild type (two-sided Mann–Whitney U test with Benjamini–Hochberg correction for multiple testing).

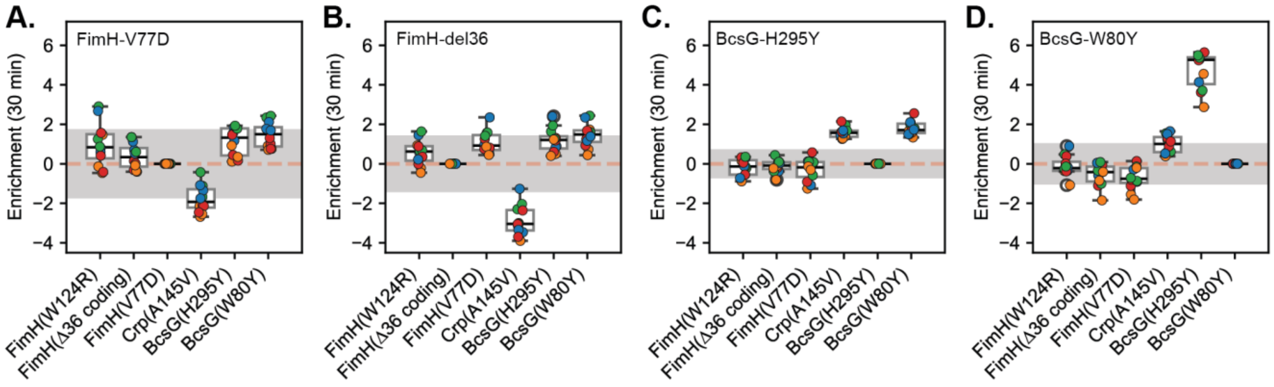


**Supplementary Figure 13: Fitness during invasion into discs pre-colonized with different variants**

**A,B,C,D:** Enrichment scores during invasion into discs pre-colonized by the FimH Δ36, FimH V77D, BcsG H295Y, and BcsG W80Y variants respectively. Enrichment scores were calculated as the log change in frequency of each variant relative to four independently barcoded neutral wild-type lineages between the beginning and end of each experiment. Each point represents an independent measurement from three replicate wells against four reference barcodes. Box plots show the median, interquartile range, and whiskers extending to 1.5× the interquartile range (n = 3 × 4).

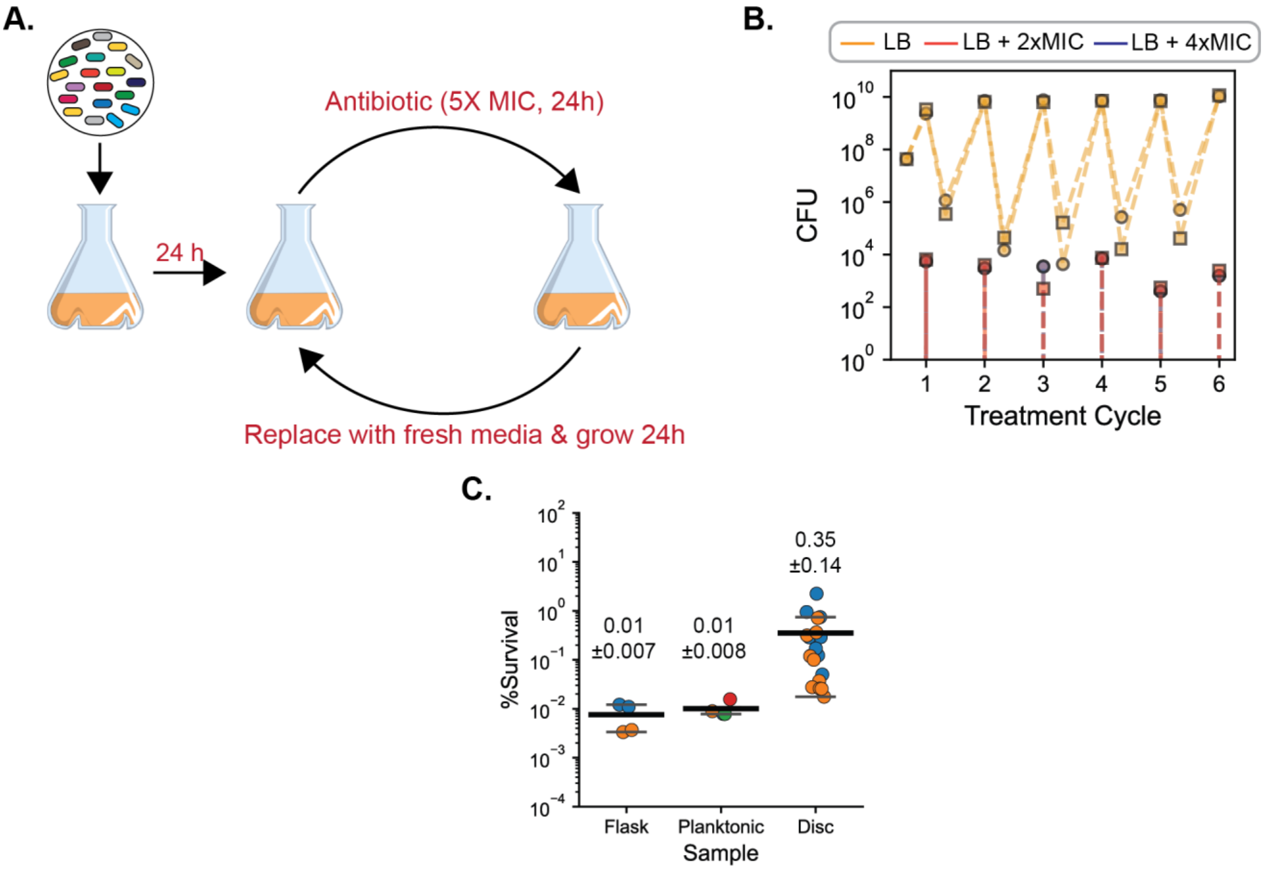


**Supplementary Figure 14: Antibiotic treatment of well-mixed cultures in shake flasks:**

**(A)** Schematic of the experimental design incorporating periodic antibiotic treatment between media replacement and growth cycles in well-mixed shaken-flasks.

**(B)** Colony-forming units (CFUs) measured over five antibiotic treatment cycles for populations grown in flasks without biofilms. CFUs were quantified following growth in LB alone (yellow), LB supplemented with 2× MIC (red), or 4× MIC (blue). Trajectories with squares and circles distinguish the two biological replicates.

**(C)** Percentage survival in well-shaken control flasks, the biofilm-associated planktonic fraction and three discs after a cycle of treatment with 5X MIC of Amikacin for 24 hours. Circles represent individual measurements and different color represent independent biological replicates.

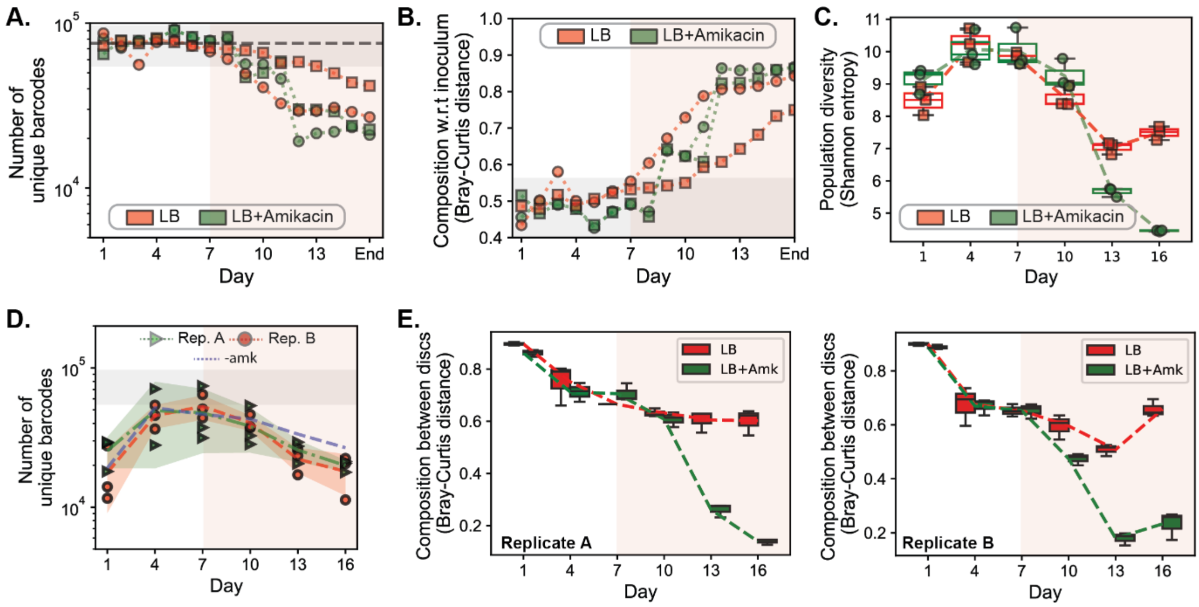


**Supplementary Figure 15: Temporal dynamics upon antibiotic treatment**

**(A)** The change in the number of unique barcodes in the biofilm-associated planktonic population of the experiment without amikacin (red) and with amikacin treatment (green). Shaded grey region indicates the 99% confidence interval (mean ± 2.52*standard deviations) of Shannon entropy measured across four independent wells at $t = 0$ in the absence of a bottleneck.

**(B)** Compositional dissimilarity between the inoculum and populations recovered from the biofilm-associated planktonic phase of the experiment without amikacin (red) and with amikacin treatment (green). Shaded grey region indicates the 99% confidence interval (mean ± 2.52*standard deviations) of Bray-Curtis distance from the inoculum measured across four independent wells at $t = 0$ in the absence of a bottleneck.

**(C)** Population diversity measured as Shannon entropy for disc-associated biofilm populations sampled from experiments conducted without amikacin (red) and with amikacin (green) for replicate B.

**(D)** The change in the number of unique barcodes in the disc-associated biofilm population sampled from three discs each over time (days) in the experiment with amikacin for replicate A (green triangles) and replicate B (red circles). The green and red dashed lines and shaded area represent the mean and the standard deviation of the 3 individually sampled discs. The blue dashed line represents the mean change in the number of unique barcodes experiment without amikacin. Shaded grey region is the 99% confidence interval of number of unique barcodes detected across four independent wells at $t = 0$ in the absence of a bottleneck.

**(E)** Bray–Curtis distances between disc-associated populations over time for discs recovered from the same well of the biofilm temporal experiment without amikacin (red) and with amikacin (green). Box plots show the median, interquartile range, and whiskers extending to 1.5× the interquartile range.

The vertical red shaded area marks the period where significant increase in resistance frequency was observed.

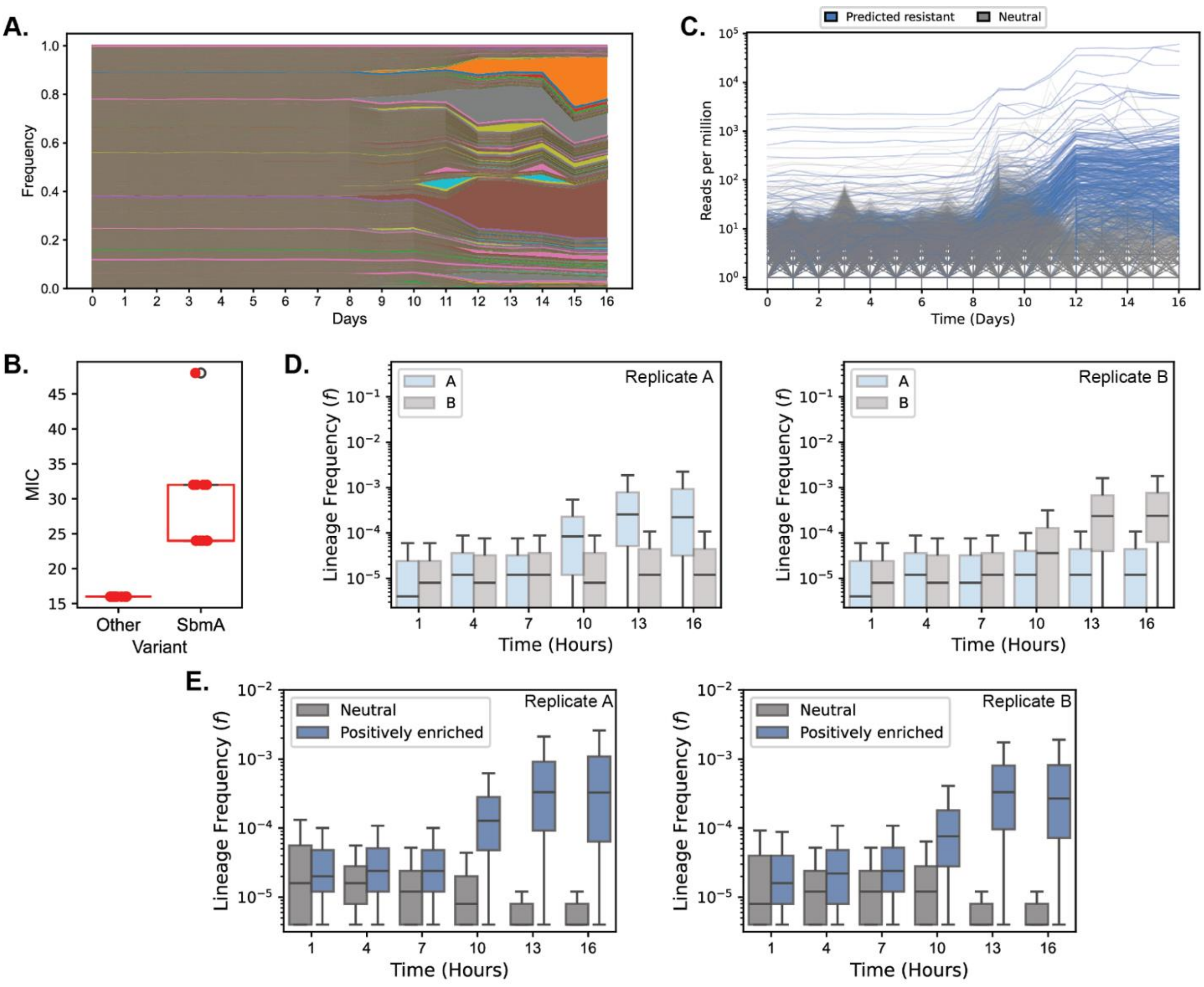


**Supplementary Figure 16: Adaptation dynamics upon antibiotic treatment**

**(A)** Muller plot showing the temporal dynamics of barcoded variants in biofilm-associated planktonic population in replicate B of the temporal biofilm experiment with antibiotic treatment.

**(B)** MIC of 10 variants with *sbmA* mutations and other 10 variants (red circles).

**(C)** Temporal trajectories of individual predicted resistant variants (light blue) or neutral variants (grey). Predicted resistant variants are ones with fold enrichment comparable to the sequenced lineages (p-value > 0.05).

**(D)** Frequency of sequenced predicted resistant lineages in Replicate A (light blue), Replicate B (grey) for significantly enriched variants isolated from replicate A (left) and replicate B (right).

**(E)** Temporal dynamics of neutral and predicted resistant variants in replicate A and replicate B of the biofilm experiment with antibiotic treatment.

Box plots show the median, interquartile range, and whiskers extending to 1.5× the interquartile range.

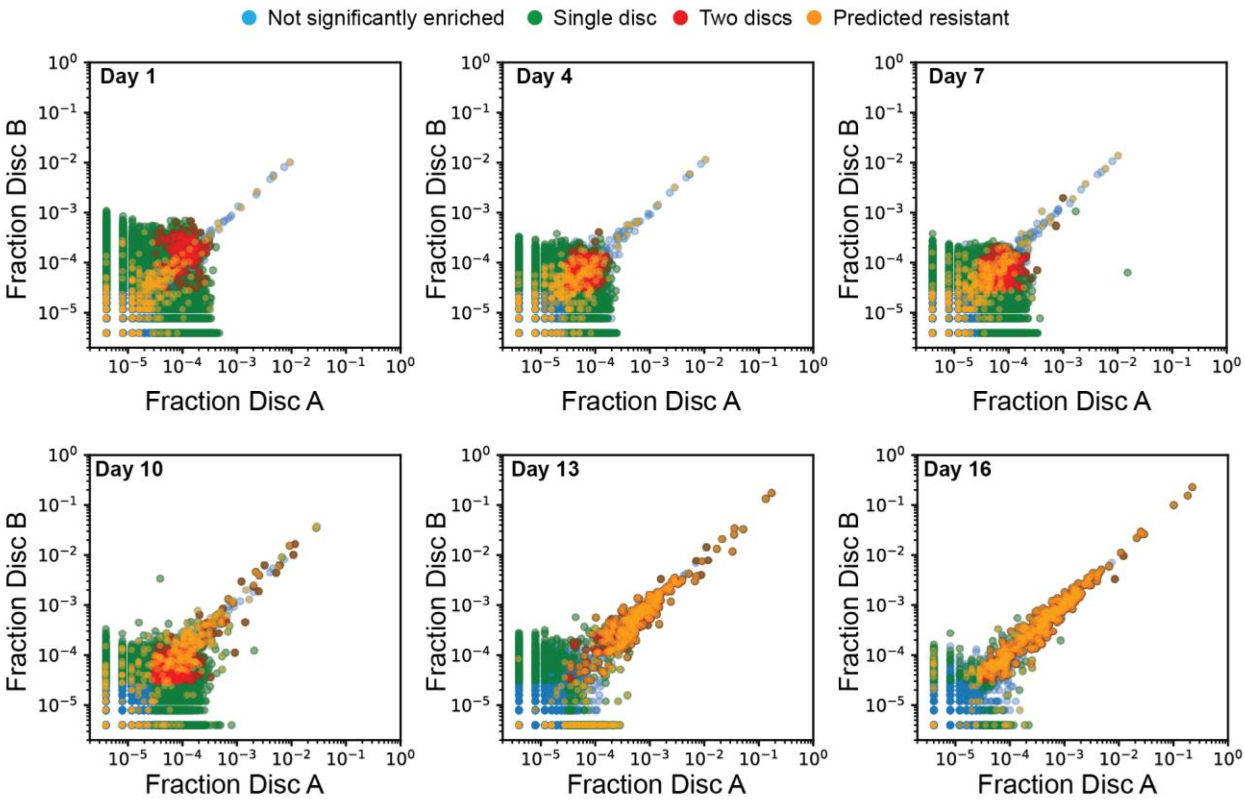


**Supplementary Figure 17:** Pairwise comparison of barcode counts over days between two discs recovered from the same well for replicate B. Variants without any significant enrichment are blue, significantly enriched on one disc are green, variants significantly enriched on both discs are red, and variants with predicted resistance are orange for replicate B.

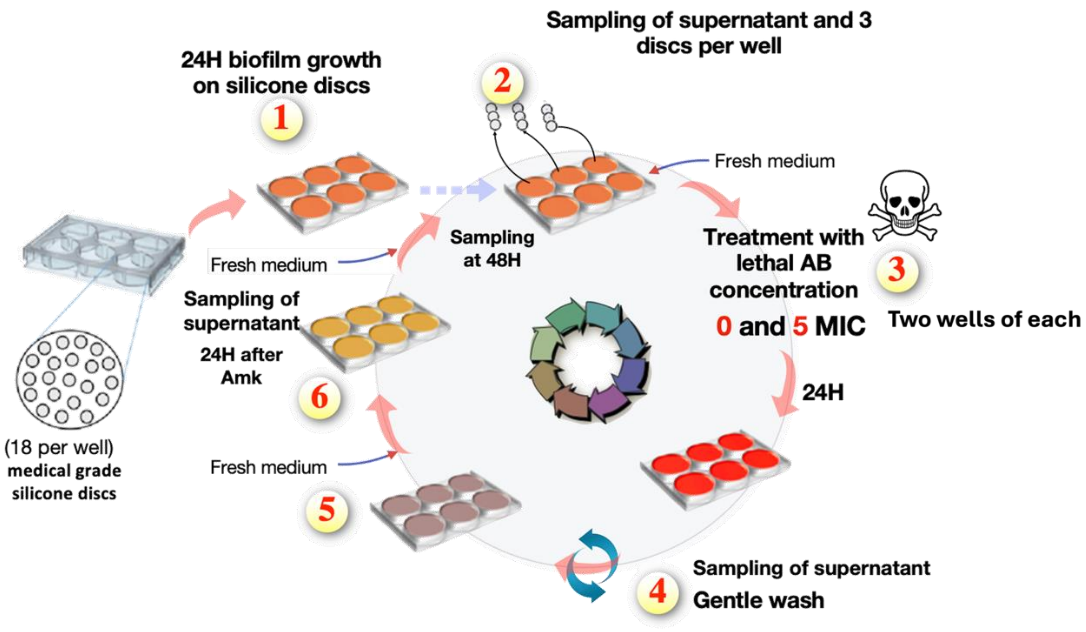


**Supplementary Figure 18:** Evolution experiment with biofilms with and without antibiotic treatment demonstrating an entire cycle of treatment.

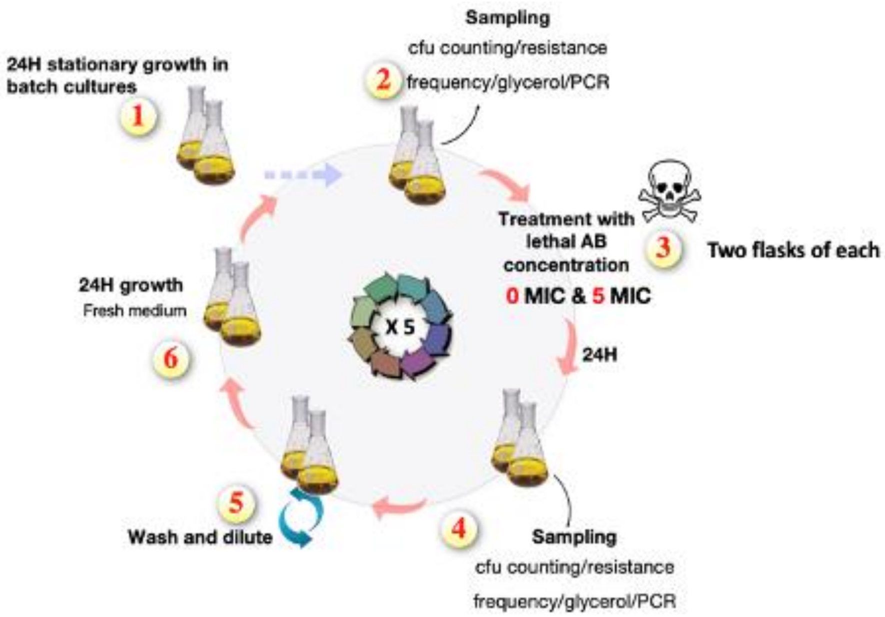


**Supplementary Figure 19:** evolution experiments in well-mixed flask controls.

**Supplementary note 1:**

We determined the resampling threshold by calibrating the normalization depth N against a reproducible baseline of the inoculum. We had re-sequenced four different replicates of the inoculum. We defined a set of high-confidence barcodes (n = 72,495) as ones that were consistently detected across all four inoculum replicates. To identify the optimal N, we performed rarefaction analysis using a pooled inoculum reference (by adding the read counts for all 4 replicates) as the source population. This population was subsampled at varying depths ranging from between 1-$10^7$ over 10 iterations to determine a relationship between sampling depth and barcode richness (**Supplementary Figure 20)**.

Our simulation indicated that a read depth of 200,000 was sufficient to recover 100% of the 72495 high confidence barcodes. We ultimately selected a standardized depth of 250,000 reads per sample. This depth provides a ~114% coverage of the core barcodes and a safety margin to reproducibly recover the population structure. The filtered data with resampled barcodes were stored as a separate filtered file. Thirteen out of 189 samples had read counts significantly lower than 200,000 reads, were excluded from the analysis.

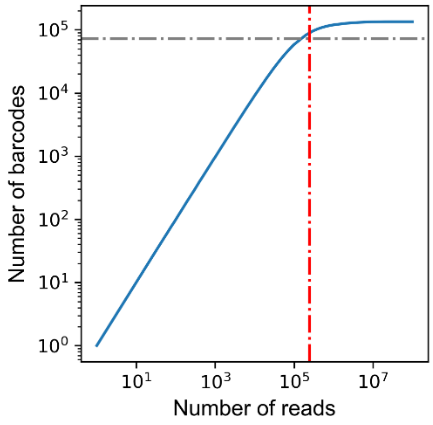


**Supplementary Figure 20: Rarefaction-based Determination of Normalization Depth.** Rarefaction curve showing the mean number of barcodes detected (blue line) as a function of sequencing depth (N) for 10 simulations. The dashed horizontal gray line represents the number of high-confidence barcodes (n = 72,495) identified in the inoculum intersection. The vertical red line indicates the selected normalization depth (N = 250,000), which ensures full recovery of the core library complexity with a protective margin for lower-abundance barcodes.

**Supplementary Note 2**

Supplementary Figure 21A shows an example trajectory for a variant that acquired a beneficial mutation and its log transformation w.r.t the wild type reference. As we observe, the trajectory may not have a beneficial mutation from the onset and an observable increase frequency exponentially begins around day 10. The one on the right represents the trajectory of another variant with a beneficial mutation. For this variant, the trajectory starts to increase exponentially on day 5, but starts to decrease eventually at day 10. The increase in frequency of a variant occurs when its fitness is higher than the mean fitness of the population. With an increase in the frequency of beneficial mutations in the population, the mean fitness of the population increases. If the mean fitness approaches a beneficial variant's fitness or surpasses it, it can eventually plateau or decrease respectively.

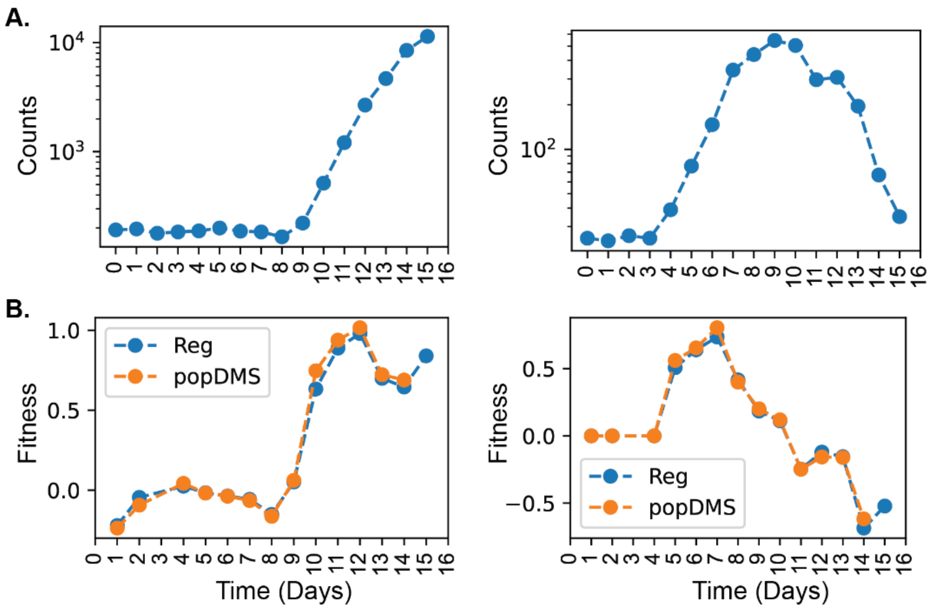


**Supplementary Figure 21: Example trajectories (A)** Example lineage trajectories of beneficial mutations. **(B)** The inferred fitness over time estimated using linear regression and the pop DMS algorithm.

Due to the change in the slope upon fixing of the beneficial mutation and the possible change in the trajectory owing to the mean population fitness, we measured the fitness score over a sliding window of every 3 consecutive time points using linear regression and popDMS (Supplementary Figure 21B). As we can observe, the value changes consistently over time for the fitness determined using each method. So, as the actual fitness, we retained the maximum fitness value.

**Filtering the data:** We estimated that the average error rate associated with sequencing was ~0.1% (or 0.001). Variants with reads less than 0.1% of total reads i.e., (25 in case of 250,000), may be associated with noise. The fitness estimates for such low count barcodes can be spurious merely due to fluctuations in the read counts. Therefore, for each variant, we determined the slope only if the number of reads for all time-points of the window was >= 25.

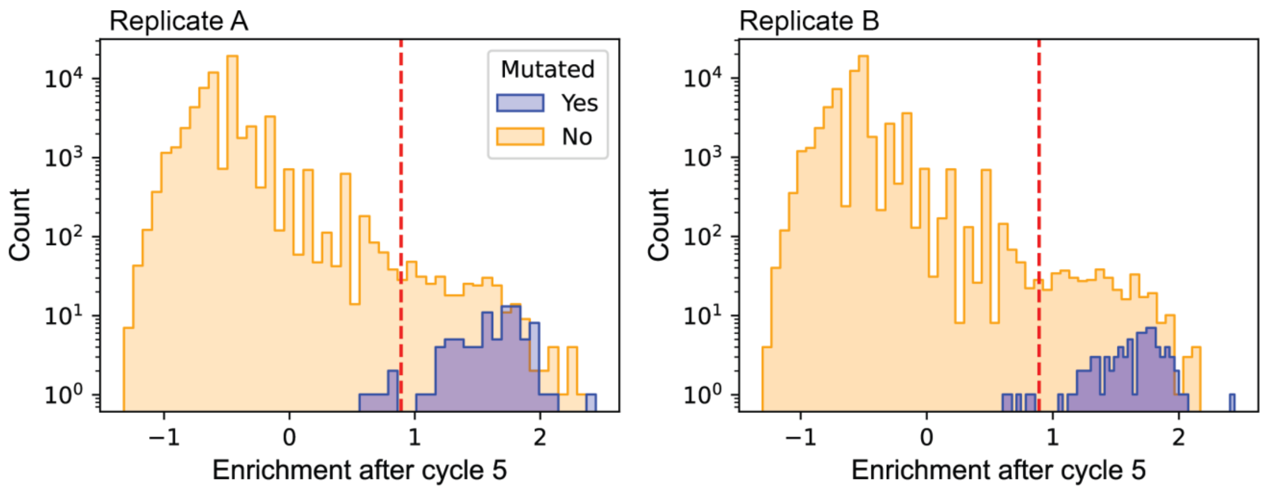


**Supplementary Figure 22:** Enrichment at the end of cycle 5 of antibiotic treatment for all variants (yellow) and sequence-verified resistant variants (blue, n = 87).

**Supplementary Note 3: The simple singular bottleneck does not explain expansion of rare lineages**

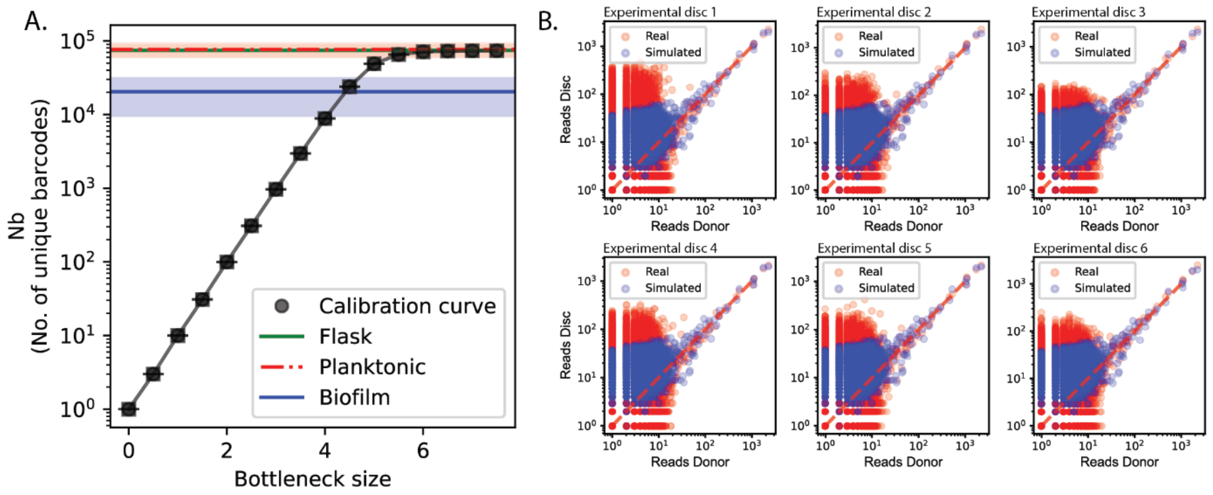


**Supplementary Figure 23: We estimated a simple bottleneck as a multinomial sampling of the inoculum with different population or bottleneck sizes.** (A) The number of unique barcodes detected as a function of the bottleneck size. (B) The correlation between the reads in the donor versus the reads on each disc for 6 simulated discs (blue) and experimental discs (red).

In the simple model, we assume that all cells have experienced a similar identical bottleneck. Therefore, in the simple model we assume that the population on the disc is a random multinomial sample of the donor library. To estimate the bottleneck, we performed simulations using the actual frequency of the donor library.

We randomly sampled the donor population at different bottleneck sizes from 1 cell to $10^9$ cells using random multinomial sampling. The sampling resulted in an array with different numbers of cells assigned to different barcodes with a probability proportional to the frequency in the donor. We repeated sampling for each bottleneck size 10 times. For each bottleneck simulation, we determined the number of unique barcodes. We plotted the number of unique barcodes against the bottleneck size. For bottleneck size < size of the number of unique barcodes in the donor (~$10^5$), there was a linear correlation between the log(number of unique barcodes) and log(bottleneck size), providing us a standard calibration curve linking number of unique barcodes to the bottleneck size (**Supplementary Figure 23A)** .

Using the mean number of unique barcodes (20388), we determined the bottleneck size of 21608 +/- 5233 (**Supplementary Figure 23A**). However, at this bottleneck we did not observe the expansion of rare barcodes as observed in the experiment (**Supplementary Figure 23B)**. However, on the discs, we observed an expansion > 2-orders of magnitude. Therefore, a singular equal bottleneck did not explain the observed results.

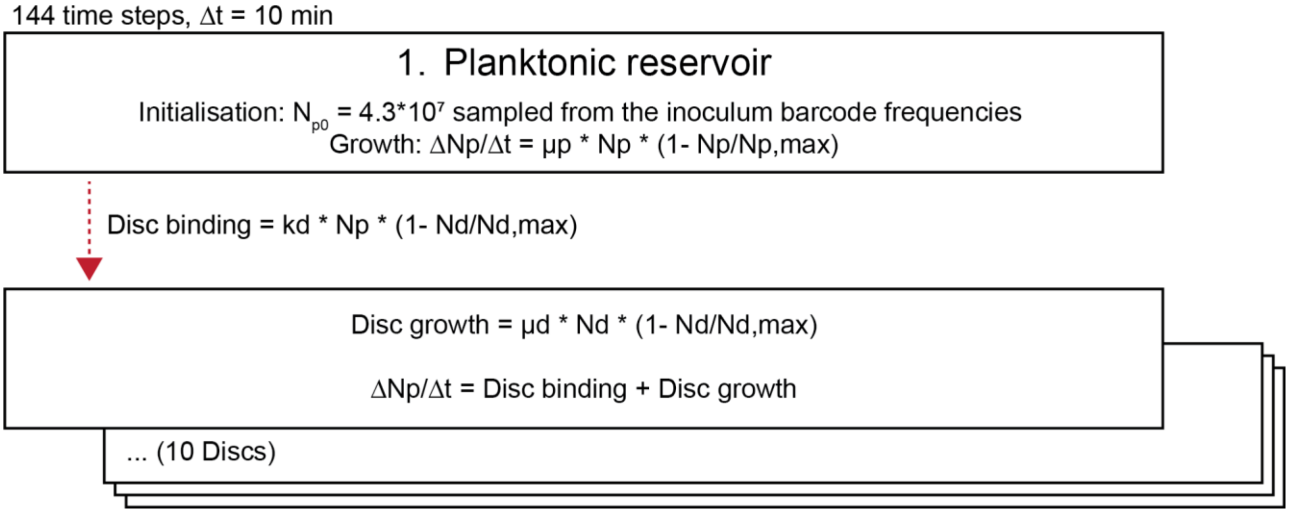


**Supplementary Figure 24:** Binding-growth model algorithm.

**Supplementary Note 4**

**Estimation of maximum carrying capacity in biofilm (disc) and planktonic phases:** Five independent overnight cultures of LF82 WT strain were grown in LB at 37°C and diluted to $OD_{600}$ of 0.05. Five mL of each diluted culture was inoculated in a well containing 10 silicone coupons and incubated for 24 h at 37°C under static condition. The planktonic phase was then sampled from each well, and the biofilms were gently washed twice with 5 mL of fresh LB. Each coupon was sampled and transferred in an eppendorf tube containing 500µL of PBS. Each eppendorf tube containing a disc was then vortexed for 1 minute followed by 10 minutes of sonication to dislodge the biofilm cells from the silicon disc. The tubes, as well as the sampled planktonic phase, were then briefly vortexed before being serially diluted and plated for CFU counting.

**Estimation of the growth rate in the biofilm phase:** Each well of a 6-well plate was sheeted with three silicone discs and inoculated with 5 mL of independent LF82 WT strain overnight cultures normalized to $OD_{600}$ = 0.05. Nine such plates were prepared and incubated statically at 37°C. Every hour, each well of a full plate was washed twice with 5 mL of fresh LB and each disc was sampled and transferred in an Eppendorf tube containing 500µL of PBS followed by 1 minute of vortex and 10 minutes of sonication. The tubes were then briefly vortexed before being serially diluted and plated for CFU counting. The kinetics over 9 hours was used to calculate the growth rate.

**Estimation of the growth rate and lag time in the planktonic phase:** Each well of a 6-well plate was inoculated with 5 mL of independent LF82 WT strain overnight cultures normalized to OD600 = 0.05. Six such plates were prepared and incubated statically at 37°C. Every 30 minutes, the planktonic culture was sampled from each well a full plate, serially diluted and plated for CFU counting. The resulting kinetics over 3 hours was used to calculate the lag time and growth rate.

All estimated parameters are listed in **Supplementary Table 1**.

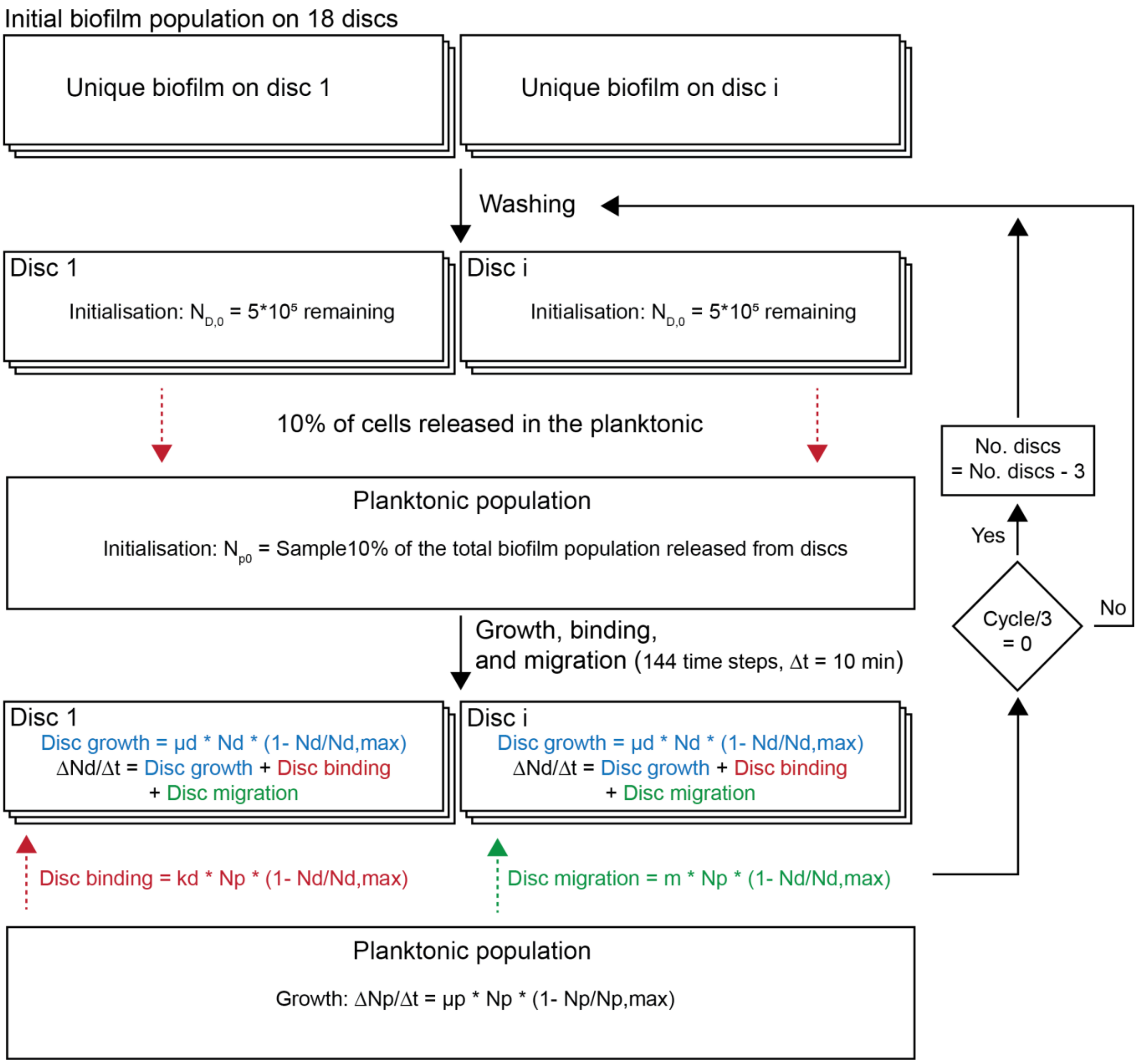


**Supplementary Figure 25:** Algorithm for population temporal evolution by binding, growth, and migration.

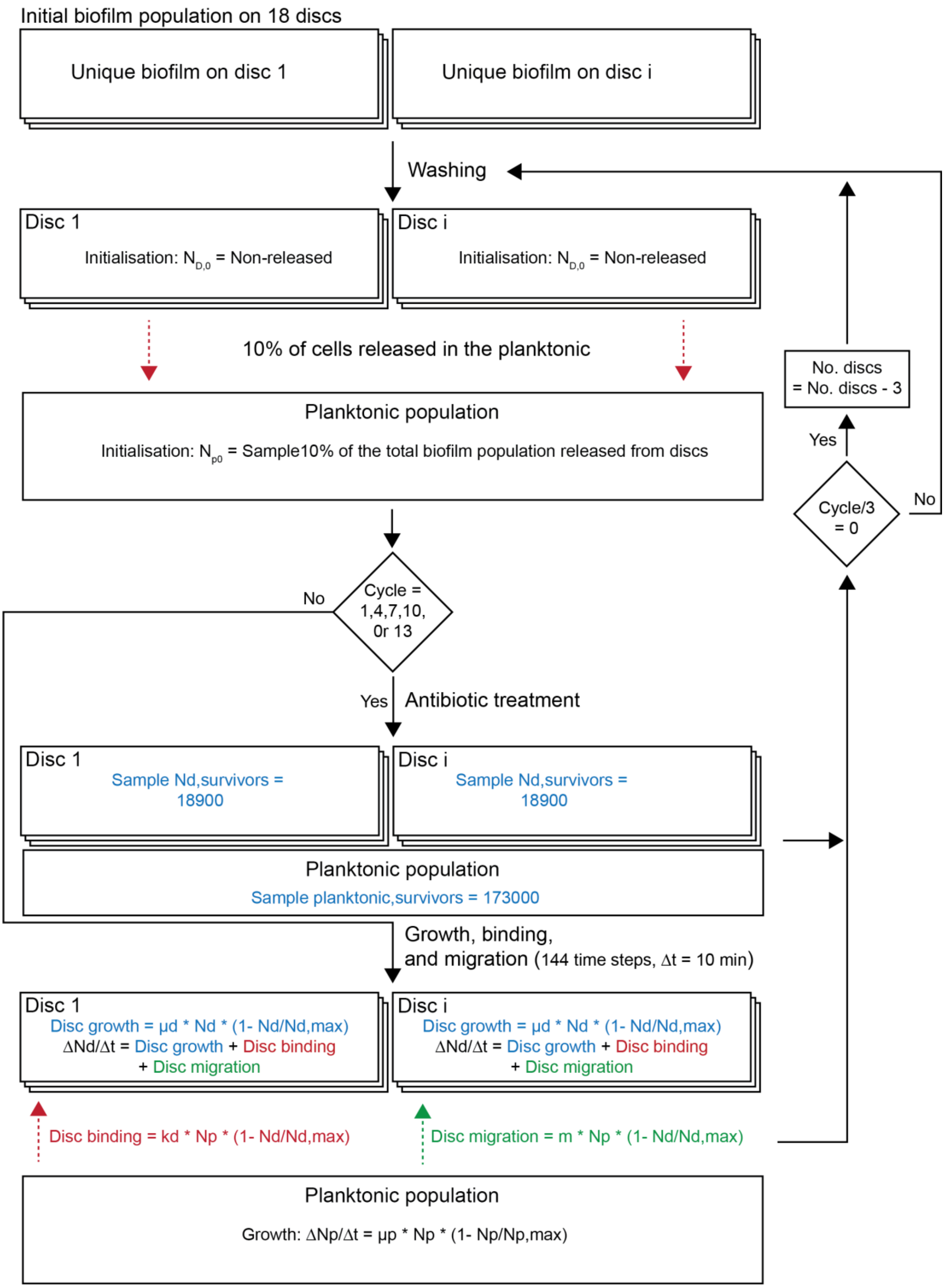


**Supplementary Figure 26:** Algorithm for population temporal evolution by binding, growth, migration, and antibiotic treatment.

### Supplementary Table 1| Estimated parameters for the binding-growth model

| | Symbol | Measurements | Mean Value | Source | Note |
|---|---|---|---|---|---|
| **Inoculum size** | $N_{po}$ | 4.30E+07 | 4.30E+07 | Determined experimentally (This Study) | CFUs/mL of the inoculum used |
| **Planktonic carrying capacity** | $N_{p,max}$ | 9.50E+09<br>1.05E+10 | 1.00E+10 | Determined experimentally (This Study) | CFUs/mL after 24 hour recovery |
| **Disc carrying capacity** | $N_{d,\ max}$ | 2.90E+06<br>1.90E+06<br>2.35E+06<br>2.85E+06<br>5.95E+06<br>3.05E+06 | 3.17E+06 | Determined experimentally (This Study) | CFUs/mL after 24 hour recovery |
| **Planktnic doubling time (min)** | $\mu_p$ | 33.08<br>35.42<br>34.21<br>35<br>34.21<br>32.02 | 33.99 | Determined experimentally (This Study) | Doubling time from exponential phase of growth curves |
| **Planktonic lag (min)** | | 60 | | Determined experimentally (This Study) | |
| **Biofilm doibling time (min)** | $\mu_d$ | 64.04893525<br>65.44130341<br>54.73272648<br>64.04893525<br>81.35945829<br>103.8034468<br>66.89555459<br>65.44130341<br>81.35945829<br>94.07187364<br>45.6106054<br>77.18717838<br>64.04893525<br>60.20599913<br>59.02548935<br>68.41590811<br>69.73106944 | 69.73106944 | Determined experimentally (This Study) | Experiment described below |

**Supplementary Table 2:** Sample-wise read counts (counts highlighted in red had very low reads and were excluded from analysis):

| S. No. | Sample | #Reads | #Barcodes | Pass/Fail | S. No. | Sample | #Reads | #Barcodes | Pass/Fail | S. No. | Sample | #Reads | #Barcodes | Pass/Fail | S. No. | Sample | #Reads | #Barcodes | Pass/Fail |
|---|---|---|---|---|---|---|---|---|---|---|---|---|---|---|---|---|---|---|---|
| 0 | 3 | 582321 | 88551 | Pass | 46 | 142 | 890960 | 58218 | Pass | 92 | 55 | 884149 | 45655 | Pass | 138 | 24 | 518369 | 70589 | Pass |
| 1 | 8 | 1085161 | 86746 | Pass | 47 | 146 | 860590 | 54935 | Pass | 93 | 91 | 651306 | 43767 | Pass | 139 | 36 | 590713 | 63884 | Pass |
| 2 | 28 | 487484 | 72334 | Pass | 48 | 161 | 1617401 | 49673 | Pass | 94 | 127 | 954313 | 37141 | Pass | 140 | 40 | 1846962 | 61594 | Pass |
| 3 | 32 | 150332 | 55946 | Fail | 49 | 165 | 12704506 | 43588 | Pass | 95 | 155 | 369335 | 17075 | Pass | 141 | 60 | 1211536 | 57758 | Pass |
| 4 | 42 | 2149843 | 76717 | Pass | 50 | 169 | 6977496 | 41691 | Pass | 96 | 179 | 922196 | 11281 | Pass | 142 | 68 | 2588747 | 55062 | Pass |
| 5 | 64 | 5285391 | 78316 | Pass | 51 | 15 | 169592 | 17797 | Fail | 97 | 20 | 105549 | 13998 | Fail | 143 | 76 | 1868939 | 49039 | Pass |
| 6 | 70 | 1442274 | 73043 | Pass | 52 | 51 | 220796 | 13390 | Pass | 98 | 56 | 327398 | 36419 | Pass | 144 | 96 | 1855974 | 45810 | Pass |
| 7 | 78 | 1131941 | 67269 | Pass | 53 | 87 | 166 | 15 | Fail | 99 | 92 | 990170 | 50734 | Pass | 145 | 104 | 786764 | 33814 | Pass |
| 8 | 100 | 895007 | 60482 | Pass | 54 | 123 | 891097 | 37647 | Pass | 100 | 128 | 2141166 | 37899 | Pass | 146 | 112 | 232712 | 22083 | Pass |
| 9 | 108 | 694722 | 49373 | Pass | 55 | 151 | 1102116 | 29658 | Pass | 101 | 156 | 2770905 | 23429 | Pass | 147 | 132 | 173138 | 14426 | Fail |
| 10 | 116 | 443535 | 40782 | Pass | 56 | 175 | 1721480 | 41450 | Pass | 102 | 180 | 758534 | 20542 | Pass | 148 | 5 | 757294 | 67836 | Pass |
| 11 | 134 | 1927481 | 32577 | Pass | 57 | 16 | 155159 | 18105 | Fail | 103 | 11 | 234958 | 64981 | Pass | 149 | 25 | 651707 | 69680 | Pass |
| 12 | 140 | 711147 | 29337 | Pass | 58 | 52 | 3661481 | 44399 | Pass | 104 | 31 | 530263 | 78563 | Pass | 150 | 37 | 590026 | 65757 | Pass |
| 13 | 144 | 536761 | 29298 | Pass | 59 | 88 | 660342 | 50622 | Pass | 105 | 35 | 762112 | 78114 | Pass | 151 | 44 | 3792447 | 63695 | Pass |
| 14 | 160 | 3213310 | 30711 | Pass | 60 | 124 | 1056870 | 39649 | Pass | 106 | 47 | 1777346 | 81099 | Pass | 152 | 61 | 1068760 | 58218 | Pass |
| 15 | 164 | 5678578 | 28958 | Pass | 61 | 152 | 2627303 | 36730 | Pass | 107 | 67 | 1926368 | 90179 | Pass | 153 | 72 | 3925933 | 55499 | Pass |
| 16 | 168 | 6675930 | 26892 | Pass | 62 | 176 | 1269515 | 30829 | Pass | 108 | 75 | 1276954 | 82554 | Pass | 154 | 80 | 1576118 | 48313 | Pass |
| 17 | 12 | 550972 | 21968 | Pass | 63 | 17 | 247703 | 21953 | Pass | 109 | 83 | 1813026 | 77602 | Pass | 155 | 97 | 24843 | 13189 | Fail |
| 18 | 48 | 363985 | 32790 | Pass | 64 | 53 | 661800 | 36357 | Pass | 110 | 103 | 1418535 | 82573 | Pass | 156 | 105 | 1214836 | 28392 | Pass |
| 19 | 84 | 1026089 | 38976 | Pass | 65 | 89 | 218853 | 39733 | Pass | 111 | 111 | 175224 | 46998 | Fail | 157 | 113 | 1094632 | 28551 | Pass |
| 20 | 120 | 222746 | 36391 | Pass | 66 | 125 | 896943 | 50863 | Pass | 112 | 119 | 271377 | 49900 | Pass | 158 | 136 | 957600 | 17816 | Pass |
| 21 | 148 | 930398 | 32030 | Pass | 67 | 153 | 2574176 | 41477 | Pass | 113 | 135 | 525854 | 58608 | Pass | 159 | 6 | 1063202 | 79987 | Pass |
| 22 | 172 | 845764 | 23164 | Pass | 68 | 177 | 1682090 | 41101 | Pass | 114 | 143 | 2923860 | 29972 | Pass | 160 | 26 | 2472061 | 69423 | Pass |
| 23 | 13 | 201846 | 18835 | Pass | 69 | 10 | 1619820 | 80716 | Pass | 115 | 147 | 2010144 | 29867 | Pass | 161 | 38 | 1973076 | 66430 | Pass |
| 24 | 49 | 2075174 | 49451 | Pass | 70 | 30 | 244251 | 70776 | Pass | 116 | 163 | 3147622 | 29088 | Pass | 162 | 41 | 1892385 | 65226 | Pass |
| 25 | 85 | 3197976 | 53962 | Pass | 71 | 34 | 916704 | 78158 | Pass | 117 | 167 | 3184096 | 23892 | Pass | 163 | 62 | 359837 | 23608 | Pass |
| 26 | 121 | 1477290 | 40603 | Pass | 72 | 43 | 2165480 | 79884 | Pass | 118 | 171 | 5641874 | 22625 | Pass | 164 | 69 | 2988535 | 21801 | Pass |
| 27 | 149 | 1534484 | 30596 | Pass | 73 | 66 | 1848382 | 91473 | Pass | 119 | 21 | 350283 | 28942 | Pass | 165 | 77 | 753724 | 20328 | Pass |
| 28 | 173 | 3771225 | 30099 | Pass | 74 | 71 | 1202877 | 82025 | Pass | 120 | 57 | 895676 | 39227 | Pass | 166 | 98 | 377780 | 16665 | Pass |
| 29 | 14 | 91586 | 16559 | Fail | 75 | 79 | 1092906 | 78351 | Pass | 121 | 93 | 1164411 | 37304 | Pass | 167 | 106 | 1222929 | 17860 | Pass |
| 30 | 50 | 723048 | 73169 | Pass | 76 | 102 | 532113 | 80927 | Pass | 122 | 129 | 988007 | 32577 | Pass | 168 | 114 | 1084323 | 21043 | Pass |
| 31 | 86 | 3449688 | 46281 | Pass | 77 | 110 | 1479656 | 56471 | Pass | 123 | 157 | 2198804 | 27385 | Pass | 169 | 133 | 2474335 | 21915 | Pass |
| 32 | 122 | 2710079 | 51996 | Pass | 78 | 118 | 816622 | 56279 | Pass | 124 | 181 | 1657793 | 20628 | Pass | 170 | 7 | 379096 | 68040 | Pass |
| 33 | 150 | 1380492 | 37854 | Pass | 79 | 139 | 492787 | 45951 | Pass | 125 | 22 | 146521 | 18046 | Fail | 171 | 27 | 740846 | 63237 | Pass |
| 34 | 174 | 714895 | 26933 | Pass | 80 | 141 | 683681 | 19195 | Pass | 126 | 58 | 146721 | 27879 | Fail | 172 | 39 | 2558494 | 60770 | Pass |
| 35 | 9 | 239604 | 73092 | Pass | 81 | 145 | 753068 | 21462 | Pass | 127 | 94 | 497028 | 31431 | Pass | 173 | 45 | 6445195 | 59879 | Pass |
| 36 | 29 | 1666940 | 77876 | Pass | 82 | 162 | 4665365 | 21935 | Pass | 128 | 130 | 109360 | 28306 | Fail | 174 | 63 | 760810 | 35218 | Pass |
| 37 | 33 | 1283696 | 75543 | Pass | 83 | 166 | 1572633 | 23362 | Pass | 129 | 158 | 383849 | 20536 | Pass | 175 | 73 | 2997825 | 32393 | Pass |
| 38 | 46 | 2828171 | 79885 | Pass | 84 | 170 | 5972827 | 20980 | Pass | 130 | 182 | 4337168 | 17802 | Pass | 176 | 81 | 809351 | 31408 | Pass |
| 39 | 65 | 3447708 | 76722 | Pass | 85 | 18 | 426214 | 27549 | Pass | 131 | 23 | 611665 | 29329 | Pass | 177 | 99 | 356721 | 28157 | Pass |
| 40 | 74 | 946386 | 72819 | Pass | 86 | 54 | 5259111 | 54003 | Pass | 132 | 59 | 1881602 | 70698 | Pass | 178 | 107 | 1514283 | 18133 | Pass |
| 41 | 82 | 2613435 | 70973 | Pass | 87 | 90 | 1302395 | 64436 | Pass | 133 | 95 | 4507827 | 74238 | Pass | 179 | 115 | 472067 | 19477 | Pass |
| 42 | 101 | 724121 | 69613 | Pass | 88 | 126 | 5252790 | 46307 | Pass | 134 | 131 | 1694153 | 53584 | Pass | 180 | 137 | 608582 | 11124 | Pass |
| 43 | 109 | 1095513 | 68417 | Pass | 89 | 154 | 5002756 | 25924 | Pass | 135 | 159 | 2608110 | 29304 | Pass | | | | | |
| 44 | 117 | 593969 | 66145 | | 90 | 178 | 1502576 | 22375 | Pass | 136 | 183 | 1915608 | 21303 | Pass | | | | | |
| 45 | 138 | 273654 | 57180 | | 91 | 19 | 49342 | 11582 | Fail | 137 | 4 | 1926542 | 81211 | Pass | | | | | |

**Supplementary Table 3 (Mutated lineages highlighted in blue):** Mutations identified in variants with unique barcodes at the end of the biofilm experiment. Sample names beginning with CXX were recovered from the biofilm-associated planktonic population and ones with names DXX & EXX were recovered from two independent discs from the same well. All barcodes were recovered from replicate A.

| | Sample | Source | Barcode | Position | Mutation | Annotation | Type | Gene | Description |
|---|---|---|---|---|---|---|---|---|---|
| 0 | C1 | Planktonic | TTCACCTATTTTGTATCATTTATTTAGTAT | No Mutation | | | | No Mutation | |
| 2 | C11 | Planktonic | TTGCGCGATAATAAATTCTATATTGTCAAT | 3 529 671 | C→T | A145V (GCA→GTA) | Non-synonymous | *crp* → | cAMP-activated global transcriptional regulator CRP |
| 20 | C15 | Planktonic | TTCCGGAATAGAACATTCGTTATTACCCAT | 4 667 186 | T→A | V77D (GTC→GAC) | Non-synonymous | *fimH* → | type 1 fimbria D-mannose specific adhesin FimH |
| 44 | C17 | Planktonic | TTCTTAAATGTTGTATCTAATATAGATAAT | No Mutation | | | | No Mutation | |
| 45 | C19 | Planktonic | TTGTCGCATGCTCGATTAGCAATTTCAAAT | 4 667 326 | T→C | W124R (TGG→CGG) | Non-synonymous | *fimH* → | type 1 fimbria D-mannose specific adhesin FimH |
| 49 | C22 | Planktonic | TTTTATTATTCGGTATGGGAAATAAAGCAT | 3 744 301 | C→T | H295Y (CAC→TAC) | Non-synonymous | *bcsG* → | cellulose biosynthesis protein BcsG |
| 56 | C24 | Planktonic | TTGACTGATTTCCAATTTTTCATATGCGAT | 4 667 042 | Δ36 bp | coding (86-121/903 nt) | Indel | *fimH* → | type 1 fimbria D-mannose specific adhesin FimH |
| 59 | C25 | Planktonic | TTCTAGCATAGTAGATCTTCGATTCGTCAT | 4 667 042 | Δ36 bp | coding (86-121/903 nt) | Indel | *fimH* → | type 1 fimbria D-mannose specific adhesin FimH |
| 62 | C26 | Planktonic | TTTTTCGATTTTTAATCTTTAATTGCCAAT | | | | | No Mutation | |
| 63 | C29 | Planktonic | TTATGCGATATTTTATATCTAATTTCGAAT | 4 037 507 | G→T | A401S (GCG→TCG) | Non-synonymous | *cyaA* → | class I adenylate cyclase |
| 65 | C3 | Planktonic | TTAGCATATCTGTGATGTAGTATAACATAT | 4 667 234 | Δ15 bp | coding (278-292/903 nt) | Indel | *fimH* → | type 1 fimbria D-mannose specific adhesin FimH |
| 69 | C30 | Planktonic | TTTTTGAATCATTTATTTCATATATTCGAT | 2 159 109 | A→C | intergenic (-130/-188) | Intergenic | *udk* ← / → *dgcE* | uridine kinase/diguanylate cyclase |
| 71 | C33 | Planktonic | TTTGCTCATAGAACATTTCTCATCACTCAT | 2 336 068 | C→A | P555H (CCC→CAC) | Non-synonymous | *atoS* → | two-component system sensor histidine kinase AtoS |
| 71 | C33 | Planktonic | TTTGCTCATAGAACATTTCTCATCACTCAT | 4 667 042 | Δ36 bp | coding (86-121/903 nt) | Indel | *fimH* → | type 1 fimbria D-mannose specific adhesin FimH |
| 77 | C50 | Planktonic | TTAAATTATATGGTATCAGTGATAGAATAT | 3 743 656 | T→C | W80R (TGG→CGG) | Non-synonymous | *bcsG* → | cellulose biosynthesis protein BcsG |
| 79 | C54 | Planktonic | TTGTTTAATCCAAAATGCGGTATACATGAT | 2 381 019 | A→T | T320S (ACC→TCC) | Non-synonymous | *arnB* → | UDP-4-amino-4-deoxy-L-arabinose aminotransferase |
| 79 | C54 | Planktonic | TTGTTTAATCCAAAATGCGGTATACATGAT | 4 667 243 | T→A | V96E (GTA→GAA) | Non-synonymous | *fimH* → | type 1 fimbria D-mannose specific adhesin FimH |
| 89 | C79 | Planktonic | TTCTTAGATTATCGATCAGTGATCGCGGAT | 3 529 671 | C→T | A145V (GCA→GTA) | Non-synonymous | *crp* → | cAMP-activated global transcriptional regulator CRP |
| 89 | C79 | Planktonic | TTCTTAGATTATCGATCAGTGATCGCGGAT | 4 630 527 | C→G | P202A (CCA→GCA) | Non-synonymous | *LF82_RS22855* → | DUF898 family protein |
| 90 | C80 | Planktonic | TTATGGGATTGCCGATTGCGGATTGCGGAT | No Mutation | | | | No Mutation | |
| 92 | C84 | Planktonic | TTGCTAGATGACCAATTAAATATTGGGTAT | No Mutation | | | | No Mutation | |
| 93 | C85 | Planktonic | TTTAGCGATACTGAATTGCTCATAGCGGAT | 1 764 564 | Δ460 bp | | Deletion | *[ydiV]* | *[ydiV]* |
| 2 | D10 | Disc A | TTCACGGATATCATATACTAGATGTGAGAT | 344 541 | G→T | intergenic (-116/+970) | Intergenic | *ipbA* ← / ← *betA* | tyrosine-type DNA invertase IpbA/choline dehydrogenase |
| 12 | D13 | Disc A | __GATACATTCTTAATTTTAAATCGTTCAT | \ | | | | No Mutation | |
| 13 | D14 | Disc A | T-------TGTCTAATCATGAATACCACAT | 3 117 907 | G→A | R605H (CGC→CAC) | Non-synonymous | *LF82_RS15440* → | capsular polysaccharide biosynthesis protein |

| | | | | | | | | | |
|---|---|---|---|---|---|---|---|---|---|
| 14 | D15 | Disc A | TTATTTTATTTTGCATAGCTCATGCCTTAT | \ | | | | No Mutation | |
| 15 | D16 | Disc A | TTTGGGTATGTTTTATGACGAATGTGGTAT | \ | | | | No Mutation | |
| 16 | D17 | Disc A | TTATCCAATTGTGCATTTTTGATACGGCAT | \ | | | | No Mutation | |
| 17 | D18 | Disc A | TTTTATGATGTCCGATACTCAATCCGCTAT | \ | | | | No Mutation | |
| 18 | D19 | Disc A | TCCATCTATTAAGGATGAATTATAGGTTAT | 3 586 452 | (C)6→5 | coding (279/2085 nt) | | *malQ* ← | 4-alpha-glucanotransferase |
| 19 | D21 | Disc A | TTTGAATATTTTTTATAGCATATGCTCTAT | 2 484 106 | T→A | S814R (AGT→AGA) | Non-synonymous | *evgS* → | acid-sensing system histidine kinase EvgS |
| 19 | D21 | Disc A | TTTGAATATTTTTTATAGCATATGCTCTAT | 4 356 136 | G→T | S380R (AGC→AGA) | Non-synonymous | *LF82_RS21480* ← | ATP-binding protein |
| 20 | D22 | Disc A | TTCTTCGATGATTGATGTACTATTTTGTAT | \ | | | | No Mutation | |
| 22 | D25 | Disc A | TTGCGCCATCTGGCTACGCGCCTGGTGCGT | 1 276 622 | C→A | Q263K (CAA→AAA) | Non-synonymous | *prfA* → | peptide chain release factor 1 |
| 22 | D25 | Disc A | TTGCGCCATCTGGCTACGCGCCTGGTGCGT | 4 064 694 | G→C | A109A (GCG→GCC) | Synonymous | *LF82_RS20105* → | SIS domain-containing protein |
| 24 | D29 | Disc A | TTCGACGATGTCATATGTGACATCTTGTAT | 4 667 326 | T→C | W124R (TGG→CGG) | Non-synonymous | *fimH* → | type 1 fimbria D-mannose specific adhesin FimH |
| 25 | D3 | Disc A | TTCTTTAATTTTCAATGGTGCATCACATAT | \ | | | | No Mutation | |
| 26 | D30 | Disc A | TTTTGGCATTCGGAATCTTTTATTATCGAT | \ | | | | No Mutation | |
| 28 | D32 | Disc A | TTTTAGCATACTGAATCTTTCATATAATAT | 4 047 949 | G→A | R160R (CGG→CGA) | Synonymous | *corA* → | magnesium/cobalt transporter CorA |
| 29 | D33 | Disc A | TTTTCTGATAGTCTATGATACATTCACCAT | \ | | | | No Mutation | |
| 34 | D38 | Disc A | TTAGTTGATCGTCCATCTTTTATATGGAAT | 2 756 447 | C→T | A238V (GCC→GTC) | Non-synonymous | *LF82_RS13695* → | terminase ATPase subunit family protein |
| 36 | D4 | Disc A | TTTGTAGATAGTTTATAATCTATCTTATAT | \ | | | | No Mutation | |
| 49 | D44 | Disc A | TTTTGGTATTTCCCATTACAAATAATTCAT | 4 480 917 | Δ1 bp | coding (957/1473 nt) | Indel | *LF82_RS22090* → | TIGR03752 family integrating conjugative element protein |
| 50 | D46 | Disc A | TTGAGTGATTACGTATGTCGAATATACAAT | \ | | | | No Mutation | |
| 52 | D48 | Disc A | TTATTTTATAGGCTATGTTACATCACAAAT | \ | | | | No Mutation | |
| 54 | D50 | Disc A | TTGGCTGATAGACAATGGCGGATCTAATAT | \ | | | | No Mutation | |
| 55 | D51 | Disc A | TTGTTTCATATCCAATTAGCTATACCGAAT | \ | | | | No Mutation | |
| 56 | D53 | Disc A | TTCCTTTATTCTCGATTGCATATCCTTAAT | 1 276 526 | G→A | G231R (GGG→AGG) | Non-synonymous | *prfA* → | peptide chain release factor 1 |
| 58 | D56 | Disc A | TTTTAAGATCGGGGATGGGGTATAACGTAT | \ | | | | No Mutation | |
| 59 | D57 | Disc A | TTGACTAATGGATGATTGGTTATTGTTGAT | \ | | | | No Mutation | |
| 61 | D59 | Disc A | _TCGTATATGCCCTATGTTGAATAGGCCAT | 1 327 677 | G→C | T33R (ACG→AGG) | Non-synonymous | *yciB* ← | septation protein A |
| 62 | D6 | Disc A | TTCGTTGATCTTCAATAGAGAATACAAAAT | \ | | | | No Mutation | |
| 67 | D66 | Disc A | TTTTCATATAGTCTATAGTTCATTTAATAT | 3 529 670 | G→A | A145T (GCA→ACA) | Non-synonymous | *crp* → | cAMP-activated global transcriptional regulator CRP |
| 68 | D67 | Disc A | TTCTGGAATATGCCATTTTTCATTCAATAT | \ | | | | No Mutation | |
| 69 | D68 | Disc A | TTGATATATCTTCGATGCTGAATTTGAGAT | \ | | | | No Mutation | |
| 70 | D69 | Disc A | TTTAGGGATGGAATATACGAGATTAGCAAT | \ | | | | No Mutation | |
| 75 | D77 | Disc A | TTTATAAATACTTCATCGGACATATCGCAT | \ | | | | No Mutation | |
| 77 | D79 | Disc A | TTAGATCATTCGGTATCAATAATAACACAT | \ | | | | No Mutation | |
| 83 | D86 | Disc A | TTGATGGATATTCGATAACTCATTCGACAT | \ | | | | No Mutation | |
| 84 | D87 | Disc A | TTTGGCGATTTTACATCGCTTATAGTACAT | \ | | | | No Mutation | |
| 85 | D89 | Disc A | TTCTTTCATTTGTAATGGCCAATTTTGTAT | 1 092 035 | G→A | P38S (CCC→TCC) | Non-synonymous | *mdtG* ← | multidrug efflux MFS transporter MdtG |
| 88 | D91 | Disc A | TTTGTTTATCAACGATTAGCTATCCACTAT | \ | | | | No Mutation | |

| | | | | | | | | | |
|---|---|---|---|---|---|---|---|---|---|
| 90 | D93 | Disc A | TTTATTTATAGCCTATGGGGCATCTTACAT | \ | | | | No Mutation | |
| 3 | E10 | Disc B | TTTAGAGATGCTCAATATACTATTCGCCAT | \ | | | | No Mutation | |
| 4 | E11 | Disc B | TTTGTGCATGGAAAATCGGTCATCTTGGAT | \ | | | | No Mutation | |
| 5 | E12 | Disc B | TTCTGTGATGGAGAATCAATGATAACGGAT | \ | | | | No Mutation | |
| 22 | E17 | Disc B | TTTATGGATTGAGAATAATATATACCCTAT | 2 790 273 | T→G | L74R (CTT→CGT) | Non-synonymous | *proX* → | glycine betaine/L-proline ABC transporter substrate-binding protein ProX |
| 24 | E18 | Disc B | TTTTTCTATCTAACATTACCTATCTTAAAT | 2 209 539 | T→C | M1M (ATG→GTG) † | Synonymous | *yehB* ← | fimbrial biogenesis outer membrane usher protein |
| 27 | E20 | Disc B | TTAGGTTATTATGGATGAGGTATTGGTGAT | 1 276 388 | C→A | Q185K (CAG→AAG) | Non-synonymous | *prfA* → | peptide chain release factor 1 |
| 30 | E22 | Disc B | TTACTATATCAGGCATCCACCATACTGTAT | 4 265 998 | T→C | V1237A (GTA→GCA) | Non-synonymous | *rpoC* → | DNA-directed RNA polymerase subunit beta' |
| 34 | E25 | Disc B | TTTCGATATTGGCAATGCGGAATGCGGTAT | \ | | | | No Mutation | |
| 35 | E26 | Disc B | TTCGCGCATTGACAATTTTTTATCCGGTAT | \ | | | | No Mutation | |
| 36 | E27 | Disc B | TTGTTATATGTATTATTCGCAATACGTGAT | \ | | | | No Mutation | |
| 39 | E31 | Disc B | TTCGAGAATATTTCATGCAATATTTAGGAT | \ | | | | No Mutation | |
| 40 | E32 | Disc B | TTTTGCTATTCAGCATGGGCTATCCACAAT | \ | | | | No Mutation | |
| 41 | E33 | Disc B | TTAACTAATCTGAAATAAGGTATTTACTAT | \ | | | | No Mutation | |
| 42 | E34 | Disc B | TTATCATATTGGACATTTTAAATTTGAAAT | \ | | | | No Mutation | |
| 44 | E36 | Disc B | TTGCGGCATCCAATATTTGTTATTCAAGAT | \ | | | | No Mutation | |
| 47 | E4 | Disc B | TTCATTGATATCCAATCGGTGATTCTCTAT | | | | | No Mutation | |
| 48 | E40 | Disc B | TTTGCTGATTTGTTATTCATGATTTTAGAT | \ | | | | No Mutation | |
| 51 | E44 | Disc B | TTTGTATATGTACAATGGCTTATTTACAAT | 4 030 927 | C→A | E79* (GAG→TAG) | Nonsense | *aslA* ← | arylsulfatase AslA |
| 53 | E47 | Disc B | TTGATAGATCATGCATAAGGAATCACAGAT | \ | | | | No Mutation | |
| 54 | E48 | Disc B | TTTCTTTATTGGCTATGTTCTATGTTCGAT | \ | | | | No Mutation | |
| 57 | E51 | Disc B | TTCTTCGATGGATGATGGAAAATTCGTAAT | \ | | | | No Mutation | |
| 58 | E53 | Disc B | TTTTCTTATTCCACATTCACAATCCCTAAT | \ | | | | No Mutation | |
| 60 | E55 | Disc B | TTGGGGGATGCATAATGCGTAATCATCGAT | 320 745 | C→A | Q80K (CAA→AAA) | Non-synonymous | *LF82_RS01505* → | LysR family transcriptional regulator |
| 62 | E58 | Disc B | TTGCTTCATACTATATATGGAATCATTCAT | \ | | | | No Mutation | |
| 63 | E59 | Disc B | TTGTTTTATAGTTAATCTTTCATCTTGCAT | \ | | | | No Mutation | |
| 64 | E6 | Disc B | TTAGAGGATTCTTGATATTTTATAAGTTAT | 3 590 832 | A→T | E359D (GAA→GAT) | Non-synonymous | *malT* → | HTH-type transcriptional regulator MalT |
| 65 | E60 | Disc B | TTGCACTATCAACAATTTTGTATTGCCGAT | \ | | | | No Mutation | |
| 66 | E61 | Disc B | TTGAGCAATTGTCAATCAGTCATTATGGAT | \ | | | | No Mutation | |
| 67 | E62 | Disc B | TTACAGCATCTCCGATCGAGAATTGTCGAT | \ | | | | No Mutation | |
| 68 | E63 | Disc B | TTAGTCAATCTCCTATCATTTATTGTCTAT | \ | | | | No Mutation | |
| 69 | E64 | Disc B | TTACGTAATTTACTATAAGCCATAATCAAT | \ | | | | No Mutation | |
| 71 | E66 | Disc B | TTGAAAAATCAACCATCCGAAATTGTATAT | \ | | | | No Mutation | |
| 73 | E70 | Disc B | TTCTCTGATTTCTCATTTGGAATAGCATAT | \ | | | | No Mutation | |
| 74 | E71 | Disc B | TTGGGCCATCTGACATGGTACATTCTCAAT | | | | | No Mutation | |
| 75 | E73 | Disc B | TTAGATTATAGACGATCAATTATCCCGGAT | \ | | | | No Mutation | |
| 77 | E76 | Disc B | TTAACAGATAGAACATATGTAATTAGTGAT | \ | | | | No Mutation | |
| 78 | E77 | Disc B | TTGATTAATGTTGGATTGAGGATAATGAAT | \ | | | | No Mutation | |

| | | | | | | | | | |
|---|---|---|---|---|---|---|---|---|---|
| 79 | E78 | Disc B | TTGTTGCATGGCAGATGCGCTATCGCATAT | \ | | | | No Mutation | |
| 82 | E81 | Disc B | TTGTATTATCTCTGATGGATTATCTCTGAT | \ | | | | No Mutation | |
| 83 | E82 | Disc B | TTGGTTTATGGTGAATTGGTGATTCCTGAT | 1 270 106 | A→T | F499I (TTC→ATC) | Non-synonymous | *dauA* ← | C4-dicarboxylic acid transporter DauA |
| 83 | E82 | Disc B | TTGGTTTATGGTGAATTGGTGATTCCTGAT | 3 101 733 | G→A | Q72* (CAG→TAG) | Nonsense | *yggL* ← | YggL family protein |
| 84 | E84 | Disc B | TTGCCCGATTGCCAATGGGTTATCCTAAAT | 4 549 466 | (T)9→8 | intergenic (+47/-80) | Intergenic | *rlmB* → / → *yjfI* | 23S rRNA (guanosine(2251)-2'-O)-methyltransferase RlmB/DUF2170 family protein |
| 84 | E84 | Disc B | TTGCCCGATTGCCAATGGGTTATCCTAAAT | 344 541 | G→T | intergenic (-116/+970) | Intergenic | *ipbA* ← / ← *betA* | tyrosine-type DNA invertase IpbA/choline dehydrogenase |
| 85 | E85 | Disc B | TTATGCAATCAACTATTCGTTATTTCGTAT | fimE/fimA | | | | FimE/FimA | |
| 85 | E85 | Disc B | TTATGCAATCAACTATTCGTTATTTCGTAT | 1 764 467 | Δ1 bp | coding (411/714 nt) | Indel | *ydiV* ← | anti-FlhDC factor |
| 88 | E88 | Disc B | TTGGATTATAAGGGATAGTCAATTCCTAAT | 971 981 | Δ1 bp | intergenic (-124/+232) | Intergenic | *ompA* ← / ← *sulA* | porin OmpA/cell division inhibitor SulA |
| 88 | E88 | Disc B | TTGGATTATAAGGGATAGTCAATTCCTAAT | 4 522 841 | C→T | A46V (GCG→GTG) | Non-synonymous | *epmA* → | elongation factor P--(R)-beta-lysine ligase |
| 90 | E9 | Disc B | TTTGGGAATTGTAAATAACTCATAGTTGAT | \ | | | | No Mutation | |
| 91 | E91 | Disc B | TTGTTGCATTGTATATGTCACATTGTCCAT | 2 258 728 | C→A | M237I (ATG→ATT) | Non-synonymous | *mglB* ← | galactose/glucose ABC transporter substrate-binding protein MglB |
| 92 | E92 | Disc B | TTCGCTCATTAGCCATACGGTATTCAGCAT | | | | | No Mutation | |
| 93 | E93 | Disc B | TTGGTGCATATGGTATAACTCATGAACTAT | | | | | No Mutation | |
| 95 | E96 | Disc B | TTACCTGATGCTTGATAAGGAATCAGTCAT | | | | | No Mutation | |

**Supplementaty Table 4:** Mutations in variants with unique barcodes at the end of the well-mixed shaken-flask experiment

| | Sample | Barcode | Position | Mutation | Annotation | Type | Gene | Description |
|---|---|---|---|---|---|---|---|---|
| 1 | A10 | TTGTCAAATCCGTGATCTTCAATTCGCAAT | 4,771,187 | A→G | F79L (TTC→CTC) | Non-synonymous | *arcA* ← | two-component system response regulator ArcA |
| 2 | A11 | TTCCAGGATGTATTATTTAAAATGCTCAAT | 4,771,375 | C→T | R16H (CGC→CAC) | Non-synonymous | *arcA* ← | two-component system response regulator ArcA |
| 3 | A12 | TTTACTAATGGTACATGACAAATAGGCGAT | 1,867,647 | Δ4,581 bp | | Deletion | *[manY]–[yebO]* | *[manY], manZ, yobD, mntP, rlmA, cspE, yobF, [yebO]* |
| 3 | A12 | TTTACTAATGGTACATGACAAATAGGCGAT | 874,147 | Δ18,750 bp | | Deletion | *yebQ–[ptrB]* | **19 genes yebQ, tnpA, htpX, prc, proQ, msrC, yebS, yebT, rsmF, yebV, yebW, pphA, yebY, copD, yobA, holE, yobB, exoX, [ptrB]** |
| 4 | A13 | TTGGTGGATTATTTATTGGGTATATGACAT | 815,113 | C→T | A69T (GCG→ACG) | Non-synonymous | *ybiY* ← | glycyl-radical enzyme activating protein |
| 4 | A13 | TTGGTGGATTATTTATTGGGTATATGACAT | 4,771,190 | T→A | M78L (ATG→TTG) | Non-synonymous | *arcA* ← | two-component system response regulator ArcA |
| 5 | A14 | TTATAATATACGTTATCTAATATTCATTAT | 3,409,281 | A→C | intergenic (-57/+52) | Non-synonymous | *nanT* ← / ← *nanA* | sialic acid transporter NanT/N-acetylneuraminate lyase |
| 5 | A14 | TTATAATATACGTTATCTAATATTCATTAT | 4,771,354 | A→G | F23S (TTC→TCC) | Non-synonymous | *arcA* ← | two-component system response regulator ArcA |
| 6 | A15 | TTATTGTATTTGGGATCTTTTATAAAAAAT | 1,981,272 | Δ1 bp | coding (690/1374 nt) | Indel | *fliI* → | flagellar protein export ATPase FliI |
| 6 | A15 | TTATTGTATTTGGGATCTTTTATAAAAAAT | 4,771,072 | A→T | L117Q (CTG→CAG) | Non-synonymous | *arcA* ← | two-component system response regulator ArcA |
| 7 | A16 | TTAAGGTATTTATTATTATCGATTGGATAT | 4,771,225 | G→A | A66V (GCG→GTG) | Non-synonymous | *arcA* ← | two-component system response regulator ArcA |
| 8 | A17 | TTACGATATTCATCATTTAGGATTGTTCAT | 4,771,213 | C→T | R70H (CGC→CAC) | Non-synonymous | *arcA* ← | two-component system response regulator ArcA |
| 9 | A18 | TTTTCATATAGAACATCCCGTATTATTTAT | 2,009,133 | A→G | A408A (GCA→GCG) | Non-synonymous | *LF82_RS10260* → | tyrosine-type recombinase/integrase |
| 9 | A18 | TTTTCATATAGAACATCCCGTATTATTTAT | 4,771,225 | G→A | A66V (GCG→GTG) | Non-synonymous | *arcA* ← | two-component system response regulator ArcA |
| 10 | A19 | TTCTCCAATGCACTATCGGAAATTATATAT | 4,771,273 | A→T | L50Q (CTG→CAG) | Non-synonymous | *arcA* ← | two-component system response regulator ArcA |
| 11 | A2 | TTCTGTTATGGCACATAGAAAATCACGCAT | 897,869 | Δ24 bp | coding (2460-2483/4053 nt) | Indel | *ftsK* → | DNA translocase FtsK |
| 11 | A2 | TTCTGTTATGGCACATAGAAAATCACGCAT | 4,771,305 | C→T | M39I (ATG→ATA) | Non-synonymous | *arcA* ← | two-component system response regulator ArcA |

| | | | | | | | | |
|---|---|---|---|---|---|---|---|---|
| 12 | A20 | TTATATTATCGGGTATTTCAAATATTATAT | 4,771,083 | A→T | R113R (CGT→CGA) | Non-synonymous | *arcA* ← | two-component system response regulator ArcA |
| 12 | A20 | TTATATTATCGGGTATTTCAAATATTATAT | 4,771,093 | A→G | L110P (CTG→CCG) | Non-synonymous | *arcA* ← | two-component system response regulator ArcA |
| 13 | A21 | TTATTTGATTAATGATCTGCCATAATGAAT | 4,771,256 | T→A | N56Y (AAT→TAT) | Non-synonymous | *arcA* ← | two-component system response regulator ArcA |
| 14 | A22 | TTTTTATATAATAAATCGCGAATACCATAT | 1,261,260 | C→T | R91H (CGC→CAC) | Non-synonymous | *treA* ← | alpha,alpha-trehalase |
| 14 | A22 | TTTTTATATAATAAATCGCGAATACCATAT | 4,771,306 | A→T | M39K (ATG→AAG) | Non-synonymous | *arcA* ← | two-component system response regulator ArcA |
| 15 | A23 | TTAATTGATGCTGCATCCACAATTTCGTAT | 4,771,090 | G→T | T111N (ACT→AAT) | Non-synonymous | *arcA* ← | two-component system response regulator ArcA |
| 15 | A23 | TTAATTGATGCTGCATCCACAATTTCGTAT | 4,771,324 | G→T | A33E (GCG→GAG) | Non-synonymous | *arcA* ← | two-component system response regulator ArcA |
| 16 | A24 | TTATCTAATCAAATATTGGCAATGGTTCAT | 2,143,383 | A→C | V15G (GTC→GGC) | Non-synonymous | *fcl* ← | GDP-L-fucose synthase |
| 16 | A24 | TTATCTAATCAAATATTGGCAATGGTTCAT | 4,771,096 | T→A | E109V (GAA→GTA) | Non-synonymous | *arcA* ← | two-component system response regulator ArcA |
| 17 | A26 | TTAATTGATGCTGCATCCACAATTTCGTAT | 4,771,225 | G→A | A66V (GCG→GTG) | Non-synonymous | *arcA* ← | two-component system response regulator ArcA |
| 17 | A26 | TTAATTGATGCTGCATCCACAATTTCGTAT | 726,543 | T→A | Y156N (TAC→AAC) | Non-synonymous | *cydB* → | cytochrome d ubiquinol oxidase subunit II |
| 18 | A27 | TTAATGTATTAGCTATGATTTATTTATAAT | 4,771,187 | A→G | F79L (TTC→CTC) | Non-synonymous | *arcA* ← | two-component system response regulator ArcA |
| 19 | A28 | TTGTGGGATTATTAATTGCAGATTCGAAAT | 4,771,225 | G→A | A66V (GCG→GTG) | Non-synonymous | *arcA* ← | two-component system response regulator ArcA |
| 20 | A29 | TTATAAAATACGTCATTGCGCATTCAGTAT | 4,740,700 | C→G | A105G (GCG→GGG) | Non-synonymous | *prfC* → | peptide chain release factor 3 |
| 20 | A29 | TTATAAAATACGTCATTGCGCATTCAGTAT | 4,771,079 | G→A | R115C (CGC→TGC) | Non-synonymous | *arcA* ← | two-component system response regulator ArcA |
| 21 | A33 | TTATAGCATCCTATATTTATTATATGCCAT | 4,771,120 | A→C | I101S (ATC→AGC) | Non-synonymous | *arcA* ← | two-component system response regulator ArcA |
| 21 | A33 | TTATAGCATCCTATATTTATTATATGCCAT | 269,034 | C→T | R90C (CGT→TGT) | Non-synonymous | *LF82_RS01240* → | type VI secretion system tip protein VgrG |
| 21 | A33 | TTATAGCATCCTATATTTATTATATGCCAT | 3,018,493 | G→T | L476L (CTG→CTT) | Non-synonymous | *ygfK* → | putative selenate reductase subunit YgfK |
| 22 | A35 | TTCTCTAATTATCTATTTCAGATGCCCAAT | 4,771,061 | T→G | T121P (ACC→CCC) | Non-synonymous | *arcA* ← | two-component system response regulator ArcA |
| 23 | A38 | TTGGGTAATGAACGATACGGTATTAATTAT | 4,771,315 | C→G | G36A (GGC→GCC) | Non-synonymous | *arcA* ← | two-component system response regulator ArcA |
| 24 | A39 | TTTCCTTATTTTGAATTCCAAATGGTGAAT | 4,771,250 | G→T | P58T (CCG→ACG) | Non-synonymous | *arcA* ← | two-component system response regulator ArcA |

| | | | | | | | | |
|---|---|---|---|---|---|---|---|---|
| 25 | A4 | TTGTTTTATCTTAAATTCATTATTCCTTAT | 4,771,461 | T→C | intergenic (-40/-56) | Intergenic | *arcA* ← / → *LF82_RS23530* | two-component system response regulator ArcA/protein YjjY |
| 26 | A40 | TTGGCGCATTGGCAATGATATATAAAGGAT | 1,950,885 | A→T | intergenic (+15/+23) | Intergenic | *ftnA* → / ← *yecH* | non-heme ferritin/YecH family protein |
| 26 | A40 | TTGGCGCATTGGCAATGATATATAAAGGAT | 4,771,223 | G→A | R67C (CGT→TGT) | Non-synonymous | *arcA* ← | two-component system response regulator ArcA |
| 27 | A42 | TTGCCGTATTGAATATTCGTGATGATATAT | 2,100,149 | A→C | Q54H (CAA→CAC) | Non-synonymous | *pduT* → | propanediol utilization microcompartment protein PduT |
| 27 | A42 | TTGCCGTATTGAATATTCGTGATGATATAT | 125,368 | T→A | intergenic (+126/-33) | Intergenic | *LF82_RS00585* → / → *LF82_RS00590* | bacteriocin immunity protein/HNH endonuclease signature motif containing protein |
| 27 | A42 | TTGCCGTATTGAATATTCGTGATGATATAT | 4,771,242 | C→G | K60N (AAG→AAC) | Non-synonymous | *arcA* ← | two-component system response regulator ArcA |
| 28 | A43 | TTTAACAATCAATTATCTCGTATCTATTAT | 4,771,234 | A→C | L63R (CTT→CGT) | Non-synonymous | *arcA* ← | two-component system response regulator ArcA |
| 29 | A44 | TTAATACATTCGGTATTTTGCATGTCTGAT | 1,620,924 | C→T | L340F (CTT→TTT) | Non-synonymous | *LF82_RS08150* → | FRG domain-containing protein |
| 29 | A44 | TTAATACATTCGGTATTTTGCATGTCTGAT | 4,771,343 | C→G | G27R (GGC→CGC) | Non-synonymous | *arcA* ← | two-component system response regulator ArcA |
| 30 | A46 | TTAACGTATTGCCCATTTGCAATTATTTAT | 726,543 | T→A | Y156N (TAC→AAC) | Non-synonymous | *cydB* → | cytochrome d ubiquinol oxidase subunit II |
| 31 | A47 | TTTATTTATACGAAATCCAGTATATTTTAT | 4,771,165 | T→C | E86G (GAA→GGA) | Non-synonymous | *arcA* ← | two-component system response regulator ArcA |
| 32 | A48 | TTCAAATATTTGGTATAGAGGATATCTAAT | 4,771,305 | C→T | M39I (ATG→ATA) | Non-synonymous | *arcA* ← | two-component system response regulator ArcA |
| 33 | A5 | TTGGCTGATATTATATTAAGTATATCCAAT | 3,396,493 | A→T | Y71N (TAC→AAC) | Non-synonymous | *arcB* ← | aerobic respiration two-component sensor histidine kinase ArcB |
| 34 | A6 | TTTATTAATGATTTATATTTAATTGTTCAT | | | | Non-synonymous | | |
| 35 | A7 | TTGCATTATAATTTATTCCTTATATTACAT | 4,771,225 | G→A | A66V (GCG→GTG) | Non-synonymous | *arcA* ← | two-component system response regulator ArcA |
| 36 | A8 | TTTATAGATAAGTGATAAATCATCTTTAAT | 4,771,325 | C→G | A33P (GCG→CCG) | Non-synonymous | *arcA* ← | two-component system response regulator ArcA |

**Supplementaty Table 5:** Mutations in variants with unique barcodes at the end of the biofilm evolution with antibiotic treatment

| | Sample | Source | Barcode | position | mutation | annotation | Type | gene | description |
|---|---|---|---|---|---|---|---|---|---|
| 0 | F1 | Planktonic AMK | TTGTTGTA-CTTTTATGGCATATAATTTAT | 400 594 | G→T | E137* (GAA→TAA) | Non-synonymous | *LF82_RS01845* → | peptide antibiotic transporter SbmA |
| 1 | F10 | Planktonic AMK | TTACTACATATTATATGCATCATCGCCTAT | 401 231 | T→G | L349R (CTC→CGC) | Non-synonymous | *LF82_RS01845* → | peptide antibiotic transporter SbmA |
| 45 | F11 | Planktonic AMK | TTTGCGAATTTTTAATAGTTAATTTGCCAT | 401 262 | (T)5→4 | coding (1077/1221 nt) | Indel | *LF82_RS01845* → | peptide antibiotic transporter SbmA |
| 46 | F12 | Planktonic AMK | TTGAAGGATTTTACATTGAGAATCTACCAT | 400 850 | Δ1 bp | coding (665/1221 nt) | Indel | *LF82_RS01845* → | peptide antibiotic transporter SbmA |
| 47 | F16 | Planktonic AMK | TTGGCAGATGGTTCATGAGTCATTGTCTAT | 401 051 | Δ1 bp | coding (866/1221 nt) | Indel | *LF82_RS01845* → | peptide antibiotic transporter SbmA |
| 49 | F17 | Planktonic AMK | TTCCTGCATCGTGTATTTATTATCCCCTAT | 400 687 | A→C | T168P (ACG→CCG) | Non-synonymous | *LF82_RS01845* → | peptide antibiotic transporter SbmA |
| 50 | F18 | Planktonic AMK | TTTGGCTATGCGTTATCCGCTATTTGTCAT | 400 202 | (T)6→5 | coding (17/1221 nt) | Indel | *LF82_RS01845* → | peptide antibiotic transporter SbmA |
| 51 | F19 | Planktonic AMK | TTTGCTCATTAAGGATGGCAAATCTAGAAT | 401 028 | Δ9 bp | coding (843-851/1221 nt) | Indel | *LF82_RS01845* → | peptide antibiotic transporter SbmA |
| 52 | F2 | Planktonic AMK | TTTCAGGATCTTGTATTTGCTATTGACTAT | 400 686 | Δ1 bp | coding (501/1221 nt) | Indel | *LF82_RS01845* → | peptide antibiotic transporter SbmA |
| 54 | F21 | Planktonic AMK | TTCATCCATCTCGTATATCTCATAGGAAAT | 401 167 | C→A | Q328K (CAG→AAG) | Non-synonymous | *LF82_RS01845* → | peptide antibiotic transporter SbmA |
| 54 | F21 | Planktonic AMK | TTCATCCATCTCGTATATCTCATAGGAAAT | 4 667 271 | Δ63 bp | coding (315-377/903 nt) | Indel | *LF82_RS23020* → | type 1 fimbriae D-mannose specific adhesin FimH |
| 64 | F28 | Planktonic AMK | TTCTAGCATATGAGATCGCCCATCTAGCAT | 305 558 | C→A | intergenic (-268/-325) | Intergenic | *LF82_RS01410* ← / → *LF82_RS25025* | vacuolating autotransporter toxin Vat/IS1 family transposase |
| 64 | F28 | Planktonic AMK | TTCTAGCATATGAGATCGCCCATCTAGCAT | 401 099 | T→A | V305E (GTA→GAA) | Non-synonymous | *LF82_RS01845* → | peptide antibiotic transporter SbmA |

| | | | | | | | | | |
|---|---|---|---|---|---|---|---|---|---|
| 64 | F28 | Planktonic AMK | TTCTAGCATATGAGATCGCCCATCTAGCAT | 2 313 573 | G→A | L17L (CTG→TTG) | Synonymous | *napB* ← | nitrate reductase cytochrome c-type subunit |
| 64 | F28 | Planktonic AMK | TTCTAGCATATGAGATCGCCCATCTAGCAT | 4 366 465 | Δ43 bp | coding (74-116/219 nt) | Indel | *LF82_RS24890* ← | hypothetical protein |
| 65 | F31 | Planktonic AMK | TTGAATTATGTCACATCGT---TTATAGAT | 13 472 | G→T | K528N (AAG→AAT) | Non-synonymous | *LF82_RS00065* → | molecular chaperone DnaK |
| 65 | F31 | Planktonic AMK | TTGAATTATGTCACATCGT---TTATAGAT | 13 481 | G→T | E531D (GAG→GAT) | Non-synonymous | *LF82_RS00065* → | molecular chaperone DnaK |
| 65 | F31 | Planktonic AMK | TTGAATTATGTCACATCGT---TTATAGAT | 398 619 | Δ1,659 bp | | Indel | *[LF82_RS01830]–[LF82_RS01845]* | *[LF82_RS01830], LF82_RS01835, LF82_RS01840, [LF82_RS01845]* |
| 65 | F31 | Planktonic AMK | TTGAATTATGTCACATCGT---TTATAGAT | 2 174 195 | G→A | D174N (GAT→AAT) | Non-synonymous | *LF82_RS10950* → | multidrug efflux RND transporter permease subunit MdtC |
| 66 | F33 | Planktonic AMK | TTAGTTCATTTGCGATAATCTATCTATTAT | 400 990 | G→A | E269K (GAG→AAG) | Non-synonymous | *LF82_RS01845* → | peptide antibiotic transporter SbmA |
| 67 | F34 | Planktonic AMK | TTTTTCTATAACTTATAGGTCATTCATAAT | 401 099 | T→A | V305E (GTA→GAA) | Non-synonymous | *LF82_RS01845* → | peptide antibiotic transporter SbmA |
| 68 | F36 | Planktonic AMK | TTCTGGTATTTCAGATTAGGTATAATAGAT | 401 287 | Δ1 bp | coding (1102/1221 nt) | Indel | *LF82_RS01845* → | peptide antibiotic transporter SbmA |
| 69 | F38 | Planktonic AMK | TTCCAGTATCGTCAATGATTAATAATTTAT | 400 202 | (T)6→5 | coding (17/1221 nt) | Indel | *LF82_RS01845* → | peptide antibiotic transporter SbmA |
| 70 | F39 | Planktonic AMK | TTTTTAAATATTTGATGATGCATACAAGAT | 401 221 | Δ1 bp | coding (1036/1221 nt) | Indel | *LF82_RS01845* → | peptide antibiotic transporter SbmA |
| 71 | F40 | Planktonic AMK | TTTTAGCATAGGGGATGCCCCATTATCAAT | 401 099 | T→A | V305E (GTA→GAA) | Non-synonymous | *LF82_RS01845* → | peptide antibiotic transporter SbmA |
| 72 | F43 | Planktonic AMK | TTGGGGAATAAAGTATCGCTCATTCCTCAT | 400 344 | G→A | W53* (TGG→TGA) | Non-synonymous | *LF82_RS01845* → | peptide antibiotic transporter SbmA |
| 72 | F43 | Planktonic AMK | TTGGGGAATAAAGTATCGCTCATTCCTCAT | 3 450 670 | C→T | R13C (CGC→TGC) | Non-synonymous | *LF82_RS17065* → | DUF997 family protein |
| 73 | F48 | Planktonic AMK | TTCGAGTATATTCAATTTTGGATGAAATAT | 400 202 | (T)6→5 | coding (17/1221 nt) | Indel | *LF82_RS01845* → | peptide antibiotic transporter SbmA |
| 74 | F5 | Planktonic AMK | TTGGGTGATGTGATATCGTACATTGCTTAT | 401 086 | Δ229 bp | coding (901-1129/1221 nt) | Indel | *LF82_RS01845* → | peptide antibiotic transporter SbmA |

| | | | | | | | | | |
|---|---|---|---|---|---|---|---|---|---|
| 75 | F50 | Planktonic AMK | TTTCGCGATTTGTAATCAGGTATTTGAGAT | 401 099 | T→A | V305E (GTA→GAA) | Non-synonymous | *LF82_RS01845* → | peptide antibiotic transporter SbmA |
| 75 | F50 | Planktonic AMK | TTTCGCGATTTGTAATCAGGTATTTGAGAT | 3 410 983 | C→T | E53K (GAA→AAA) | Non-synonymous | *LF82_RS16865* ← | transcriptional regulator NanR |
| 76 | F56 | Planktonic AMK | TTGTTGTATGTCATATGTCGTATCACTTAT | 401 099 | Δ1 bp | coding (914/1221 nt) | Indel | *LF82_RS01845* → | peptide antibiotic transporter SbmA |
| 79 | F62 | Planktonic AMK | TTGCACGATATGTTATCCTTGATTCCTCAT | 401 152 | Δ1 bp | coding (967/1221 nt) | Indel | *LF82_RS01845* → | peptide antibiotic transporter SbmA |
| 81 | F66 | Planktonic AMK | TTCCGCAATCGTTGATTATTGATAATTGAT | 249 813 | G→A | T62M (ACG→ATG) | Non-synonymous | *LF82_RS01155* ← | type VI secretion system-associated protein TagO |
| 81 | F66 | Planktonic AMK | TTCCGCAATCGTTGATTATTGATAATTGAT | 274 220 | C→T | pseudogene (364/765 nt) | Synonymous | *LF82_RS25020* → | hypothetical protein |
| 81 | F66 | Planktonic AMK | TTCCGCAATCGTTGATTATTGATAATTGAT | 401 291 | T→C | L369P (CTG→CCG) | Non-synonymous | *LF82_RS01845* → | peptide antibiotic transporter SbmA |
| 81 | F66 | Planktonic AMK | TTCCGCAATCGTTGATTATTGATAATTGAT | 1 169 032 | A→G | S91S (AGT→AGC) | Synonymous | *LF82_RS05730* ← | lysogenization regulator HflD |
| 81 | F66 | Planktonic AMK | TTCCGCAATCGTTGATTATTGATAATTGAT | 1 424 509 | G→A | A233V (GCG→GTG) | Non-synonymous | *LF82_RS07185* ← | porin OmpN |
| 81 | F66 | Planktonic AMK | TTCCGCAATCGTTGATTATTGATAATTGAT | 2 219 954 | C→T | P351L (CCT→CTT) | Non-synonymous | *LF82_RS11145* → | DUF4132 domain-containing protein |
| 81 | F66 | Planktonic AMK | TTCCGCAATCGTTGATTATTGATAATTGAT | 2 523 887 | T→C | Y19C (TAC→TGC) | Non-synonymous | *LF82_RS12570* ← | hypothetical protein |
| 81 | F66 | Planktonic AMK | TTCCGCAATCGTTGATTATTGATAATTGAT | 2 598 494 | A→G | S746S (AGT→AGC) | Synonymous | *LF82_RS12930* ← | cyclic-guanylate-specific phosphodiesterase PdeF |
| 81 | F66 | Planktonic AMK | TTCCGCAATCGTTGATTATTGATAATTGAT | 3 457 799 | A→G | K226R (AAG→AGG) | Non-synonymous | *LF82_RS17105* → | multidrug efflux RND transporter periplasmic adaptor subunit AcrE |
| 81 | F66 | Planktonic AMK | TTCCGCAATCGTTGATTATTGATAATTGAT | 3 504 579 | G→A | M1I (ATG→ATA) † | Non-synonymous | *LF82_RS17435* → | type II secretion system protein GspH |
| 81 | F66 | Planktonic AMK | TTCCGCAATCGTTGATTATTGATAATTGAT | 4 331 717 | T→C | T566A (ACC→GCC) | Non-synonymous | *LF82_RS21380* ← | glycerol-3-phosphate 1-O-acyltransferase |
| 81 | F66 | Planktonic AMK | TTCCGCAATCGTTGATTATTGATAATTGAT | 4 537 127 | (TGGCGC)3→2 | coding (215-220/1848 nt) | Indel | *LF82_RS22395* → | DNA mismatch repair endonuclease MutL |

| | | | | | | | | | |
|---|---|---|---|---|---|---|---|---|---|
| 81 | F66 | Planktonic AMK | TTCCGCAATCGTTGATTATTGATAATTGAT | 4 682 727 | A→G | T355A (ACT→GCT) | Non-synonymous | *LF82_RS23095* → | carnitine transporter CniT |
| 83 | F68 | Planktonic AMK | TTTTGTGATGGGGTATTGTGGATTGACTAT | 401 287 | Δ1 bp | coding (1102/1221 nt) | Indel | *LF82_RS01845* → | peptide antibiotic transporter SbmA |
| 84 | F69 | Planktonic AMK | TTGTATCATTTTTCATGTTAGATATAGAAT | 401 099 | T→A | V305E (GTA→GAA) | Non-synonymous | *LF82_RS01845* → | peptide antibiotic transporter SbmA |
| 84 | F69 | Planktonic AMK | TTGTATCATTTTTCATGTTAGATATAGAAT | 1 364 786 | G→A | R61C (CGC→TGC) | Non-synonymous | *LF82_RS06870* ← | CMD domain-containing protein |
| 85 | F72 | Planktonic AMK | TTCGCATATCAGAAATTCATCATATTAGAT | 401 099 | T→A | V305E (GTA→GAA) | Non-synonymous | *LF82_RS01845* → | peptide antibiotic transporter SbmA |
| 86 | F75 | Planktonic AMK | TTTTGGAATCGAGGATGTATTATCGGATAT | 401 167 | C→T | Q328* (CAG→TAG) | Non-synonymous | *LF82_RS01845* → | peptide antibiotic transporter SbmA |
| 87 | F77 | Planktonic AMK | TTCCACCATTCGAGATTGCTTATACTATAT | 400 687 | A→C | T168P (ACG→CCG) | Non-synonymous | *LF82_RS01845* → | peptide antibiotic transporter SbmA |
| 88 | F78 | Planktonic AMK | TTACGCAATCTTTAATTGGAGATCTTTCAT | 400 386 | C→A | C67* (TGC→TGA) | Non-synonymous | *LF82_RS01845* → | peptide antibiotic transporter SbmA |
| 89 | F79 | Planktonic AMK | TTGGTGCATTGGGGATATGGCATGGCGTAT | 400 687 | Δ1 bp | coding (502/1221 nt) | Indel | *LF82_RS01845* → | peptide antibiotic transporter SbmA |
| 90 | F87 | Planktonic AMK | TTAGTCGATTCGAGATAGGGTATCAGGTAT | 401 099 | Δ1 bp | coding (914/1221 nt) | Indel | *LF82_RS01845* → | peptide antibiotic transporter SbmA |
| 92 | F9 | Planktonic AMK | TTGTGTTATGAGGTATTGTCTATTCGCCAT | 400 883 | Δ1 bp | coding (698/1221 nt) | Indel | *LF82_RS01845* → | peptide antibiotic transporter SbmA |
| 93 | F90 | Planktonic AMK | TTCTAGTATTTGTGATCTCCCATTTAATAT | 344 541 | G→T | pseudogene (11/185 nt) | Synonymous | *LF82_RS23720* → | hypothetical protein |
| 93 | F90 | Planktonic AMK | TTCTAGTATTTGTGATCTCCCATTTAATAT | 400 202 | (T)6→5 | coding (17/1221 nt) | Indel | *LF82_RS01845* → | peptide antibiotic transporter SbmA |
| 95 | F96 | Planktonic AMK | TTAGCTTATTTTTCATGCCCTATGTTGAAT | 401 289 | C→A | Y368* (TAC→TAA) | Non-synonymous | *LF82_RS01845* → | peptide antibiotic transporter SbmA |
| 35 | G10 | Disc A AMK | TTTACTGATTGGCCATTAAAGATGGATCAT | 401 099 | T→A | V305E (GTA→GAA) | Non-synonymous | *LF82_RS01845* → | peptide antibiotic transporter SbmA |
| 37 | G11 | Disc A AMK | TTTTTGTATACGTTATCGTCTATGTGGCAT | 401 077 | Δ147 bp | coding (892-1038/1221 nt) | Indel | *LF82_RS01845* → | peptide antibiotic transporter SbmA |

| | | | | | | | | | |
|---|---|---|---|---|---|---|---|---|---|
| 38 | G12 | Disc A AMK | TTTAATCATCGCGCATTCGTTATTTTCAAT | 400 517 | G→A | W111* (TGG→TAG) | Non-synonymous | *LF82_RS01845* → | peptide antibiotic transporter SbmA |
| 39 | G13 | Disc A AMK | TTGTTTAATGTTTTATACATAATGCTCCAT | 400 856 | Δ135 bp | coding (671-805/1221 nt) | Indel | *LF82_RS01845* → | peptide antibiotic transporter SbmA |
| 52 | G15 | Disc A AMK | TTTTACAATATTGCATAACCGATACGATAT | 399 947 | Δ351 bp | | Indel | *LF82_RS01840–[LF82_RS01845]* | *LF82_RS01840, [LF82_RS01845]* |
| 52 | G15 | Disc A AMK | TTTTACAATATTGCATAACCGATACGATAT | 1 527 430 | C→A | L368L (CTG→CTT) | Synonymous | *LF82_RS07650* ← | glutamate:gamma-aminobutyrate antiporter |
| 52 | G15 | Disc A AMK | TTTTACAATATTGCATAACCGATACGATAT | 3 976 064 | A→C | L353R (CTC→CGC) | Non-synonymous | *LF82_RS19670* ← | ATPase RavA |
| 53 | G16 | Disc A AMK | TTCTTTGATGTGCTATAGCATATCACAAAT | 400 748 | C→T | A188V (GCA→GTA) | Non-synonymous | *LF82_RS01845* → | peptide antibiotic transporter SbmA |
| 55 | G17 | Disc A AMK | TTTTCGTATTTAAAATCGAACATCTATCAT | 400 717 | T→A | W178R (TGG→AGG) | Non-synonymous | *LF82_RS01845* → | peptide antibiotic transporter SbmA |
| 55 | G17 | Disc A AMK | TTTTCGTATTTAAAATCGAACATCTATCAT | 3 423 891 | A→T | L68Q (CTG→CAG) | Non-synonymous | *LF82_RS16935* ← | p-hydroxybenzoic acid efflux pump subunit AaeA |
| 59 | G20 | Disc A AMK | TTGCCTTATAGTATATGCGCAATGATGCAT | 401 099 | T→A | V305E (GTA→GAA) | Non-synonymous | *LF82_RS01845* → | peptide antibiotic transporter SbmA |
| 60 | G23 | Disc A AMK | TTGCCCCATTGTCTATCGCGCATTTCTTAT | 400 687 | Δ1 bp | coding (502/1221 nt) | Indel | *LF82_RS01845* → | peptide antibiotic transporter SbmA |
| 61 | G27 | Disc A AMK | TTCTACAATTTTAAATTTTTAATCACCCAT | 401 063 | (CG)4→3 | coding (878-879/1221 nt) | Indel | *LF82_RS01845* → | peptide antibiotic transporter SbmA |
| 62 | G28 | Disc A AMK | TTTCGCTATCATGTATTTTCCATGTCTAAT | 400 757 | Δ1 bp | coding (572/1221 nt) | Indel | *LF82_RS01845* → | peptide antibiotic transporter SbmA |
| 65 | G3 | Disc A AMK | TTTTGGCATAATCTATCTACAATCCGCTAT | 401 303 | G→A | W373* (TGG→TAG) | Non-synonymous | *LF82_RS01845* → | peptide antibiotic transporter SbmA |
| 67 | G33 | Disc A AMK | TTCCGTAATAGTCCATACTTAATCTAAGAT | 400 687 | Δ1 bp | coding (502/1221 nt) | Indel | *LF82_RS01845* → | peptide antibiotic transporter SbmA |
| 68 | G34 | Disc A AMK | TTTTTATATCGAGTATTGTTTATAGGCTAT | 400 202 | (T)6→5 | coding (17/1221 nt) | Indel | *LF82_RS01845* → | peptide antibiotic transporter SbmA |
| 69 | G35 | Disc A AMK | TTAAATTATAACTTATTCTCCATAGAATAT | 400 687 | Δ1 bp | coding (502/1221 nt) | Indel | *LF82_RS01845* → | peptide antibiotic transporter SbmA |

| | | | | | | | | | |
|---|---|---|---|---|---|---|---|---|---|
| 70 | G38 | Disc A AMK | TTCGAACATTTAGTATAGAGGATACAGGAT | 400 630 | Δ1 bp | coding (445/1221 nt) | Indel | *LF82_RS01845* → | peptide antibiotic transporter SbmA |
| 70 | G38 | Disc A AMK | TTCGAACATTTAGTATAGAGGATACAGGAT | 3 395 641 | G→A | P355S (CCG→TCG) | Non-synonymous | *LF82_RS16815* ← | aerobic respiration two-component sensor histidine kinase ArcB |
| 72 | G40 | Disc A AMK | TTACGCTATAACTCATTTTAAATCGCTGAT | 400 487 | (T)5→4 | coding (302/1221 nt) | Indel | *LF82_RS01845* → | peptide antibiotic transporter SbmA |
| 72 | G40 | Disc A AMK | TTACGCTATAACTCATTTTAAATCGCTGAT | 3 209 715 | T→A | K92N (AAA→AAT) | Non-synonymous | *LF82_RS15870* ← | OB fold stress tolerance protein YgiW |
| 74 | G42 | Disc A AMK | TTTCACGATGGATAATAGCGGATAAACGAT | 400 242 | G→A | W19* (TGG→TGA) | Non-synonymous | *LF82_RS01845* → | peptide antibiotic transporter SbmA |
| 74 | G42 | Disc A AMK | TTTCACGATGGATAATAGCGGATAAACGAT | 3 209 715 | T→A | K92N (AAA→AAT) | Non-synonymous | *LF82_RS15870* ← | OB fold stress tolerance protein YgiW |
| 76 | G44 | Disc A AMK | TTCGGTTATACGGGATCACATATTTTAAAT | 401 007 | Δ1 bp | coding (822/1221 nt) | Indel | *LF82_RS01845* → | peptide antibiotic transporter SbmA |
| 80 | G52 | Disc A AMK | TTGTCTCATTCAGTATCCATTATGATTTAT | 400 349 | T→A | L55* (TTG→TAG) | Non-synonymous | *LF82_RS01845* → | peptide antibiotic transporter SbmA |
| 81 | G54 | Disc A AMK | TTCCATCATCGTCTATTTTTAATGTATAAT | 400 504 | G→C | A107P (GCC→CCC) | Non-synonymous | *LF82_RS01845* → | peptide antibiotic transporter SbmA |
| 86 | G55 | Disc A AMK | TTTGCTCATCGTTAATAACTCATAATCTAT | 400 858 | Δ1 bp | coding (673/1221 nt) | Indel | *LF82_RS01845* → | peptide antibiotic transporter SbmA |
| 87 | G57 | Disc A AMK | TTACAGGATCCCTAATAAGTCATCTTTAAT | 401 099 | T→A | V305E (GTA→GAA) | Non-synonymous | *LF82_RS01845* → | peptide antibiotic transporter SbmA |
| 88 | G58 | Disc A AMK | TTCACTGATCTAACATCAAATATCCACTAT | 400 374 | C→A | Y63* (TAC→TAA) | Non-synonymous | *LF82_RS01845* → | peptide antibiotic transporter SbmA |
| 88 | G58 | Disc A AMK | TTCACTGATCTAACATCAAATATCCACTAT | 400 380 | T→G | I65M (ATT→ATG) | Non-synonymous | *LF82_RS01845* → | peptide antibiotic transporter SbmA |
| 90 | G68 | Disc A AMK | TTAATAGATTAGCGATGATTGATTCCTTAT | 400 747 | G→A | A188T (GCA→ACA) | Non-synonymous | *LF82_RS01845* → | peptide antibiotic transporter SbmA |
| 91 | G70 | Disc A AMK | TTCCACTATGCTGGATTGCACATTCCTGAT | 401 099 | Δ1 bp | coding (914/1221 nt) | Indel | *LF82_RS01845* → | peptide antibiotic transporter SbmA |

| | | | | | | | | | |
|---|---|---|---|---|---|---|---|---|---|
| 93 | G82 | Disc A AMK | TTATGTCATAACCTATCTGCAAT-TCTTAT | 400 410 | G→A | W75* (TGG→TGA) | Non-synonymous | *LF82_RS01845* → | peptide antibiotic transporter SbmA |
| 94 | G87 | Disc A AMK | TTAATGTATTAGCGATCTGCCATCAGATAT | 400 487 | (T)5→4 | coding (302/1221 nt) | Indel | *LF82_RS01845* → | peptide antibiotic transporter SbmA |
| 94 | G87 | Disc A AMK | TTAATGTATTAGCGATCTGCCATCAGATAT | 3 427 277 | C→T | V1223V (GTG→GTA) | Synonymous | *LF82_RS16955* ← | AsmA2 domain-containing protein |
| 39 | H10 | Disc B AMK | TAACATCCCGAATTAGTTATAGATTAT | 400 475 | Δ1 bp | coding (290/1221 nt) | Indel | *LF82_RS01845* → | peptide antibiotic transporter SbmA |
| 40 | H11 | Disc B AMK | TTCGTTCATCCGACATGAGATATTCGGTAT | 400 748 | C→T | A188V (GCA→GTA) | Non-synonymous | *LF82_RS01845* → | peptide antibiotic transporter SbmA |
| 40 | H11 | Disc B AMK | TTCGTTCATCCGACATGAGATATTCGGTAT | 1 544 473 | T→C | intergenic (-115/+238) | Intergenic | *LF82_RS07715* ← / ← *LF82_RS07720* | fimbrial chaperone protein FimC/Fml fimbriae subunit |
| 51 | H15 | Disc B AMK | TTTAGGTATGTTTTATCCCTAATAAGTAAT | 401 099 | T→A | V305E (GTA→GAA) | Non-synonymous | *LF82_RS01845* → | peptide antibiotic transporter SbmA |
| 52 | H17 | Disc B AMK | TTTATTCATACACCATGGTTTATTCCAAAT | 169 715 | T→A | R27R (CGT→CGA) | Synonymous | *LF82_RS00795* → | bifunctional glycosyl transferase/transpeptidase |
| 52 | H17 | Disc B AMK | TTTATTCATACACCATGGTTTATTCCAAAT | 400 748 | C→T | A188V (GCA→GTA) | Non-synonymous | *LF82_RS01845* → | peptide antibiotic transporter SbmA |
| 52 | H17 | Disc B AMK | TTTATTCATACACCATGGTTTATTCCAAAT | 3 569 118 | C→A | V372V (GTG→GTT) | Synonymous | *LF82_RS17780* ← | DUF4153 domain-containing protein |
| 53 | H18 | Disc B AMK | TTAACGCATACGTTATCCCAAATTTTTTAT | 401 099 | T→A | V305E (GTA→GAA) | Non-synonymous | *LF82_RS01845* → | peptide antibiotic transporter SbmA |
| 60 | H28 | Disc B AMK | TTGCATTATAAGATATGAGATATGTTGTAT | 401 221 | Δ1 bp | coding (1036/1221 nt) | Indel | *LF82_RS01845* → | peptide antibiotic transporter SbmA |
| 61 | H29 | Disc B AMK | TTACTCTATGTTTTATCGTGCATCCAGTAT | 400 521 | T→A | Y112* (TAT→TAA) | Non-synonymous | *LF82_RS01845* → | peptide antibiotic transporter SbmA |
| 63 | H31 | Disc B AMK | TTACGTCATAAAATATTGTGTATCCATTAT | 401 035 | Δ54 bp | coding (850-903/1221 nt) | Indel | *LF82_RS01845* → | peptide antibiotic transporter SbmA |
| 65 | H34 | Disc B AMK | ATTGCGGATGTTCCATCCAGAATACCCCAT | 400 757 | Δ1 bp | coding (572/1221 nt) | Indel | *LF82_RS01845* → | peptide antibiotic transporter SbmA |
| 65 | H34 | Disc B AMK | ATTGCGGATGTTCCATCCAGAATACCCCAT | 1 672 992 | A→G | V128A (GTA→GCA) | Non-synonymous | *LF82_RS08405* ← | beta-glucuronidase |

| | | | | | | | | | |
|---|---|---|---|---|---|---|---|---|---|
| 65 | H34 | Disc B AMK | ATTGCGGATGTTCCATCCAGAATACCCCAT | 4 187 164 | T→C | E259E (GAA→GAG) | Synonymous | *LF82_RS20735* ← | cell division protein FtsN |
| 67 | H38 | Disc B AMK | TTGGTGCATTAGCTATGGTACATCTCGCAT | 400 501 | +G | coding (316/1221 nt) | Indel | *LF82_RS01845* → | peptide antibiotic transporter SbmA |
| 71 | H43 | Disc B AMK | TTGCCGCATCGTATATGTCCAATTTTGCAT | 400 433 | G→A | W83* (TGG→TAG) | Non-synonymous | *LF82_RS01845* → | peptide antibiotic transporter SbmA |
| 72 | H44 | Disc B AMK | TTGATCAATTTCAAATTAAAAATCTATAAT | 401 153 | Δ1 bp | coding (968/1221 nt) | Indel | *LF82_RS01845* → | peptide antibiotic transporter SbmA |
| 77 | H58 | Disc B AMK | TTAACTTATGCTGCATCTTGAATCTGTAAT | 400 699 | G→T | E172* (GAA→TAA) | Non-synonymous | *LF82_RS01845* → | peptide antibiotic transporter SbmA |
| 78 | H6 | Disc B AMK | TTAACTTATGGGGAATGGAGGATTTATCAT | 401 168 | A→C | Q328P (CAG→CCG) | Non-synonymous | *LF82_RS01845* → | peptide antibiotic transporter SbmA |
| 79 | H60 | Disc B AMK | TTTACCTATGGTTTATTTGTAATCCTCAAT | 30 257 | G→A | G105E (GGG→GAG) | Non-synonymous | *LF82_RS00150* → | 4-hydroxy-tetrahydrodipicolinate reductase |
| 79 | H60 | Disc B AMK | TTTACCTATGGTTTATTTGTAATCCTCAAT | 400 757 | Δ1 bp | coding (572/1221 nt) | Indel | *LF82_RS01845* → | peptide antibiotic transporter SbmA |
| 82 | H66 | Disc B AMK | TTGCCTCATTTCCAATCGGGTATATAATAT | 400 262 | +T | coding (77/1221 nt) | Indel | *LF82_RS01845* → | peptide antibiotic transporter SbmA |
| 82 | H66 | Disc B AMK | TTGCCTCATTTCCAATCGGGTATATAATAT | 3 889 239 | A→C | L183R (CTG→CGG) | Non-synonymous | *LF82_RS19240* ← | lipoprotein NlpA |
| 83 | H69 | Disc B AMK | TTCCAAGATTAATGATATACTATTAGGAAT | 400 344 | G→A | W53* (TGG→TGA) | Non-synonymous | *LF82_RS01845* → | peptide antibiotic transporter SbmA |
| 84 | H72 | Disc B AMK | TTCTTGCATATTATATAATCTATTGTCAAT | 3 516 450 | A→C | I61M (ATT→ATG) | Non-synonymous | *LF82_RS17490* ← | elongation factor G |
| 87 | H82 | Disc B AMK | TTCATTCATCACTTATGCAAAATCTTCAAT | 401 138 | Δ1 bp | coding (953/1221 nt) | Indel | *LF82_RS01845* → | peptide antibiotic transporter SbmA |
| 88 | H83 | Disc B AMK | TTGTGAAATAGTGGATAAAAGATGAACAAT | 401,03 | A→G | E282G (GAG→GGG) | Non-synonymous | *LF82_RS01845* → | peptide antibiotic transporter SbmA |
| 89 | H88 | Disc B AMK | TTGCCCTATCAGTGATAATTTATTTTAAAT | 400 192 | A→T | K3* (AAG→TAG) | Non-synonymous | *LF82_RS01845* → | peptide antibiotic transporter SbmA |
| 89 | H88 | Disc B AMK | TTGCCCTATCAGTGATAATTTATTTTAAAT | 1 408 999 | A→T | V258E (GTA→GAA) | Non-synonymous | *LF82_RS07085* ← | low conductance mechanosensitive channel YnaI |

| | | | | | | | | | |
|---|---|---|---|---|---|---|---|---|---|
| 91 | H91 | Disc B AMK | TTAG----TTTGCAATTTGCAATTTCGTAT | 400 686 | Δ1 bp | coding (501/1221 nt) | Indel | *LF82_RS01845* → | peptide antibiotic transporter SbmA |
| 93 | H96 | Disc B AMK | TTTTTTGATCGCACATAGTTTATGGGTTAT | 401 287 | Δ1 bp | coding (1102/1221 nt) | Indel | *LF82_RS01845* → | peptide antibiotic transporter SbmA |

Supplementary figures, tables, methods, and notes
